# Functional characterization of a multi-cancer risk locus on chromosome band 2q33.1 near *CASP8*

**DOI:** 10.64898/2026.01.06.697866

**Authors:** Hyunkyung Kong, Jiyeon Choi, Tongwu Zhang, Cathrin Gräwe, Mai Xu, Rohit Thakur, Hayley Sowards, Rebecca C Hennessey, Andrew Vu, Jianxin Shi, D Timothy Bishop, Julia Newton-Bishop, Jeremie Nsengimana, Mark M. Iles, Maria Teresa Landi, Michiel Vermeulen, Matthew H. Law, Laufey T. Amundadottir, Melanoma Meta-Analysis Consortium, Kevin M Brown

**Affiliations:** Laboratory of Translational Genomics, Division of Cancer Epidemiology and Genetics, National Cancer Institute, Bethesda, MD, 20892, USA; Integrative Tumor Epidemiology Branch, Division of Cancer Epidemiology and Genetics, National Cancer Institute, Bethesda, MD, 20892, USA; Department of Molecular Biology, Radboud Institute for Molecular Life Sciences, Radboud University Nijmegen, Nijmegen, 6525 XZ, Netherlands; Division of Cancer Epidemiology and Genetics, National Cancer Institute, Bethesda, MD, 20892, USA; Leeds Institute of Medical Research, School of Medicine, University of Leeds, Leeds, UK; Leeds Institute for Data Analytics, University of Leeds, Leeds, UK; Population Health Sciences Institute, Faculty of Medical Sciences, Newcastle University and NIHR Biomedical Research Centre, Newcastle upon Tyne, UK; NIHR Leeds Biomedical Research Centre, Leeds Teaching Hospitals NHS Trust, Leeds, UK; Division of Molecular Genetics, Netherlands Cancer Institute, Amsterdam, Netherlands; Population Health Department, QIMR Berghofer Medical Research Institute, Herston, QLD, Australia; School of Biomedical Sciences, Faculty of Health, Queensland University of Technology, Brisbane, QLD, Australia; School of Biomedical Sciences, University of Queensland, Brisbane, QLD, Australia

**Keywords:** Melanoma, genome-wide association study, GWAS, expression quantitative trait locus, eQTL, fine-mapping, caspase 8, CASP8

## Abstract

Genome-wide association studies (GWAS) of melanoma have identified numerous susceptibility loci. However, causal genes and variants underlying risk have yet to be established for most. It is becoming apparent that many functional variants underlying complex traits act via *cis*-regulation that may be context-specific, dependent on availability of specific transcription factors/complexes in specific cell types and cell-states. To characterize a risk locus on chromosome band 2q33.1 associated with melanoma, breast cancer, and keratinocyte cancers, we integrated fine-mapping, cell-type specific expression quantitative trait locus (eQTL) analysis, a massively parallel reporter assay, individual luciferase assays, and SNP-based proteomics. Integrated analysis implicates the presence of multiple functional variants lying primarily within a promoter for *CASP8*. A haplotype containing rs3769823 appeared have the largest effect on expression. Strikingly, both tumor/normal context and this risk-associated haplotype play critical roles in mediating allelic *cis*-regulatory activity. Quantitative mass spectrometry for rs3769823 identified both E4F1, a transcriptional repressor, and IRF2, a transcriptional activator, as binding preferentially to risk-associated rs3969823-A. The binding of these transcription factors was validated via EMSA, supershift, and chromatin immunoprecipitation (ChIP) assays. The relative levels of E4F1 and IRF2 differ by cell-type and play a role in mediating transcriptional activity in a cell-type specific manner. Our results indicate that the top credible causal set variant rs3769823 likely influences expression of *CASP8* and *FLACC1* in a cell-type specific manner and may be a relevant functional variant for multiple cancers associated with this locus.

## Introduction

Over the past decade, genome-wide association studies (GWAS) have identified 54 genome-wide significant susceptibility loci for cutaneous malignant melanoma (CMM) in populations of European ancestry ^1–10^. Of these, Barrett and colleagues initially identified a genome-wide significant locus for melanoma on chromosome band 2q33.1 adjacent to the Caspase 8 gene (*CASP8*) ^5^. Multiple subsequent melanoma GWAS meta-analyses since have replicated this finding, including the most recent meta-analysis of 36,760 melanoma cases (rs10931936-T, *P* = 2.12 × 10^-12^, OR = 1.08)^10^. This region is a multi-cancer risk locus, with highly-correlated risk alleles for melanoma and other cancers sharing a common haplotype, including signals for breast cancer (rs3769821, *P* = 3.97 x 10^-18^, r^2^ with rs10931936 = 0.63) ^11^, keratinocyte cancers including both cutaneous basal cell carcinoma (BCC, rs6714430, *P* = 5.42 x 10^-55^, r^2^ with rs10931936 = 0.99), squamous cell carcinoma (SCC; rs10931936, *P* = 1.03 x 10^-7^), or both combined (rs6743068, *P* = 1.50 x 10^-48^, r^2^ with 10931936 = 0.99)^12,13^, and non-small cell lung cancer (rs3769821, *P* = 4.45 x 10^-8^, r^2^ with rs10931936 = 0.63)^14^. Additionally, the region also harbors a highly-correlated signal for prostate cancer (rs59308963, *P* = 2.41 x 10^-8^, r^2^ with rs10931936 = 0.93)^15^, where the protective allele is highly correlated with risk alleles for melanoma and the other cancers. However, whether causal variants and genes are shared across multiple cancers remains under-explored.

The topologically-associated domain (TAD) at 2q33.1 harbors multiple strong *a priori* candidate genes with well-established roles in apoptosis, including Caspase-8 (*CASP8*), Caspase 10 (*CASP10*), and the CASP8 and FADD-like apoptosis regulator (*CFLAR*). CASP8 is an apical caspase that initiates apoptosis through the extrinsic pathway via a subset of proteins in the tumor necrosis factor superfamily (TNFRSF) that includes TNFR1 (tumor necrosis factor receptor-1), CD95/Fas, and TRAIL-R (TNF-related apoptosis-inducing ligand receptor) ^16,17^. CASP8 is composed of a catalytic domain, a protease domain and two death domains comprising the prodomain, which is important for forming the death-inducing signaling complex (DISC), death receptor-FADD-CASP8 ^18,19^. Like all caspases, CASP8 is present as a monomeric procaspase zymogen^20^. Binding of CASP8 to the DISC facilitates zymogen maturation by dimerization and proteolytic cleavage of the caspase itself ^21,22^. Fully processed active CASP8 is released from DISC and activates downstream effector caspases. Notably, two proteins encoded by adjacent genes at this locus, *CASP10* and *CFLAR*, are highly similar to the N-terminus of *CASP8* and play a different role in the apoptotic pathway by interacting with FADD and CASP8 ^23,24^. While CASP10 is an apical caspase like CASP8, CFLAR splice variants inhibit different processing steps of CASP8 activation through heterodimerization ^24–26^. Given that most common trait loci are thought to function via *cis*-gene regulation, often at long distances, other genes within this TAD cannot be ruled out as candidate causal genes.

In this study, we fine-mapped multiple cancer GWAS datasets and integrate tissue-based and cell-type specific quantitative trait locus (QTL) data in order to identify likely causal variants and genes. We identified multiple highly-correlated potentially *cis*-regulatory variants, and establish rs3769823 as a key functional variant, likely acting via altered binding of the IRF2 and E4F1 transcription factors. We establish *cis*-regulation of *CASP8* as a likely causal mechanism underlying melanoma and breast cancer risk at this locus, and given the strong LD between signals, this mechanism may potentially play a role in risk of other associated cancers.

## Results

### Comparison of the 2q33.1 melanoma risk signal with GWAS of susceptibility to other cancers

The most recent GWAS meta-analyses of melanoma identified rs10931936 as the sentinel variant at the 2q33.1 risk locus (rs10931936-T, *P* = 2.12 × 10^-12^, OR = 1.08)^10^. Conditional and joint analysis of summary GWAS meta-analysis data^27,28^ identified no additional genome-wide significant association signals at this locus^10^. The next most significant signal within 1 Mb in either direction of rs10931936 is rs563855920 (*P*_conditional_ = 3.05 x 10^-4^; **Figure S1**; **Table S1**) more than 300 kb away, suggesting that there are no additional major signals.

This region has been identified as a genome-wide significant risk locus via GWAS of multiple other cancer types including breast cancer (122,977 cases, rs3769821-C, *P* = 3.97 × 10^-18^, 1000 Genomes EUR r^2^ with rs10931936 = 0.63)^11^ and keratinocyte cancers (47,742 cases, rs6743068-A, *P*_MTAG_ = 1.50 × 10^-48^, r^2^_rs10931936_ = 0.99)^13^, including cutaneous BCC (31,787 cases, rs6714430-C, *P* = 5.42 × 10^-55^, r^2^_rs10931936_ = 0.99)^13^ and SCC (9,674 cases, rs10931936-T, *P* = 1.93 × 10^-7^)^13^ where the genotypes of these risk alleles are correlated. To formally assess whether these traits may share common causal variants with melanoma risk, we compared melanoma fine-mapping data to other cancer datasets using both HyPrColoc^29^ and eCAVIAR^30^ and summary statistic data for the set of SNPs +/- 100 kb centered over the melanoma GWAS lead SNP. Colocalization analyses suggest a likely shared causal between melanoma risk and breast cancer (HyPrColoc PP = 0.93; maximum eCAVIAR CLPP = 0.04). Across the larger region, HyPrColoc finds the strongest evidence for shared causals for rs3769821 (HyPrColoc SNP score = 0.52, r^2^ = 0.63 to rs10931936) and rs3769823 (SNP score = 0.11, r^2^ = 0.77 to rs10931936) (**Figure S2, Tables S2** and **S3**). Colocalization analyses of keratinocyte cancers (both cutaneous BCC and SCC combined; PP = 0.99, maximum CLPP = 0.09), cutaneous BCC (PP = 0.98; maximum CLPP = 0.02), and SCC (PP = 0.96; maximum CLPP = 0.02) all show evidence for colocalization with melanoma risk, primarily showing the strongest evidence for melanoma lead SNP rs10931936 (SNP score is 0.42, 0.22 and 0.29, respectively for all keratinocyte cancers, BCC, and SCC) (**Tables S2** and **S3**). LocusCompare^31^ plots of regional summary data for melanoma compared to keratinocyte cancers or BCC suggest that the signals for these non-melanoma skin cancers may be complex and involve multiple potential signals (**Figures S3**-**S5**). Overall, these results suggest that a risk signal tagged by rs10931936 is common to all tested cancers and nominate several potential shared causal variants.

### Fine-mapping of melanoma risk-associated variants at 2q33.1

To comprehensively identify candidate causal variants within the 2q33.1 melanoma risk locus, we performed fine-mapping using multiple complementary approaches. Firstly, using GWAS summary data from Landi and colleagues^10^, we identified a set of 24 variants with a log-likelihood ratio (LLR) of up to 1:1000 relative to the lead SNP at the locus (rs10931936; **Figure 1**, **Table S4**). We also performed Bayesian fine-mapping using DAP-G^32,33^, identifying two clusters of potential candidate variants. The first cluster consisted of a 95% credible set of 15 variants (set 1; marked by rs10931936, PIP = 0.16; **Figure 1**, **Tables S6** and **S7**). A second cluster consisted of a set of six variants with cumulatively low posterior inclusion probabilities (PIP; set 2; led by rs77292590, PIP = 0.01; **Tables S5** and **S6**) consistent with conditional analysis of the melanoma GWAS that suggested no major second signal at this locus. Considering the possibility of a *cis*-regulatory function for this locus, we also fine-mapped using PAINTOR^34,35^, weighting the priors using melanocyte-specific epigenomic annotations from the RoadMap Epigenome Project and locations of a set of genes specifically expressed in melanocytes^36^ (**Figure 1**, **Table S7**), identifying a 95% credible set of seven SNPs. Of these, rs3769823, which was also fine-mapped by the other methods, had by far the highest posterior probability (0.789) of any variant. PAINTOR identified only one variant not in the DAP-G credible set 1 (rs74574949, r^2^ = 0.02 with rs10931936, DAP-G credible set 3, PIP = 0.0002; **Table S5**). Lastly, we assessed whether other variants not successfully imputed in the meta-analysis, and thus not directly fine-mapped, could also be credible causal variants. Here, we chose an LD-based cutoff of r^2^ = 0.625 to the lead variant, identifying an additional nine potential candidate causal variants. Combining variants from these analyses, we identified a total set of 27 credible causal variants for the melanoma risk signal (**Figure 1**, **Table S8**).

**Figure 1.**
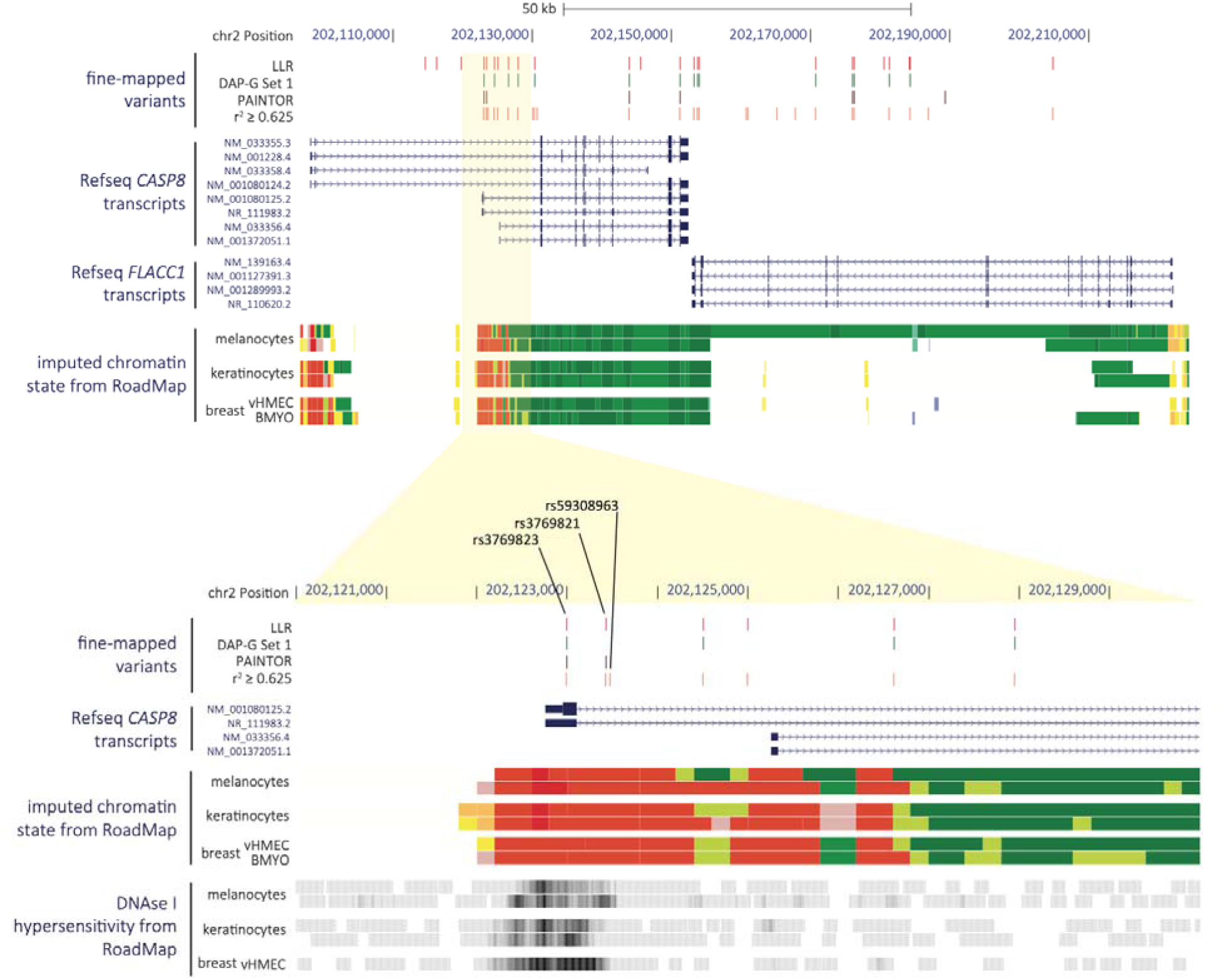
Fine-mapping of multiple melanoma GWAS signals on chromosome band 2q33.1. (Top) A view of the 2q33.1 locus (hg19) including all fine-mapped candidate causal variants nominated by log-likelihood ratio (LLR; red), Bayesian fine-mapping either unweighted (DAP-G, green) or weighted using melanocyte-specific epigenomic annotations (PAINTOR, brown), or LD to the melanoma lead SNP (rs10931936, r^2^ ≥ 0.625, orange). Also shown are tracks representing NCBI RefSeq genes and imputed chromatin state (ChromHMM) from two primary melanocyte cultures, two primary keratinocyte cultures, and two breast mammary cultures generated by the Roadmap Epigenome Project (Red: Active TSS; Orange-Red: Flanking Active TSS; Yellow: Enhancers; Green-Yellow: Genic enhancers; Green: Strong transcription, Dark Green: Weak transcription). (Bottom) A zoomed-in view of the same annotations for a region encompassing select candidate causal variants. Data and rendering was performed using the UCSC Genome Browser.

### Potential molecular mechanisms underlying the 2q33.1 locus

Of these fine-mapped variants, only one, rs3769823, is protein-coding, being a missense variant of codon 14 (K14R) of a single *CASP8* isoform (isoform G; ENST00000358485.4; RefSeq NM_001080125.2; **Figures 1** and **S6**), where the allele encoding K14 is associated with increased risk. We assessed potential functions via Jpred4, which suggested that CASP8^K14R^ may be located within a helical structure (**Figure S7**). To assess potential impacts on protein function, we used five different *in silico* prediction tools via PredictSNP2 ^37^. Of these, the majority (4/5) predicted rs3769823 to be benign; FunSeq2 however suggested that this variant may be deleterious (**Table S9**). While we cannot rule out a protein-coding function for rs3769823, these data suggest rs3769823 is benign.

We also assessed splicing QTLs (sQTL) from primary human melanocytes^38^. Notably, a significant risk-associated SNP, rs10804111 (*P*_GWAS_ = 6.79 × 10^-10^, LD to rs10931936 r^2^ = 0.62 and D’ = 1; **Table S1**), is a significant sQTL for alternative splicing of the *CASP8* exon 8 to 9 junction (*P*_sQTL_ = 7.33 × 10^-12^; **Figure S8**). Specifically, this generates an alternative *CASP8* transcript (isoform H; ENST00000339403.6, RefSeq NR_111983.2; **Figures 1** and **S6**), retaining part of intron 8 and leading to frameshift and premature termination of the *CASP8* transcript, which may act as a dominant-negative inhibitor of the caspase cascade^39^. Here, rs10804111-C is associated with higher melanoma risk and lower levels of the alternative isoform H (**Figure S8B**). To test the potential of rs10804111 to mediate alternative splicing, we generated C- or T-allele-specific mini-genes (**Figure S9A,** see material and methods). Using qRT-PCR in melanoma cells, we confirmed the association between rs10804111-C and lower usage of the exon junction for isoform H (exon 8-9 junction, *P* = 7.23 x 10^-5^; exon 8L-9 junction, *P* = 1.78 x 10^-3^; **Figure S9B**). However, we note fine-mapping of the melanoma GWAS signal does not identify rs10804111 as a member of any credible causal set at this locus, and formal colocalization between the melanoma risk signal and melanocyte sQTL signal did not provide significant evidence of colocalization.

Given the lack of strong support for protein-coding or splice variants explaining the 2q33.1 signal, as well as a clustering of fine-mapped variants within regions marked as promoter or enhancer in melanocytes (**Figure 1**, **Table S8**), we turned to eQTL analysis to explore the potential of this signal being explained by *cis*-gene regulation. We first investigated previously-published cell-type specific eQTL data generated from primary cultures of human melanocytes (n=106; **Table S10**)^36^. While multiple genes within the TAD harboring the association signal had nominally significant eQTL, *CASP8* is the only significant eQTL gene for the lead melanoma GWAS SNP (rs10931936; *P_CASP8_* _eQTL_ = 1.2 x 10^-9^, slope = 0.67, where lower *CASP8* expression associated with the T-risk allele; **Figure 2A**, **Table S10**)^36^. Conditional analysis of melanocyte *CASP8* eQTL identified multiple variants comprising only a single eQTL signal within this locus (**Table S11**). Colocalization analysis of the *CASP8* eQTL and melanoma GWAS summary data using both HyPrColoc^29^ and eCAVIAR^30^ suggest *CASP8* as a strong candidate gene (HyPrColoc PP = 0.97, eCAVIAR colocalization posterior probability/CLPP = 0.09 for rs3769823) with the strongest evidence of shared causal variants for rs3769823 (HyPrColoc SNP score = 0.86, eCAVIAR CLPP = 0.09, r^2^ to rs10931936 = 0.77) and rs3769821 (HyPrColoc SNP score = 0.14, eCAVIAR CLPP = 0.03, r^2^ to rs10931936 = 0.63) (**Figure 3**, **Table S12** and **S13**). Collectively these results suggest a shared common causal variant between melanoma risk and melanocyte-specific *cis*-regulation of *CASP8*. While no other genes had nominally significant melanocyte eQTLs for the sentinel melanoma risk SNP rs10931936, other strongly risk-associated SNPs at this locus were, notably for the *CASP8*-adjacent gene *FLACC1* (rs3769821, *P* = 2.42 x 10^-5^, r^2^ to rs10931936 = 0.63; lead *FLACC1* eQTL SNP rs796181752, *P* = 9.01 x 10^-6^; **Table S10**), where the direction of effect relative to risk allele is opposite to that for *CASP8*. Colocalization using HyPrColoc^29^ and eCAVIAR^30^ give discordant results, with eCAVIAR suggesting potential eQTL colocalization (CLPP = 0.04 for rs3769821; **Table S13**) while HyPrColoc does not (HyPrColoc PP < 0.5; **Table S12**). A LocusCompare^31^ plot suggests *cis*-regulation of *FLACC1* in melanocytes may be complex and involve multiple potential signals potentially explaining the discordant results (**Figure S10**). Collectively, these data suggest that the melanoma GWAS signal may be explained by *cis*-regulation of *CASP8* and *FLACC1*, with opposite directions of effect for the two genes.

**Figure 2.**
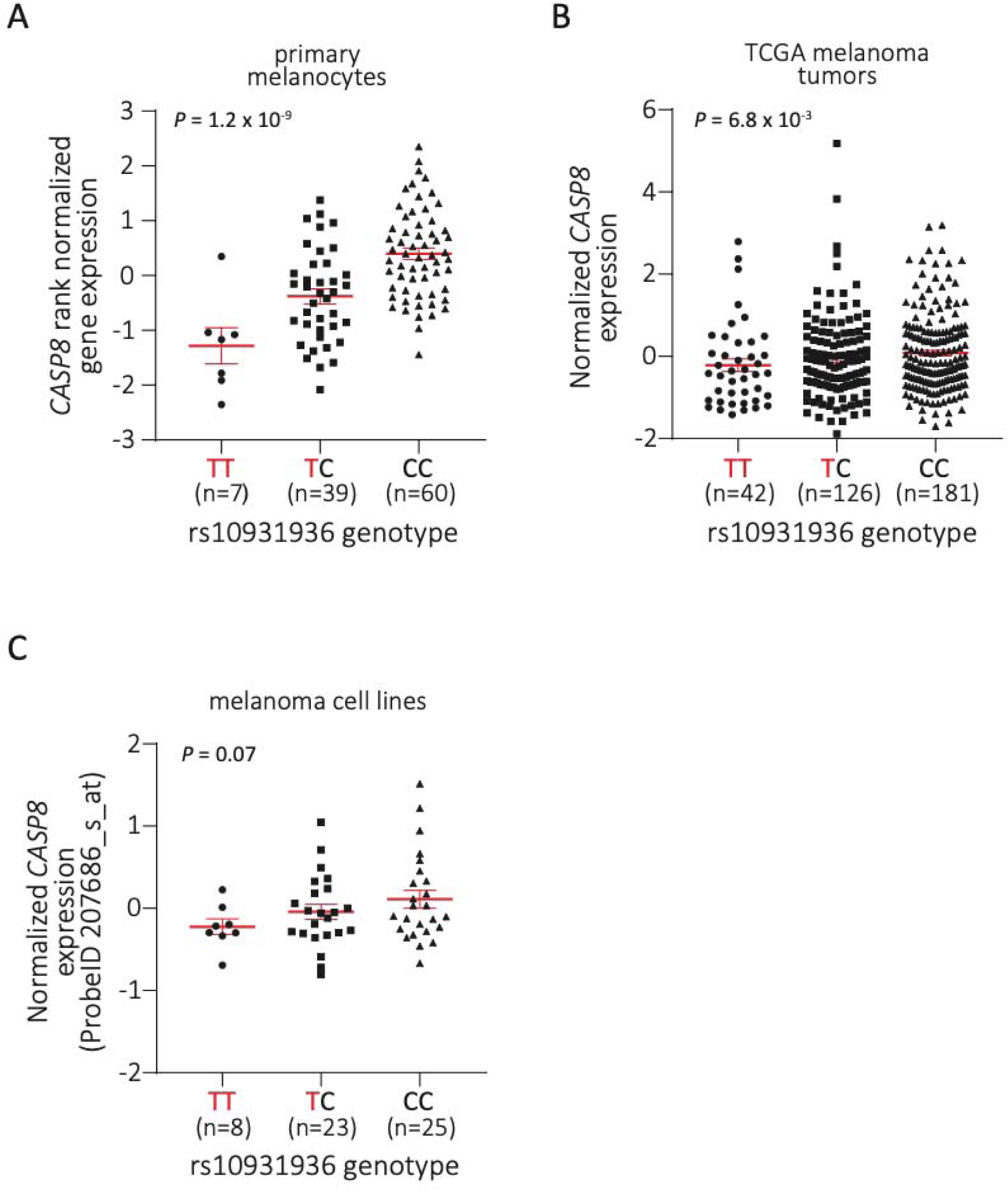
The melanoma risk-associated T allele of rs10931936 is correlated with lower *CASP8* expression. eQTL analyses were performed for rs10931936 combining genotype and expression level data derived from (A) 106 human primary melanocyte cultures (TT, *n*=7; TC, *n*=39; CC, *n*=60), (B) 349 melanoma tumors from TCGA-SKCM (TT, *n*=42; TC, *n*=126; CC, *n*=181), and (C) a panel of 59 early-passage melanoma cell lines (TT, *n*=8; TC, *n*=23; CC, *n*=25), where the risk-T allele is labeled in red. A significant eQTL effect with higher expression driven from the C-protective allele was observed for *CASP8* in both melanocytes and melanoma tumors (*P* = 1.2 x 10^-9^ and *P* = 6.8 x 10^-3^, respectively), while the result was marginal but in the same direction in melanoma cell lines (*P* = 0.07). Significance determined by linear regression; mean with SEM are plotted along with individual data values.

**Figure 3.**
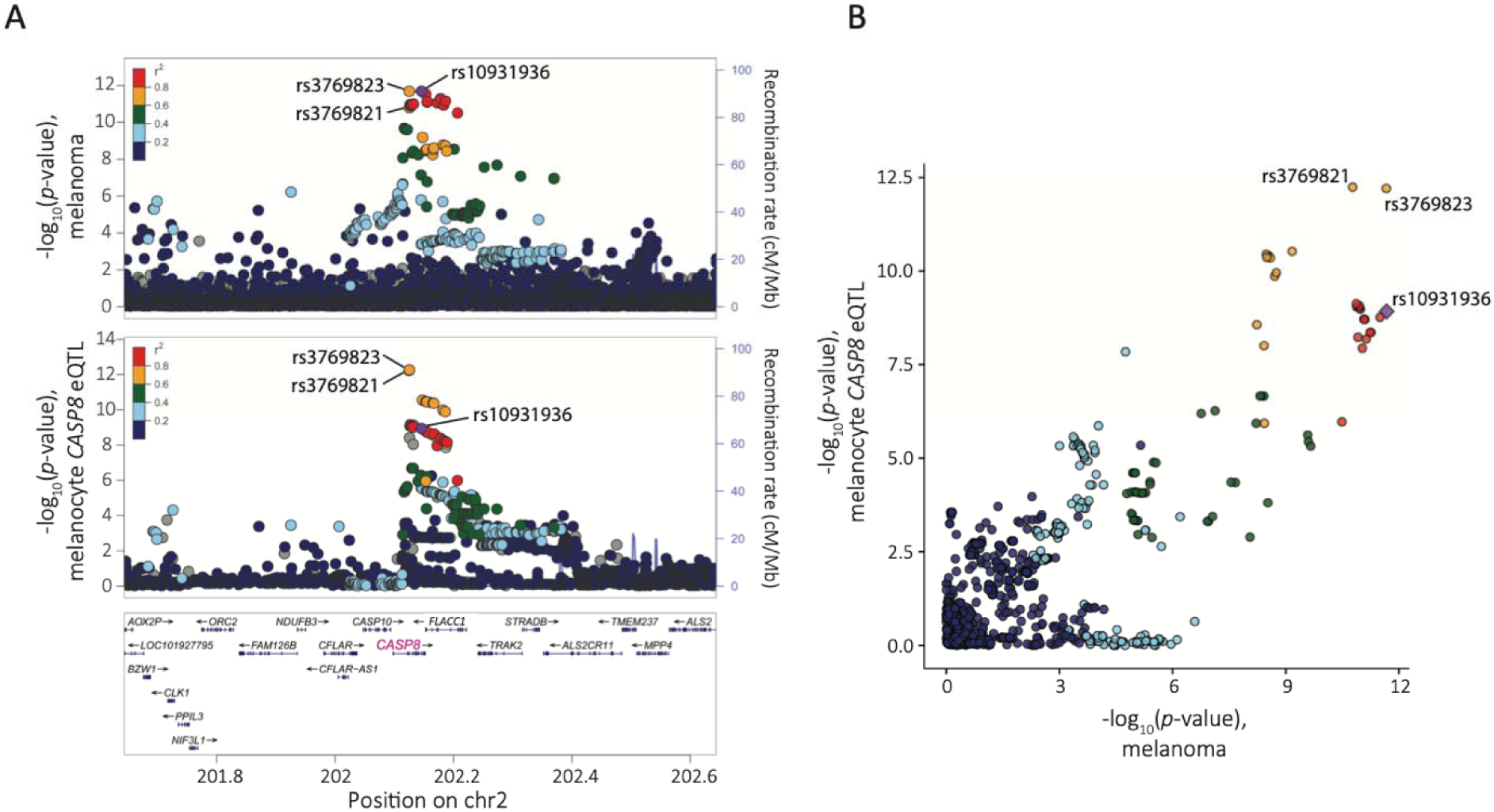
*CASP8* eQTL in human primary melanocyte cultures is colocalized with the melanoma GWAS. (A) LocusZoom plots present –log10 *P*-values for melanoma GWAS (upper) and melanocyte *CASP8* eQTL (lower) for a 1 Mb region encompassing rs10931936. The melanoma risk lead SNP rs10931936 is labeled and highlighted in purple in both panels, and LD (r^2^ based on 1000G EUR) of all other SNPs to the melanoma GWAS lead SNP is color-coded. (B) A LocusCompare plot compares *P*-values between melanoma GWAS (*x*-axis) and melanocyte *CASP8* eQTL (*y* axis) for the same region. Genomic coordinates are based on hg19.

These data are consistent with results from a transcriptome wide association study (TWAS) study we previously performed using the same GWAS summary statistics and melanocyte eQTL data^10^, where only *CASP8* is significant and *FLACC1* is marginal (*P_CASP8_*= 5.57 x 10^-12^; *P_FLACC1_* = 3.83 x 10^-5^; **Table S14**). Assessing models trained on each of 47 tissue types in the GTEx v.7; GTEx portal (https://gtexportal.org)^40^ identified *CASP8* as a significant TWAS gene in 30 out of 47 GTEx tissues, strikingly with the direction of effect for six tissues being opposite to that observed for melanocytes, including multiple brain tissues and pituitary (**Table S15**)^10^.

TWAS across GTEx tissues likewise found a significant positive association between imputed *FLACC1* levels and risk in 37 tissues (**Table S16**). We also assessed eQTLs in both 349 melanoma tumors from TCGA-SKCM^36,41,42^ as well as 59 early-passage melanoma cell lines using expression microarray data^43^. In TCGA melanomas, we observed an FDR-significant eQTL for *FLACC1* (*P* = 6.57 x 10^-4^), as well marginal eQTLs for *CASP8* (*P* = 6.82 x 10^-3^; **Figure 2B**) and *TRAK2* (*P* = 0.03) (**Table S17**). In melanoma cultures, the rs10931936 risk-T allele is marginally associated with multiple transcripts, including *CASP8* (*P* = 0.07; **Figure 2C**, **Table S18**). Both *CASP8* eQTLs were in the same direction as observed in melanocytes.

Finally, given potential colocalization of melanoma risk with signals for breast cancer and cutaneous SCC, we also assessed candidate genes specifically in mammary and skin tissues profiled by GTEx v.7^40^. Colocalization analyses with the melanoma risk signal suggests melanoma risk may share common causal variants with eQTLs for *CASP8* (eCAVIAR CLPP = 0.012 for rs3769821) and *FLACC1* (eCAVIAR CLPP = 0.05 for rs3769823) in mammary tissues, and *FLACC1* in skin-not-sun-exposed (HyPrColoc PP = 0.97; SNP score = 0.62 and CLPP = 0.013 for rs3769823) and skin-sun-exposed tissues (HyPrColoc PP = 0.79; SNP score = 0.15 and CLPP = 0.012 for rs3769823) (**Tables S19** and **S20**). TWAS of melanoma risk using mammary and skin tissues found both *CASP8* and *FLACC1* to be significant TWAS genes with the same direction of effect observed in melanocytes (**Tables S15** and **S16**). These results suggest that the 2q33.1 melanoma risk locus may not only share common causal variants with breast and other cancers, but also with *cis*-regulatory variants influencing *CASP8* and *FLACC1* levels in breast and skin tissues, suggesting a potential shared *cis*-regulatory etiology for multiple cancers.

### Prioritization of potential *cis*-regulatory variants using epigenomic data

Given the strong evidence for the 2q33.1 GWAS signals for melanoma as well as perhaps other cancers being explained by *cis*-regulation of *CASP8* or potentially *FLACC1*, we next prioritized credible set variants for assessment of allelic *cis*-regulatory activity. We prioritized for further study eight variants with active imputed ChromHMM chromatin states consistent with melanocyte gene regulatory regions as annotated by Regulome DB and HaploReg (**Figure 1**, **Table S8**; individual melanocyte regulatory marks are shown in **Figure S11**)^44^. Most of these eight variants also overlapped H3K27ac histone marks and or open chromatin (FAIRE-seq) from a panel 11 melanoma cultures^45^ (**Figure S12**). Given the shared GWAS signal between melanoma, breast cancer, and keratinocyte cancers, we additionally prioritized fine-mapped variants that overlap regulatory imputed ChromHMM states in primary keratinocytes and mammary epithelial tissues (**Table S8**); all such variants were already prioritized from melanocyte ChromHMM data. Thus, we proceeded to move forward with eight fine-mapped variants for functional assessment. Of these eight, three variants in close proximity (rs3769823, rs3769821, rs59308963, within 641 bp), are located in a single continuous region of active transcription start site (TSS)-proximal promoter state (TssA, **Table S8**)^44^ in melanocytes, keratinocytes, and mammary epithelial cells, overlapping the first exon and first intron of two annotated *CASP8* isoforms, isoform G and H (**Figures S11** and **S12**), potentially representing a *cis*-regulatory haplotype.

### Identification of functional *cis*-regulatory fine-mapped variants

To identify potentially functional *cis*-regulatory variants from these eight candidates, we performed individual reporter assays for each in the context of ∼140 bp of DNA sequence surrounding the variant cloned in both forward and reverse orientations and tested in two different melanoma cell lines (summarized in **Table S21**). Only one variant, rs3769823, showed a significant allelic difference in both forward and reverse orientations in both cell lines (**Figures 4A** and **S13**, **Table S21**), albeit with the opposite direction of effect relative to the melanocyte *CASP8* eQTL. rs3769821 also had a significant albeit weaker allelic effect in the reverse direction in both cell lines with the direction of effect matching the *CASP8* eQTL, while in one cell line in the forward direction there we noted a reproducible allelic difference in the opposite direction (**Figure S14**, **Table S21**). Lastly, two other variants, rs59308963 and rs1861270 (**Table S21**), showed significant but weak allelic difference across cell lines only in the reverse orientation (**Figure S15**, **Table S21**), with the effect direction for only rs59308963 matching the *CASP8* eQTL. We also examined two previously published melanoma MPRA studies which assayed five of these variants in human melanoma cells as well as HEK293T kidney cells (MPRA v.1^46^; **Figure S16**), or six of these variants in both melanoma cells and human primary melanocytes (MPRA v.2)^47^ with both studies testing a similar insert size (**Table S21**). Of the eight variants tested in either study, only rs3769823 showed an FDR-significant (FDR < 0.01) allelic effect in melanoma cells (*P*_MPRA v.1_ = 1.86 x 10^-12^, *P*_MPRA v.2_ = 1.09 x 10^-52^; **Table S21**), where the direction of effect was consistent with data from individual reporter assays and where the allelic effect is opposite to the *CASP8* eQTL. Likewise, in the second MPRA study, of the six variants tested, only rs3769823 was FDR-significant in primary melanocytes (*P*_MPRA v.2_ = 3.96 x 10^-9^), albeit with a smaller effect size (beta = -0.11) as compared to that observed in melanoma cells (beta_MPRA v.1_ = -0.65; beta_MPRA v.2_ = -0.45) (**Table S21**). Of the remaining variants, rs3769821 was significant only in a joint analysis of data from both melanoma and HEK293T kidney cells (*P*_MPRA v.1_ = 3.11 x 10^-4^; **Table S21**) with direction of effect matching that observed in individual luciferase assays (reverse orientation) and consistent with the *CASP8* eQTL. Given the potential of shared signal in this region with breast cancer, we also tested three variants, rs3769823, rs3769821, rs59308963, in multiple breast cancer cell lines replicating the significant allelic effect and direction observed in melanoma for rs3769823, but with inconsistent results for the other two variants (**Figures S13**-**S15**). Collectively, these data suggest the potential for multiple *cis*-regulatory variants, with risk-associated allelic effects in differing directions and in some cases possibly dependent on cellular context, with the strongest evidence for rs3769823.

**Figure 4.**
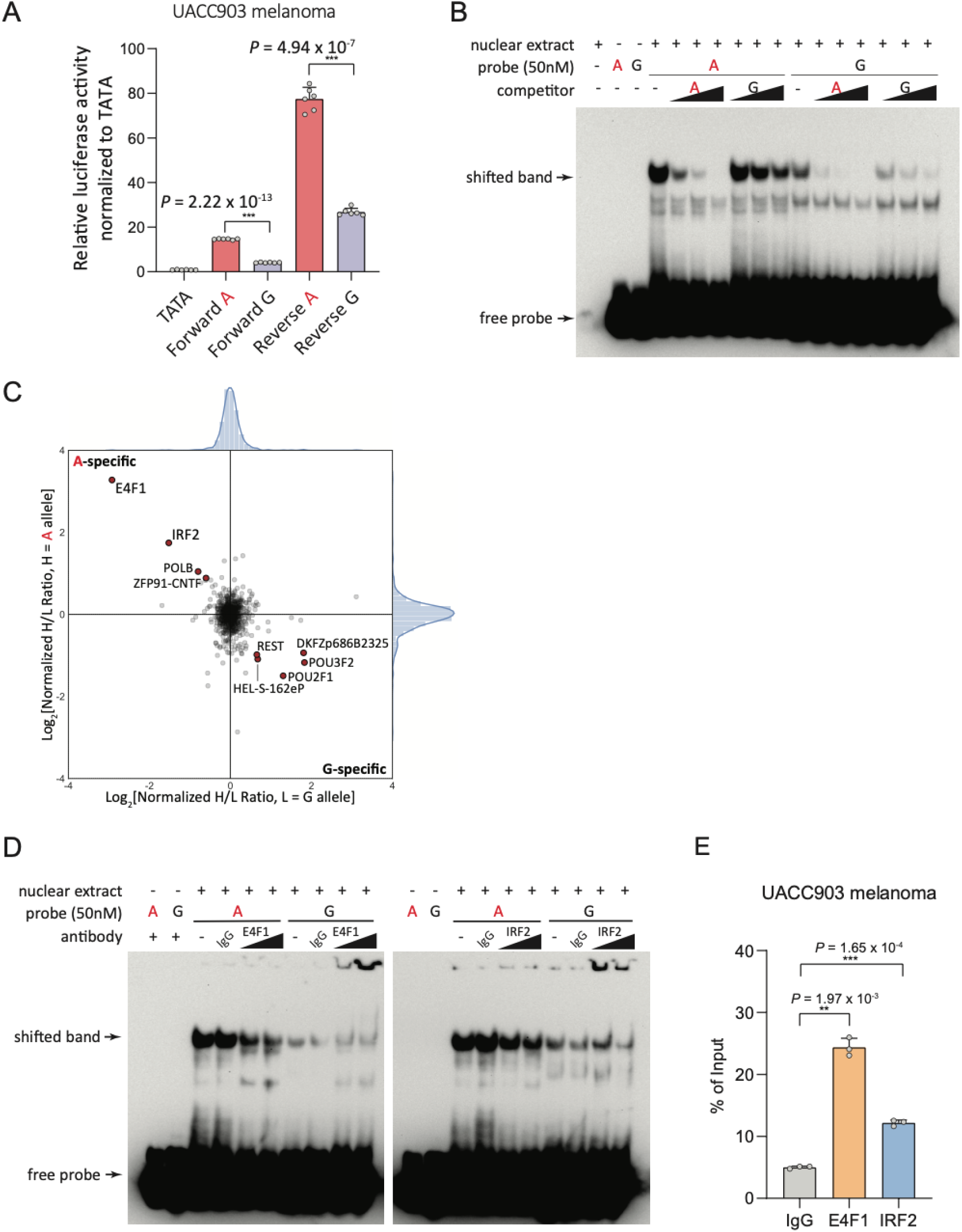
The melanoma-associated rs3769823 is a functional *cis*-regulatory variant and displays allele-preferential binding by E4F1 and IRF2. (A) Individual luciferase reporter activity assays for rs3769823 were conducted using the melanoma cell line UACC903. 138 bp sequences encompassing rs3769823 construct were cloned 5’ of the pGL4.23 minimal TATA promoter and transfected. Luciferase activity was measured 24 h after transfection and was normalized against *Renilla* luciferase activity. One representative set from three biological replicate experiments is shown. Mean with SEM are plotted. Individual *P* values are shown for A-risk allele versus G-protective allele for the replicate shown (two-tailed, unpaired t-test assuming unequal variances). TATA, minimal promoter control; A, risk allele construct marked in red; G, protective allele construct. (B) EMSAs were performed using 21 bp biotin-labeled double-stranded oligonucleotides for the A-risk (red) or G-protective (black) alleles of rs3769823 and nuclear extract from UACC903 melanoma cells. 2x, 5x, or 10x molar excess of unlabeled competitor was added in specified lanes. One representative set from three replicate experiments is shown. (C) Allele-specific rs3769823 binding proteins were identified by quantitative mass spectrometry using UACC903 melanoma nuclear extracts and 21 bp biotinylated double-stranded DNA probed with A-risk or G-protective alleles. The dimethyl-labeling ratios of proteins bound to A/G or G/A probes are plotted on the *x* and *y* axes for label-swapping experiments. Red-filled circles highlight proteins enriched above the background in both experiments. (D) Antibody super-shift assay of rs3769823 EMSA in UACC903 melanoma cells using anti-E4F1 or anti-IRF2 antibody is shown, where the A-risk specific shifted band (arrows) is diminished. One representative set from three independent experiments is shown. (E) ChIP was performed using antibody to anti-E4F1, anti-IRF2 or anti-IgG and chromatin prepared from UACC903 melanoma cells, followed by qPCR. DNA quantity was normalized to input DNA for each immunoprecipitation. Mean of qPCR triplicates with SEM are plotted and *P*-values shown for one representative experiment from three biological replicates.

Three of these variants are in close proximity and located within a common melanocyte enhancer element, and individual luciferase assays for all three reflect enhancer activity well above empty vector controls. We speculated that these may influence enhancer function in the context of a haplotype and thus performed reporter assays for a larger 641 bp region harboring all three variants in the context of the four most common haplotypes observed in the 1000 Genomes EUR sample set. Here, the haplotype harboring risk alleles of all three represent 27% of alleles, which that containing protective alleles for each is 64% (**Table S22**). Comparing the all-risk (A/C/ATTCTGTC for rs3769823, rs3769821, and rs59308963) to all-protective (G/T/-) haplotypes in melanocytes, we observed significantly lower reporter activity for the risk haplotype in the forward direction (cloned relative to the direction of *CASP8* transcription (**Figure S17A).** This direction matches the eQTL direction of effect for rs3769823 and *CASP8* in melanocytes, melanomas, skin, and mammary tissues (**Figure S18A-F**). In contrast, reporter assays in melanoma cells and breast cancer cells showed the opposite direction of effect, with higher reporter expression driven from the all-risk haplotype (**Figures S17B, C**). These data suggest a complex *cis*-regulatory architecture overlapping annotated melanocyte promoter for multiple *CASP8* isoforms, that allelic patterns of regulation may be highly dependent on both adjacent functional variants as well as cellular context, and where the observed direction of effect in melanocytes *in vitro* is consistent with *CASP8* as plausible causal gene.

### The genomic region containing rs3769823, rs3769821 and rs59308963 physically interacts with *FLACC1*

Given that rs3769823, rs3769821, rs59308963 are all correlated with not only *CASP8* expression but also *FLACC1* in melanocytes (rs3769823, *P* = 7.14 x 10^-4^, slope = -0.23, **Figure S18A**; rs3769821, *P* = 2.42 x 10^-5^, slope = -0.28; rs59308963, *P* = 0.01, slope = -0.19; **Table S10**), we investigated whether these or other credible causal variants physically interact with *FLACC1*. We first assessed interactions using a region-specific Capture-C assay performed in five independent human primary melanocyte cultures, where capture baits were tiled across the entire region of association ^48,49^ at 2q33.1. Here, we observed a direct physical contact between the restriction fragments containing these three variants and the *FLACC1* promoter (**Figure S19A**, **Table S23** and **S24**). We did not observe physical interaction between these variants and any other gene within the TAD containing the association signal. We further assessed this interaction using chromatin conformation capture (3C) in two independent melanocyte cultures as well as the UACC903 melanoma cell line, using the region harboring these three variants as bait and using primers spanning *CASP8* and *FLACC1*. Here, we confirmed a physical interaction between these variants and the *FLACC1* promoter in both melanocyte cultures, with little evidence of a strong association in melanoma cells (**Figure S19B**). These data suggest that the transcriptional regulatory region harboring rs3769823, rs3769821, and rs59308963 could potentially play a role in regulation not only of *CASP8*, but also *FLACC1*.

### Multiple transcription factors bind to rs3769823 in an allele-preferential manner

Given potential *cis*-regulatory roles for rs3769823, rs3769821, rs59308963, we further assessed whether these variants influence the binding of nuclear proteins to the sequences encompassing them. We first performed electromobility shift assays (EMSAs) using nuclear extracts from melanoma cell lines, primary melanocytes, or breast cancer cell lines. We observed preferential binding of nuclear protein to the A-risk allele compared to G-protective allele of rs3769823 across all cell lines tested, where competition with unlabeled probe containing the A-risk allele is stronger than that containing the G allele, demonstrating specificity (**Figures 4B** and **S20**). For rs3769821, we observed suggestive evidence of preferential binding of nuclear protein to the T-protective allele compared to C-risk allele in melanocytes, melanoma, and breast cancer cell lines, albeit with weak specificity (**Figure S21A**). Finally, for rs59308963, EMSA results show preferential binding to the insertion-risk allele compared to deletion allele with weak specificity in melanocytes, melanoma, and breast cancer cells (**Figure S22A**). These data suggest a strong and consistent pattern of allelic protein binding to rs3769823, consistent with the large effect observed in reporter assays, and weaker evidence for allelic binding for the other two variants.

We also performed quantitative mass spectrometry with oligonucleotides corresponding to the risk or the protective allele of each of these variants incubated with nuclear extracts from melanoma or breast cancer cell lines to directly identify allele preferential binding proteins. For rs3769823, we noted shared allele-specific binding proteins between both cell lines; specifically, IRF2 and E4F1 were observed to preferentially bind the A-risk allele, while REST and POU2F1 bound the G-protective allele (**Figures 4C** and **S23A**). The data for A-specific binding proteins are consistent with the patterns observed from the EMSA data, suggesting E4F1 or IRF2 as potential risk-allele binding transcription factors. Further, sequence-based motif prediction was also consistent with these mass spectrometry data, indicating that the sequence around rs3769823 forms a consensus binding site for both E4F1 and IRF2, favoring binding to the A-risk allele (**Figure S24**). For rs3769821, most of the potential preferential binding was found to be the T-protective allele via EMSA, however mass spectrometry found few T-specific binders and no overlapping T-binding proteins between cells; PRDM14 and ZBTB9 were both found to preferentially bind the C-allele (**Figure S21B**). We also identified BNC2 as a strong risk-insertion binding protein to the risk-associated allele of rs59308963 in melanoma cells (**Figure S22B**).

Based on the clear results and strong allelic pattern of *cis*-regulatory activity for rs3769823, we sought to verify whether E4F1 or IRF2 proteins bound the A-allele by using antibodies against these proteins in conjunction with EMSAs. Antibodies against either E4F1 or IRF2 consistently resulted in loss of A allele-specific protein binding in both melanoma and breast cancer cell lines (**Figures 4D** and **S23B**). To further establish the binding of E4F1 or IRF2 to rs3769823, we performed chromatin immunoprecipitation (ChIP) for E4F1 or IRF2 followed by quantitative PCR, noting an enrichment of binding at rs3769823 in melanoma cell lines with homozygous for rs3769823-A as well as primary melanocytes heterozygous for this variant (**Figures 4E** and **S25B**). We assessed enrichment at other locations across the *CASP8* gene that had previously been shown via ChIP-seq to be bound by IRF2 or E4F1, and DHS data from ENCODE (**Figure S25A**), however ChIP only showed enrichment of both in the region over rs3769823 (UACC903(A/A), *P*_E4F1_ = 1.46 x 10^-3^, *P*_IRF2_ = 0.016; C87(A/G), *P*_E4F1_ = 3.25 x 10^-3^, *P*_IRF2_ = 9.47 x 10^-3^; **Figure S25B**). Using the heterozygous melanocyte culture, we also assessed allelic enrichment in the immunoprecipitants using quantitative real-time PCR genotyping assay and noted a significant enrichment of the A allele as compared to the G allele at rs3769823 in primary melanocytes for both transcription factors (**Figure S25C**). Together, these data clearly suggest that E4F1 and IRF2 may act through rs3769823 at this locus.

### E4F1 and IRF2 as potential repressor and activator of *CASP8* transcription, respectively

Given the allelic binding observed for E4F1 and IRF2 to the A-risk allele of *cis*-regulatory SNP rs3769823, we next explored relationships between levels of these transcription factors and expression of *CASP8* and *FLACC1* in various cell and tissue types. In primary melanocytes, we observe clear negative correlation between both *E4F1* mRNA levels with *CASP8* (*E4F1*, *P* = 3.80 x 10^-6^, Pearson r = -0.43; **Figure S26**, **Table S25**) while *IRF2* is not significant (*IRF2, P* = 0.39, r = 0.08; **Figure S26**, **Table S25**); multiple linear regression shows both rs3769823 and *E4F1* to be significant (*E4F1 P* = 1.70 x 10^-15^; rs3769823 *P* = 5.39 x 10^-7^; *IRF2 P* = 0.43; **Table S26**). These data are consistent with a known role for E4F1 as a transcriptional repressor^50^ and suggests a role in regulation of *CASP8*. This direction of effect is consistent with transcriptional activity observed in reporter assays in melanocytes potentially being attributable in part of a repressive effect for E4F1 when considering the longer three-variant risk haplotype, but not the short (138 bp) fragment assays considering rs3769823 alone. In contrast, in TCGA melanoma tumors, while significant negative correlation with *E4F1* was observed (*E4F1*, *P* = 2.26 x 10^-4^, r = -0.20; **Figure S26**, **Table S25**), a much stronger positive with *CASP8* was observed for *IRF2* (*IRF2*, *P* = 1.69 x 10^-27^, r = 0.54; **Figure S26, Table S25**); both transcription factors and rs3769823 are significant via multiple linear regression (*E4F1 P* = 0.0018; *IRF2 P* = 2 x 10^-16^; rs3769823 *P* = 0.035; **Table S27**), again with the strongest effect for *IRF2.* We did not find significant evidence for an interaction between rs3769823 and either of the transcription factors (**Table S27**). We were able to replicate this positive correlation specifically between *IRF2* and *CASP8* in independent panels of melanoma cell lines (*P* = 4.23 x 10^-4^, r = 0.38; **Table S25**) and the Leeds Melanoma Cohort (*P* = 8.74 x 10^-4^, r = 0.30; **Table S25**). This direction of effect is consistent with a potential role for *IRF2* as a potential activator in both individual and haplotype-based reporter assays, but inconsistent with the direction of the relatively weak eQTL for CASP8 in melanoma tumors. We formally tested for potential interactions between *E4F1* and rs3769823 in melanocyte expression data as well as between *E4F1* and/or *IRF2* and rs3769823 in TCGA melanoma tumors but did not find significant supporting evidence (**Tables S26** and **S27**).

We also assessed transcripts encoding two T-protective allele binding proteins for correlations with potential target genes, observing a positive correlation between the transcriptional repressor *REST,* as well as *POU2F1,* and *CASP8* in melanocytes (*REST*, *P* = 7.21 x 10^-11^; r = 0.58; *POU2F1*, *P* = 8.40 x 10^-3^, r = 0.25), a weak negative correlation for *POU2F1* in TCGA melanomas (*P* = 0.04, r = -0.11), and a positive correlation for *REST* in the Leeds Melanoma Cohort (*P* = 1.75 x 10^-14^, r = 0.30) (**Table S25**). This positive correlation with *CASP8* in melanocytes and some melanoma tumors is inconsistent with the established role for REST as a transcriptional repressor. Still, this direction of effect is consistent with a potential role for REST repressing reporter activity in both individual (short fragment) reporter assays for rs3769823 as well as both previously-reported MPRA assays. Multiple linear regression considering all three individually correlated risk-A and protective-G binding alleles and genotype for rs3769823 in melanocytes suggests all four are significant (*REST* P = 6.71 x 10^-7^; rs3769823 *P* = 4.29 x 10^-6^; *POU2F1 P* = 0.0019; *E4F1 P* = 0.010; **Table S28**). We note similarly complex patterns of correlation between both risk- and protective-allele rs3769823 binding proteins in non-melanocytic tissues (GTEx skin, GTEx breast mammary, and TCGA breast invasive carcinoma; **Table S25**) and expression of *CASP8* as well as *FLACC1* (**Table S25**). These data are consistent with the activity of the regulatory region encompassing rs3769823 being modulated by a complex interplay of multiple allele-specific transcriptional regulators interacting with risk and protective alleles in a tissue-specific manner.

Finally, we also investigated correlations between *CASP8* and allele-specific binding proteins for rs3769821 (**Table S29**) and rs59308963 (**Table S30**), observing negative correlations with the transcripts for rs3769821 risk-allele binding protein ZBTB9 and the rs59308963 risk-allele binding protein BNC2 (*ZBTB9*, TCGA *P* = 2.15 x 10^-11^, TCGA r = -0.35, Leeds *P* = 3.40 x 10^-3^, Leeds r = -0.12; *BNC2*, TCGA *P* = 2.70 x 10^-4^, TCGA r = -0.19, Leeds *P* = 1.43 x 10^-13^, Leeds r = -0.29; **Tables S29** and **S30**). For *FLACC1*, we note similar correlations in melanocytes to those observed for *CASP8*, with negative correlation observed for *E4F1* (*P* = 3.40 x 10^-6^, r = - 0.43), and positive correlations observed for *REST* and *POU2F1* (*REST*, *P* = 1.72 x 10^-10^, r = 0.57; *POU2F1*, *P* = 4.71 x 10^-12^, r = 0.61), but do not observe consistent correlations for *IRF2* or *REST* across melanoma datasets (**Table S25**). Similar to what is observed for rs3769823, these correlations vary considerably between melanocytic and non-melanocytic tissues (**Tables S29** and **S30**) observing similarly complex patterns of correlation. Collectively, these data suggest a potentially complex gene regulatory interplay, prominently of multiple allele-specific rs3769823 binding proteins, as well as other transcriptional regulators, some of which may bind this region differentially in risk and protective haplotypes.

To determine the effects of depleting the two rs3769823 A-risk allele-specific binding proteins on *CASP8* transcription in melanoma, primary melanocyte cultures, and breast cancer cell lines, we knocked these genes down using siRNAs. Knockdown of *E4F1* resulted in an increase in *CASP8* levels while *IRF2* knockdown results in reduction of *CASP8* RNA levels in melanoma (**Figure 5A** and **5B**, **S27A** and **S27B**), breast cancer (**Figure S28A** and **S28B**), and melanocytes (**Figure S29A**), consistent with respective roles for E4F1 and IRF2 as a potential repressor and activator in cells of melanocytic lineage. Short fragment (138 bp) reporter assays for rs3769823 in conjunction with knockdown of A-risk allele binding protein IRF2 resulted in a significant decrease of expression driven primarily from the A-risk allele compared to the protective G-allele in both forward and reverse directions in melanoma (UACC903 cells, *P*_forward_ = 1.22 x 10^-7^, *P*_reverse_ = 2.84 x 10^-8^, **Figure 5C**; UACC1113, *P*_forward_ = 1.65 x 10^-5^, *P*_reverse_ = 2.20 x 10^-3^, **Figure S27C**) and breast cancer cell lines (T47D, *P*_forward_ = 2.92 x 10^-7^, *P*_reverse_ =3.18 x 10^-7^, **Figure S28C**), consistent with a potentially prominent role for IRF2 in *CASP8* regulation in melanoma cells. Consistent with the weaker correlations between *E4F1* and *CASP8* in melanoma cells, *E4F1* knockdown only modestly increased reporter activity for the A-risk allele, and only in the forward direction, suggesting a potentially weak repressor role for E4F1 in the context of melanoma cells (UACC903, *P*_forward_ = 8.05 x 10^-4^, *P*_reverse_ = 0.74, **Figure 5C**; UACC1113, *P*_forward_ = 3.47 x 10^-3^, *P*_reverse_ = 0.02, **Figure S27C**). In primary melanocytes, reporter assays for rs3769823 with knockdown of A-risk-allele binding E4F1 resulted in a significant increase of promoter activity in the A-risk allele compared to the protective G-allele in both direction (C23 melanocytes, *P*_forward_ = 1.27 x 10^-9^, *P*_reverse_ = 0.013, **Figure S29B**), suggesting a potentially repressor role for E4F1 in regulation of *CASP8* in melanocytes.

**Figure 5.**
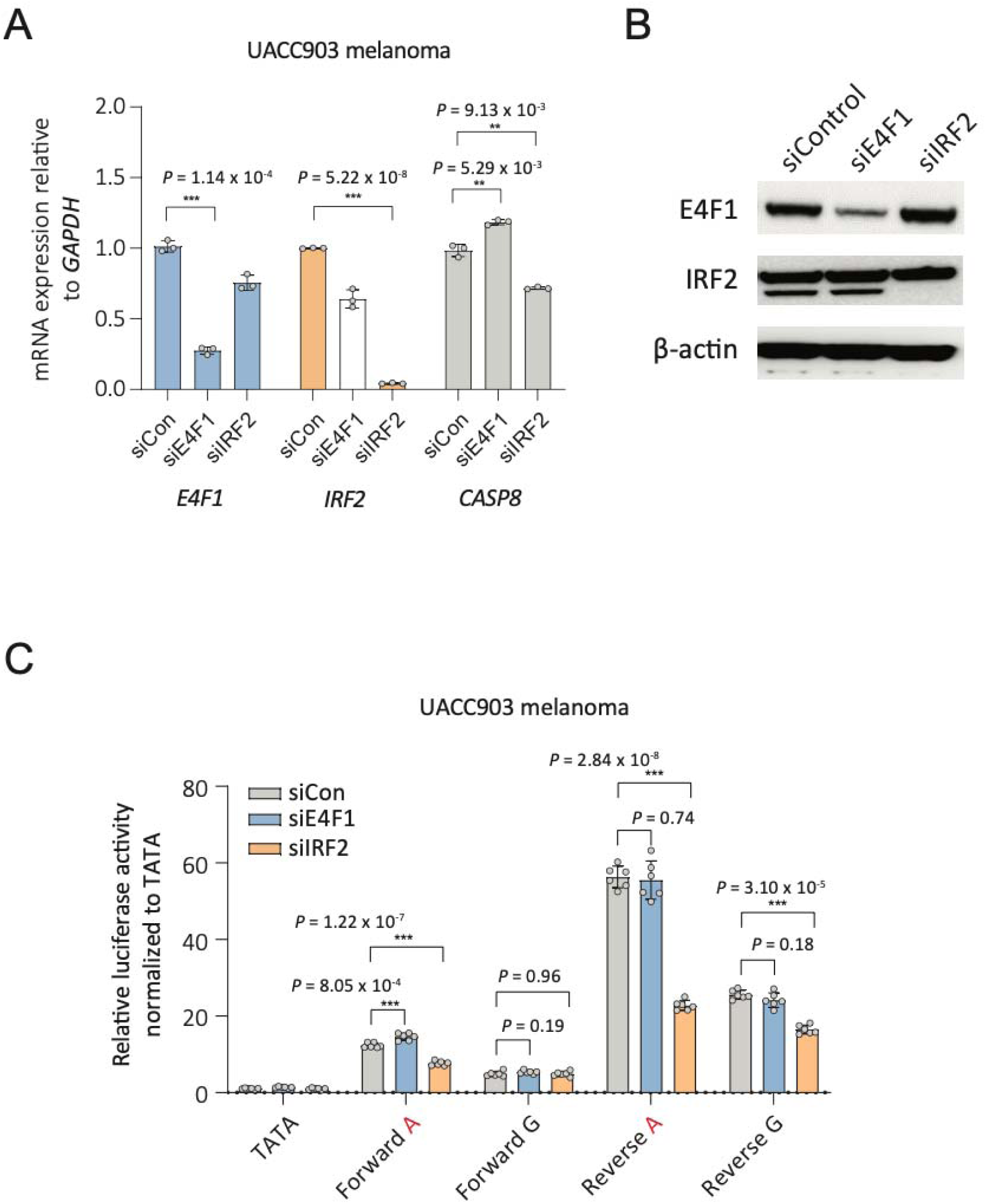
*E4F1* or *IRF2* and rs3769823 regulate *CASP8* expression in UACC903 melanoma cells. (A) *E4F1* or *IRF2* were knocked down using respective pools of four different siRNAs in UACC903 melanoma cells, and *E4F1*, *IRF2* and *CASP8* levels were measured. *GAPDH*-normalized *E4F1*, *IRF2* or *CASP8* mRNA levels are shown as fold change over those from non-targeting siRNA. A representative experiment from three biological replicates is shown (individual datapoints, mean, and SEM are plotted). *P*-values are shown from one representative set. (B) Protein levels were examined using anti-E4F1, anti-IRF2, or anti-GAPDH antibody with UACC903 cell lysates of siRNAs transfected with either non-targeting control, *E4F1*, or *IRF2* siRNAs. GAPDH was used as a loading control. One representative set of three replicate experiments is shown. (C) Individual luciferase assays were performed by co-transfecting rs3769823 luciferase constructs with *E4F1*, *IRF2,* or non-targeting control siRNAs into UACC903 melanoma cells. *Renilla*-normalized relative luciferase activities are shown relative to an empty construct containing only a minimal promoter (TATA). One representative set is shown from three biological replicates. Mean with SEM is plotted, *n* = 6 technical replicates. A two-tailed t-test assuming unequal variances was used to calculate all *P* values shown against control siRNA.

## Discussion

In this study, we focused on a melanoma susceptibility locus over the *CASP8* gene on 2q33.1 that was originally identified by Barrett and colleagues^5^ and has since been replicated in a large melanoma GWAS meta-analysis (rs10931936-T, *P*_fixed_ = 2.12 x 10^-12^, *P*_random_ = 2.17 x 10^-8^, OR = 1.08)^10^. In order to comprehensively identify additional independent signals as well as nominate one or more sets of credible causal variants associated with such signals, we used multiple complementary approaches. Conditional and joint analyses previously found no evidence for a second genome-wide significant signal at 2q33.1^10^; here, we observe no evidence for a prominent independent melanoma risk signal of marginal significance. Fine-mapping using Bayesian approaches similarly identified one prominent signal at this locus with little evidence for additional signals.

Germline variants is this region have been identified as associated with multiple cancers, including breast cancer (rs3769821) ^11^, keratinocyte cancers (cutaneous BCC and SCC combined, rs6743068) ^13^, cutaneous BCC (rs6714430) ^13^, cutaneous SCC (rs10931936) ^13^, and non-small cell lung cancer (rs3769821) ^51^ where the direction of effect is the same and the risk alleles share a common haplotype with the melanoma risk allele of rs10931936. In addition, a highly correlated risk signal was previously identified for prostate cancer (rs59308963) ^52^ where the direction of effect is opposite to that of melanoma and the other cancers. This locus was also identified via GWAS for chronic lymphocytic leukemia (CLL; rs3769825) ^53^, however this signal appears to be considerably less correlated with the melanoma risk signal (r^2^ with rs10931936 = 0.26). For several of these cancers with summary data readily available, we evaluated evidence for colocalization, with the results suggesting sharing of one or more causal variants between melanoma and signals for breast cancer, cutaneous BCC, and cutaneous SCC and further suggesting that our work to fine-map identify functional, potentially causal variants for melanoma risk may be relevant for these other cancers.

Using multiple complementary fine-mapping approaches, we identified a set of 27 potential causal variants. Collectively, these variants present multiple potential alternative or complementary functional mechanisms for influencing gene function and cancer risk. Notably, rs3769823 is a missense protein-coding variant in codon 14 of an alternative first exon of *CASP8* utilized by multiple isoforms and expressed in human primary melanocytes (isoforms G and H; **Figure S6**). However, a majority of algorithms designed to predict variant function suggest this variant is likely benign and K14R is not located within conserved domains or known or predicted sites of post-translational modification. Despite this, we did not directly test for an effect of the two alleles of this variant on CASP8 protein function and cannot rule out a role for altered protein sequence as a mechanism underlying risk at this locus. Zhang and colleagues studied the functionality of rs3769823 by expressing the two alleles in lung cancer cell lines and noted differences in proliferation and cell migration *in vitro* and *in vivo* ^54^. Notably, expression from the K14R (protective) allele generated more CASP8 protein, which the authors posited may be due to altered protein stability. Downstream phenotypic assays did not distinguish between the effects of protein abundance and protein sequence-specific effects, and potential transcriptional differences for the two constructs were not controlled for, so a protein-coding function for this variant on CASP8 remains possible but still unclear. Further work is needed to assess potential effects of this protein-coding change on CASP8 protein function.

We also observed a significant *CASP8* sQTL in melanocyte and other GTEx eQTL datasets relevant to cancers with associations for this locus (marked by rs10804111; r^2^ to rs10931936 = 0.62; D’ = 1) reflecting alternative splicing events at the exon 8 to exon 9 junction. The alternative splicing event is more strongly associated with the protective allele generates an alternative *CASP8* transcript, isoform H. This transcript retains only part of canonical intron 8 and introduces a frameshift in the transcript, leading to premature termination. The resultant protein retains the *CASP8* death effector domain but lacks the catalytic domain, and notably, Himeji and colleagues demonstrated that this protein could inhibit apoptosis mediated by wild-type CASP8 in a dominant-negative manner^39^. Thus, the transcript preferentially driven by the protective allele could potentially lead to loss of *CASP8* function; this stands in functional contrast to the melanocyte eQTL observed where the protective allele of rs10931936 is associated with higher rather than lower levels of *CASP8* transcript. Nonetheless, while LD of rs10804111 with rs10931936 is comparable to that of other fine-mapped credible causal variants for melanoma at this locus, none of the fine-mapping approaches we applied nominates rs10804111 as a credible causal variant itself. Still, much as for the protein-coding change caused by rs3769823, we cannot rule out a role for altered *CASP8* splicing contributing to melanoma risk.

In contrast, eQTL and TWAS results in human primary melanocytes^10,36^ provide strong evidence suggesting *cis*-gene regulation as a likely mechanism underlying melanoma risk attributable to this locus, where melanoma risk-associated alleles were significantly correlated with lower levels of *CASP8*. Both TWAS and colocalization approaches suggest that melanoma risk and melanocyte eQTL for *CASP8* likely share one or more common causal variant, with colocalization strongly nominating rs3769823. Both also provide some support for *FLACC1*, where in contrast to the *CASP8* eQTL, higher *FLACC1* levels are associated with the melanoma risk allele. Several fine-mapped melanoma risk variants at this locus are clearly located within melanocyte gene-regulatory regions, including regions annotated as active promoters, consistent with a potential role for this signal regulating *CASP8* levels. Consistent with observed *FLACC1* QTLs, melanocyte-specific capture-C and 3C assays showed a physical interaction between fine-mapped melanoma risk variants and the *FLACC1* promoter region (**Figure S19**) suggesting risk variants indeed may play a role in regulating both genes. Notably, none of these analyses suggest a prominent role for other nearby genes, including the apoptosis regulatory genes *CASP10* and *CFLAR*.

Based on these data, we comprehensively assessed fine-mapped risk variants for potential *cis*-regulatory function, first identifying eight variants overlapping with *cis*-regulatory histone marks in disease relevant tissue (melanocytes, keratinocytes, human mammary epithelial cells, and breast myoepithelial primary cells). We assessed each for allelic *cis*-regulatory potential both in two previously published massively parallel reporter assays performed in melanocytes and/or melanoma cells, identifying rs3769823 as the only significant variant, with allelic effects in both melanoma cells and to a lesser degree melanocytes. We likewise tested each in individual luciferase reporter assays in two melanoma cell lines, again identifying rs3769823 as the only variant with consistently significant allelic *cis*-regulatory activities in both cell lines and in both forward and reverse orientations where the direction of reporter effect was consistent with the melanocyte *FLACC1* eQTL but opposite that of *CASP8* (e.g., the risk allele is associated with higher reporter expression). Nonetheless, in individual assays, several variants displayed allelic activity in one orientation, including two additional variants rs3769821 and rs59308963, located nearby rs3769823 in the same annotated gene promoter region, suggesting potential functional contributions from more than one variant. Testing commonly occurring rs3769823/rs3769821/rs59308963 haplotypes in a reporter assay revealed cell-type dependent effects; the risk-associated A/C/ATTCTGTC haplotype was associated with lower expression relative to the protective G/T/-haplotype in melanocytes but higher expression in melanoma cells in the forward orientation relative to *CASP8* transcription (**Figure S17A, B**). This stands in contrast to the results from Camp and colleagues, who tested risk and protective haplotypes of rs3769823/rs3769821 and observed lower reporter gene expression driven from the risk haplotype in both normal breast and breast cancer cell lines ^55^. The differing direction of effect observed in melanocytes for rs3769823 in short-fragment versus longer-fragment haplotype analysis suggests that while rs3769823 is strongly functional, the allelic effects are influenced by other nearby and potentially functional variants.

All three variants showed allelic patterns of protein binding via EMSA. Previously, in the context of lung cancer susceptibility, Long and colleagues observed a *cis*-regulatory element shared across many lung cell types overlapping rs3769823 and noted that this variant is predicted to alter IRF8 DNA binding ^56^. Here, we applied a quantitative mass spectrometry workflow and identified multiple allele-specific nuclear binding proteins from both melanoma and breast cancer cells that bound these variants in an allele specific manner. In particular, we identified proteins that typically act either as a transcriptional repressor (E4F1) or activator (IRF2) as both preferentially binding the A-risk allele of rs3769823, and verified binding of both factors via ChIP in melanoma cells and primary melanocytes. While we did not observe significantly different IRF8 binding in this context, the DNA-binding motifs of IRF2 and IRF8 are quite similar. We observed a similar situation for G-protective allele specific binding proteins, where both an activator (POU2F1) and repressor (REST) were found to bind to this allele in an allele preferential manner. Thus, allelic transcriptional activity may be highly dependent on relative availability of these factors as well as competition for binding under different cellular contexts. Indeed, gene expression correlation analyses suggest, for example, that E4F1 may play a prominent role for *CASP8* regulation in melanocytes, while the activator IRF2 may play a larger role in regulation of *CASP8* in melanomas. Indeed, reporter assay data for rs3769823 in conjunction with *E4F1* or *IRF2* knockdown are consistent with this, where in melanoma cells knockdown of IRF2 resulted in a significant decrease of expression driven from the A-risk allele, while knockdown of E4F1 resulted in a weaker increase of reporter activity driven from the A-allele. Given the similarity of DNA-binding motifs for IRF proteins, it may be possible that in addition, tissue-specific availability of other IRF family members could also drive or repress gene expression via rs3769823 and possibly explain the pleiotropy at this signal for prostate cancer where the direction of effect is opposite that of melanoma and the other cancers ^15^. In contrast knockdown of E4F1 in melanocytes resulted in a significant increase of activity driven from the A-risk allele. We further observed additional allele specific binding proteins for potentially functional adjacent variants rs3769821 and rs59308963, which could well further modulate transcriptional output themselves as well as potentially contribute to preference of transcription factor binding to rs3769823.

Our work provides strong evidence for two potential causal genes at this locus, *CASP8* and *FLACC1*. Of these, *CASP8* represents a very strong *a priori* candidate cancer risk gene, given its long-established role as an apoptotic initiator caspase and the fact that lower *CASP8* levels are associated with cancer risk. Little is known about the function of *FLACC1* (formerly *ALS2CR12*). It has been found to be component of sperm tail flagellum ^57^, and functional studies of *ALS2CR12*-like genes in *C. elegans* suggest that similar genes play a role in gene and or repetitive DNA silencing, a function appearing to be dependent on the biogenesis of small RNAs ^58,59^. Notably this region harbors multiple genes involved in apoptosis, including *CASP10*, encoding another apoptotic initiator caspase ^60–62^ sharing similar substrate specificity to CASP8 ^63^ but possibly acting as a negative regulator of CASP8-mediated cell death ^64^. Also located within this region is *CFLAR* (also known as c-FLIP), encoding a protein structurally-similar to CASP8 but lacking caspase activity and acting to prevent CASP8 dimerization and inhibiting apoptosis ^19,65–69^. Despite the interacting roles of these genes, we observe little evidence that the risk signal drives differences in expression of these antiapoptotic genes. Nonetheless, we cannot rule out potential regulation of these genes from this signal under specific contexts. We do note, however, that at least in human melanocytes we observe no evidence of physical interaction between fine-mapped cancer risk variants (**Figure S19**) and the promoter regions of either gene.

In summary, our study functionally interrogated the multi-cancer risk locus on chromosome band 2q33.1, providing strong evidence for *cis*-regulation of *CASP8* via multiple functional variants as a primary causal mechanism influencing cancer risk.

## Material and methods

### Melanoma GWAS summary data

For all analyses using melanoma GWAS data, we used summary statistics from both histologically confirmed, as well as self-reported melanoma cases (from 23andMe and UK Biobank), and controls as previously described^10^. In total, summary data are derived from 36,760 melanoma cases and 375,188 controls; all participants provided informed consent and participation was IRB approved. Participants from 23andMe provided informed consent and participated in the research online under a protocol approved by the external AAHRPP-accredited IRB, Ethical and Independent Review Services (E and I Review).

### Conditional and joint analysis of melanoma GWAS

Conditional and joint analysis was performed with summary melanoma GWAS meta-analysis data by using Genome-wide Complex Trait Analysis (GCTA, v.1.26.0)^28^ to identify independent associated variants. We set the collinearity threshold (-cojo-collinear) to R^2^ = 0.05 to detect only completely independent SNPs. Linkage disequilibrium (LD) between SNPs was calculated using 5,000 randomly selected individuals of European ancestry from UK Biobank.

### Colocalization of the 2q33.1 melanoma association signal with other GWAS and QTL data

For colocalization analyses, we used GWAS summary data from breast cancer (122,977 cases and 105,974 controls; dbGaP accession number phs001265.v1.p1; downloaded from https://bcac.ccge.medschl.cam.ac.uk/bcacdata/oncoarray/oncoarray-and-combined-summary-result/gwas-summary-results-breast-cancer-risk-2017/)^11^ and keratinocyte cancers (47,742 cases and 634,414 controls; dbGaP accession number phs000360.v3.p1)^13^, including cutaneous BCC (31,787 cases and 619,351 controls)^13^ and cutaneous squamous cell carcinoma (SCC; 9,674 cases and 625,657 controls)^13^. Colocalization analyses for the 2q33.1 locus were performed using both Hypothesis Prioritization in multi-trait Colocalization (HyPrColoc)^29^ and eQTL and GWAS Causal Variant Identification in Associated Regions (eCAVIAR)^30^ as implemented in the ezQTL web-based tool (https://analysistools.cancer.gov/ezqtl/)^70^. For both, we used the set of SNPs in +/- 100 kb window size from the melanoma GWAS lead SNP rs10931936^36,38^. For HyPrColoc, a posterior probability (PP) above 50% (0.5) was considered to display a positive colocalization and SNP score above 5% (0.05) was considered to share a common causal variant. For eCAVIAR, the colocalization posterior probability (CLPP) score was calculated with a maximum number of two causal SNPs in the locus; a CLPP score above 1% (0.01) was considered to display a positive colocalization.

### GWAS fine-mapping

Fine-mapping was performed using several approaches. Firstly, in order to inclusively nominate candidate causal variants for assessment, we used the log-likelihood ratio (LLR) of each SNP in the region relative to lead variant rs10931936, and considered all variants within a ratio of >1:1000. Variants not successfully imputed from the GWAS that thus could not be fine-mapped in this manner nor via other methods requiring summary data, hence we used an LD-based threshold. Specifically, we chose this threshold to be consistent with the observed LD (r^2^) between LLR-mapped credible set variants with, where the minimum LD from this set with rs10931936 was r^2^ = 0.625 (1000 Genomes Projects Phase 3 EUR data; 1000G EUR). We thus considered variants not present in the summary dataset where 1000G EUR r^2^ ≥ 0.625. We also performed Bayesian fine-mapping of melanoma GWAS summary data^10^ for the 2q33.1 risk region using both Deterministic Approximation of Posteriors (DAP-G)^32,33^ and Probabilistic Annotation INtegraTOR (PAINTOR)^34,35^. DAP-G (version 1.0) analysis used a 500 Kb window size centered over the association signal (median position of chr2:202,123,966) and allowing for a maximum number of five causals. A UK Biobank (UKBB) LD reference panel (from 337K UKBB participants, downloaded from https://alkesgroup.broadinstitute.org/UKBB_LD/)^71^ was aligned with the summary statistics from the melanoma GWAS meta-analysis^10^ using R (version 4.1.0) for input into DAP-G. PAINTOR analyses were performed as described previously^46^ using PAINTOR 3.0 with a window size of 100 Kb, maximum causals set to 4, 4 melanocyte-specific epigenomic annotations and pairwise LD derived from 1000 Genomes phase 3 EUR (1000G EUR) data computed using PLINK version 1.9 and R version 4.1.0. Functional annotations include a set of 2,000 melanocyte-specific expressed genes from our melanocyte dataset^36^, melanocyte enhancers, transcribed regions, and a histone mark (H3T11ph) from ENCODE and Roadmap.

### Prediction of variant effect on protein function and structure

We predicted the functional impact of missense variant rs3769823 on function of CASP8 protein through the PredictSNP2 ^37^ web-based tool (https://loschmidt.chemi.muni.cz/predictsnp2/), which integrates five functional prediction methods including CADD (Combined Annotation Dependent Depletion) ^72^, DANN (Deleterious Annotation of genetic variants using Neural Network) ^73^, FATHMM (Functional Analysis Through Hidden Markov Model) ^74^, FunSeq2 ^75^, and GWAVA (Genome-Wide Annotation of VAriants)^76^. We also predicted potential effects of CASP8^K14R^ on secondary structure of the protein through JPred4 ^77^ web server (https://www.compbio.dundee.ac.uk/jpred4/), which also predicts solvent accessibility and coiled-coil regions. Predictions from AlphaMissense^78^ for rs3769823 are not available.

### Quantitative trait locus (QTL) data and other expression data

QTL data from 106 primary human melanocyte cultures primarily of European ancestry were generated as described previously for both gene expression (eQTL)^36^ as well as for splicing (sQTL) and CpG methylation (meQTL)^38^ (dbGaP accession: phs001500.v1.p1). Conditional analysis of the *CASP8* melanocyte eQTL was performed using QTLTools (https://qtltools.github.io/qtltools/)^79^, combining genotype and expression level data, along with a matrix of covariates including the top three genotype principal components and the top 15 PEER factors. For other tissues, we used pre-analyzed *cis*-eQTL data from the Genotype-Tissue Expression (GTEx) project (v.7), which were downloaded from the GTEx portal (https://gtexportal.org/home/index.html)^40^. eQTL analysis was performed previously^36^ using relative linear copy-number values as a covariate for 349 melanoma tumors from The Cancer Genome Atlas Project (TCGA) skin cutaneous melanoma (SKCM) dataset^41,42^. eQTL data for 59 early passage melanoma cell lines was performed as described previously^43^ using expression and genotyping data from NCBI GEO (accessions GSE78995 for expression data; GSE99193 for genotyping data). We specifically assessed genes within the TAD harboring the lead SNP at 2q33.1 (rs10931936; chr2:201,760,000-203,080,000; hg19) based on Hi-C results from SKMEL5 melanoma cells as visualized by the 3D Genome Browser (http://3dgenome.fsm.northwestern.edu/view.php).

### *CASP8* isoform analysis

To assess splicing QTLs (sQTLs), we used the same genotype data, population structure covariates, and statistical approaches as used for melanocyte eQTL analysis^36^, replacing normalized gene expression levels with normalized splice junction events. STAR^80^ was used to map the RNA-Seq reads onto the genome (hg19) and then LeafCutter^81^ was applied to quantify the splice junctions following the procedures described by the authors (http://davidaknowles.github.io/leafcutter/articles/sQTL.html). Taqman real-time PCR assays targeting unique junctions of *CASP8* transcript isoforms were obtained from Thermo Fisher Scientific (Waltham, MA; all isoform transcripts: Hs01018151_m1; exon 8 to 9 junction: Hs01018160_m1; exon 8L (alternative spliced) to 9 junction: Hs04405665_m1). RNA was isolated from primary melanocyte cultures derived from 106 individuals mainly of European decent^36^, and cDNA was synthesized using iScript^TM^ Advanced cDNA Synthesis Kit (Bio-Rad, Hercules, CA). Taqman assays were performed in triplicates (technical replicates) and normalized to *TBP* or *PPIA* levels.

### Exon-trap analysis of alternatively spliced exons of *CASP8* genes

rs10804111 is an intronic variant; no variants strongly linked to the *CASP8* melanocyte sQTL were located in consensus splice or branch point sequences, and none could create a cryptic splice site responsible for the observed allelic splicing pattern. We therefore assessed potential effects of the lead sQTL SNP rs10804111 in a minigene assay. The intron from the exon 8L to 9 junction is >5,500 bp and not amenable to cloning in a simple minigene vector. We therefore constructed minigene vectors to contain *CASP8* exons 8/8L and surrounding intron (100 bp upstream of exon 8 and 100 bp downstream intronic sequence from the exon 8L junction) and *CASP*8 exon 9 and surrounding intronic sequence (100 bp upstream and 100 bp downstream intronic sequence), linked together with 201 bp of intronic sequence surrounding each allele of rs10804111 (**Figure S8**). *CASP8* exons 8 to 9 (including intron 8; 1,181 bp), along with the 5’ flanking of exon 8 and 3’ flanking of 9 sequences (100 bp for each) were custom-synthesized from Thermo Fisher Scientific and cloned in sense orientation using XhoI and BamHI restriction sites of the Exontrap vector pET01 (MoBiTec, Germany) to generate the pET01-rs10804111(C/T) mini-gene constructs containing each allele of rs10804111. Sequence-verified pET01 constructs were transfected into UACC903 melanoma cells using Lipofectamine^TM^ 2000 (Invitrogen, Waltham, MA) in 6-well format. Cells were harvested 48 h after transfection, and total RNA was extracted with the QIAGEN QIAcube using RNeasy kit with on-column DNase I treatment (Qiagen, Germany). 1 μg of total RNA was converted into cDNA with SuperScript^TM^ III reverse transcriptase (Invitrogen) and a vector-specific cDNA primer listed in **Table S31**. Taqman assays targeting unique junctions of pET01-rs10804111 were obtained from Thermo Fisher Scientific (vector exon 2 (VE2) as a control: APH6CGV; VE1 to VE2 junction: APPRMTJ; *CASP8* exon 8 to exon 9 junction: APKA42T; *CASP8* exon 8L to exon 9 junction: APNKT7M; listed in **Table S1**). Taqman assays were performed in triplicate and normalized to VE2 levels.

### Capture-C data

Capture-C data were generated and analyzed as previously described ^48,49^. Briefly, regions for bait design were determined for each locus via identification of the set of SNPs with a LLR < 100 relative to the leading SNP of a given GWAS signal. For the 2q33.1 locus, capture baits were tiled across the entire region of association (chr2:202,114,359 to 202,205,025); baited fragments are listed in **Table S32**. Capture-C libraries were made using the Arima HiC Kit (Arima Genomics, Carlsbad, CA) and the KAPA HyperPrep Kit (KAPA Biosystems, Wilmington, MA) following the manufacturer’s protocol. Data are presented for region-specific Capture-C runs with five independent human primary melanocyte cultures (C56, C140, C205, C24, and C27) with three biological replicates for each. Paired-end sequencing reads from biological replicates were pre-processed with the HiCUP pipeline^82^ and aligned to human genome version 19 via bowtie2^83,84^. Chromatin interaction loops were detected at one- and four-fragment resolutions via Capture Hi-C Analysis of Genomic Organization (CHiCAGO) pipeline v.1.16.0^85^. Because we observed no distant interactions between fine-mapped variants and gene promoter at one-fragment resolution, data presented are from four-fragment analysis. Visualization was performed using the WashU Epigenome Browser^86^.

### Chromatin conformation capture (3C)

Chromatin conformation capture (3C) assays were done based on the protocol from the Dekker Lab^87^. RP11-1078D1 bacterial artificial chromosome (BAC) clones were purchased from BACPAC Resources Center and purified with the QIAGEN large-construct Maxi Kit to cover the 2q33.1 genomic region (chr2:202,045,328 to 202,245,875). BAC libraries were made by HindIII digestion of BAC plasmids, followed by ligation and DNA purification. To generate melanoma cell and melanocyte 3C libraries, we fixed and lysed 1 x 10^8^ cells, followed by HindIII digestion and ligation. Both the BAC libraries, and melanoma and melanocytes 3C libraries were amplified with a Taqman assay containing a primer localized to various regions of chr2:202,045,328 to 202,245,875 and a fixed primer harboring rs3769823, rs3769821, and rs59308963 plus a FAM-labeled probe annealing to 3’ of the fixed primer. Primers are listed in **Table S31**. PCR cycle conditions were as follows: 95°C, 5min; [95°C, 30s; 62°C, 30s; 72°C, 30s]_x34_; 72°C, 10min. Amplification for different primer pairs from the melanoma and melanocyte 3C libraries was normalized to that of BAC libraries.

### Epigenomic annotations

Chromatin Primary Core Marks Segmentation by HMM (ChromHMM), DNase Hypersensitivity (DHS), and histone modification data for primary human melanocyte cultures, primary human keratinocyte culture, and human breast mammary cultures “breast variant human mammary epithelial cells” and “breast myoepithelial primary cells” (shown as vHMEC and BMYO, respectively, in **Figure 1**) were obtained from the Roadmap Epigenomics Project (downloaded through the UCSC Genome Browser; http://genome.ucsc.edu/). Peaks from FAIRE-seq and H3K27ac ChIP for 11 melanoma cultures samples were obtained from the Gene Expression Omnibus (GEO; accession GSE60666)^45^.

### Cell culture

We originally obtained melanoma cell lines (UACC903 and UACC1113) from the University of Arizona Cancer Center (UACC), which were maintained in RPMI-1640 medium with L-glutamine (Quality Biological, 112-025-101) supplemented with 10% FBS (GenClone, 25-514H), 20 mM HEPES (pH 7.9; Gibco, 15630), and 100 U/ml penicillin-streptomycin (Gibco, 15140). Primary melanocytes from foreskin healthy newborn males, mainly of European descent, were grown in M254 media (Invitrogen, M254500) with HMGS-2 supplement (Invitrogen, S0165). MCF7 and T47D breast cancer cell lines were purchased from ATCC. MCF7 cells were grown in DMEM with L-glutamine (Quality Biological, 112-014-101) supplemented with 10% FBS, and 100 U/ml penicillin-streptomycin. T47D cells were maintained in RPMI 1640 supplemented with 10% FBS, and 100 U/ml penicillin-streptomycin. All cell lines were grown in a 5% CO_2_ humidified incubator at 37 °C and were tested for mycoplasma contamination every 3-6 months.

### Luciferase reporter assays

Sequences encompassing each variant were PCR-amplified (primers listed in **Table S31**) from genomic DNA of HapMap CEU panel samples with the appropriate genotypes to obtain clones with each genotype, and cloned into the HindIII and XhoI sites of the pGL4.23[*luc2*/minP] (Promega, Madison, MI) luciferase vector in the 5’-to-3’ or 3’-to’5’ orientation using In-Fusion HD Cloning Kit (Takara Bio). Sequence-verified pGL4.23 constructs were co-transfected with the pGL4.73[*hRluc*/SV40] Renilla luciferase control vector (Promega) into melanoma cell lines (UACC903 and UACC1113) using Lipofectamine^TM^ 2000 (Invitrogen) in 24-well format. Luciferase activity was measured 24 h after transfection with the Dual Luciferase Reporter Assay System (Promega). Luciferase activity was normalized to the Renilla luciferase activity. All the experiments were performed in six replicate wells and repeated for at least three biological replicates. Significance between alleles was assessed using a two-tailed, unpaired t-test assuming unequal variance.

### Massively parallel reporter assay data

We used data from two massively parallel reporter assay (MPRA) datasets assessing allelic *cis*-regulatory potential for candidate causal variants from this locus in melanoma cells and melanocytes^46,47^. Firstly, in a prior MPRA analysis (MPRA v.1)^46^, 823 variants from 20 genome-wide significant loci from a prior melanoma meta-analysis^8^ were tested in a melanoma cell line, UACC903, as well as HEK293FT cells. For this study, an LD-based selection criteria was used (r^2^ > 0.4 with the sentinel SNP using 1,000 Genomes phase 3 EUR data), further filtering candidate causal variants to test only those located within annotated open chromatin regions and promoter/enhancer histone marks found in primary melanocytes and/or short-term cultures. A second MPRA study (MPRA v.2)^47^ investigated 54 loci from a more recent melanoma GWAS meta-analysis^10^ in both melanoma cells and melanocytes. Here, 1,992 candidate causal variants were tested from 54 genome-wide significant loci. Candidate causal variants were fine-mapped using log-likelihood ratios (LLR < 1:1,000 with the lead SNP) or where variants were not successfully imputed or tested in the GWAS using an LD-based criteria (r^2^ > 0.8 with the lead SNP, 1,000 Genomes EUR).

### EMSAs and supershift assays

Nuclear extracts from melanoma cell lines (UACC903 and UACC1113) or human melanocytes were prepared using the NE-PER Nuclear and Cytoplasmic Extraction Kit (Thermo Fisher Scientific). Nuclear extract lysates for MCF7 and T47D breast cancer cell lines were purchased from Abcam (ab14860 and ab14896, respectively). Oligonucleotides for each SNP were synthesized with or without biotin labeling at the 5’ end and HPLC purified (21-28nt, Life Technologies, listed in **Table S31**). Forward and reverse strands were then annealed to generate double stranded 5’-end labelled probes or unlabeled competitors. 50 nM probes were added to 2 µg of nuclear extract preincubated with 1 µg of poly d(I-C) (Roche) in LightShift^TM^ 1 × binding buffer (Thermo Fisher Scientific) for 30 min on ice. Competition experiments were performed by adding 1-100 fold more unlabeled competitor oligonucleotides to the reaction mixture 5 min before the addition of labelled probes. Anti-E4F1 (Abcam, ab70615) or anti-IRF2 (Abcam, ab245658) antibodies for supershift assays were incubated with nuclear extract for 1 h at 4 °C prior to adding poly d(I-C). The reactions were run on 5 – 10% TBE gels (Bio-Rad Criterion) on ice at 120 V, transferred to Biodyne B membranes (VWR), transferred blots were crosslinked (Stratagene UV Stratalinker 1800) and detected using the LightShift Chemiluminescent EMSA Kit (Thermo Fisher Scientific) and imaged on Chemidoc Touch (Bio-Rad). One representative data is shown from three biological replicates.

### Quantitative mass spectrometry

MCF7 breast cancer cells were grown in RPMI 1640 (Gibco) supplemented with 10% FBS, 100 U/ml penicillin and 100 μg/ml streptomycin (Gibco). The same culture conditions were used for UACC903 melanoma cells with additional 20 mM HEPES (pH 7.9). Nuclear lysates for DNA affinity purifications were collected as described previously^88^. Oligonucleotide probes with 5’-biotinylation of the forward strand were ordered from Integrated DNA Technologies (**Table S31**). DNA pulldowns and on-bead trypsin digestion were performed in Eppendorf tubes as described previously^89^. Briefly, forward and reverse oligos were annealed with a 1.5X molar excess of the reverse strand. For each pulldown, annealed DNA oligos (500 pmol) were immobilized on 10 μl (20 μl slurry) Streptavidin-Sepharose beads (GE Healthcare, Chicago, IL). Immobilized DNA oligos were incubated with nuclear extracts (500 μg) from UACC903 and MCF7, and 10 μg of non-specific competitor DNA (5 μg polydIdC, 5 μg polydAdT). After washing away unbound proteins, beads were resuspended in elution buffer (2 M Urea, 100 mM TRIS (pH 8), 10LmM DTT), peptides were alkylated with 50 mM iodoacetamide, and on-bead digested with 0.25 μg trypsin. Tryptic peptides were labeled by stable isotope dimethyl labeling on StageTips, as described previously^89^. Samples were eluted from the StageTips and matching light and medium labeled samples were combined. Samples were loaded onto a 30 cm column (heated at 40 °C) packed in-house with 1.8 μm Reprosil-Pur C18-AQ (Dr Maisch) and eluted using a gradient from 9 to 32% Buffer B (80% acetonitrile, 0.1% formic acid) over 114 min at a flow rate of 250 nL/min using an Easy-nLC 1000 (Thermo Fisher Scientific). Peptides were sprayed directly into either a Orbitrap Fusion Tribrid mass spectrometer (Thermo Fisher Scientific) or a Q Exactive mass spectrometer (Thermo Fisher Scientific). The mass spectrometers were operated as described previously^46,89^. Thermo RAW files were analyzed with MaxQuant 1.6.0.1 by searching against the UniProt curated human proteome (released June 2017) with standard settings^90^. Protein ratios were normalized by median ratio shifting and used for outlier calling. An outlier cutoff of 1.5 inter-quartile ranges in two out of two biological replicates was used. rs3769823 and rs3769821 were assayed in both UACC903 and MCF7 cells, while rs59308963 was assayed only in UACC903.

### Chromatin immunoprecipitation and genotyping

Following the manufacturer’s instructions of the Active Motif ChIP-IT High-Sensitivity kit, actively growing melanoma cells (UACC903 and UACC1649) or C87 primary melanocytes were fixed with 1% formaldehyde when 80-90% confluent. Nuclei of 1.0 – 1.5 × 10^7^ cells were prepared and sheared with ME220 Focused-ultrasonicator (Covaris, Woburn, MA) for 20 min following the instructions. 10 – 30 µg sheared chromatin from 1.5 – 4.5 × 10^6^ cells were used for each immunoprecipitation reaction with anti-E4F1, anti-IRF2 or anti-IgG (Abcam, ab37415) following the instructions. Purified immunoprecipitated DNA or input DNA was analyzed by SYBR Green qPCR for enrichment of target sites using the primers listed in **Table S31**. ChIP DNA or input DNA from UACC1649 or C87 melanocyte cell lines (heterozygous for rs3769823) was assayed for genotyping rs3769823 by Taqman genotyping assay (Assay ID: C 25808407_20). All experiments were performed in triplicate (technical replicates) and repeated for at least three biological replicates.

### siRNA-mediated knockdown of E4F1 and IRF2

SMARTpool ON-TARGETplus Human siRNAs to *E4F1* (L-011847-00-0005), *IRF2* (L-011705-02-0005) and ON-TARGETplus Non-targeting controls (D-001810-01-05) were purchased from Dharmacon (Lafayette, CO). Transfections were carried out using Lipofectamine^TM^ RNAiMAX (Invitrogen) according to the manufacturer’s instructions. Cells were collected after 48h for RNA and protein extractions.

### qPCR

RNA was isolated using RNeasy Mini Column (Qiagen), which was always complemented by DNase treatment. cDNA was synthesized from the total RNA using iScript^TM^ Advanced cDNA Synthesis Kit (Bio-Rad). Gene expression levels were quantified by qPCR using Taqman assays for *E4F1* (Hs00231773_m1), *IRF2* (Hs01082884_m1), *CASP8* (Hs01018151_m1, Hs01018149, Hs01018160_m1, Hs04405665_m1 and Hs01022432_m1), and *GAPDH* (Hs99999905_m1) from Thermo Fisher Scientific. Gene expression levels of *E4F1*, *IRF2* and *CASP8* were normalized to *GAPDH*. Each experiment was performed in triplicate and replicated three times. Significance was assessed using a Student’s two-tailed T-test assuming unequal variances.

### Western blotting

Whole-cell extracts prepared in RIPA buffer with protease inhibitors (Complete Protease Inhibitor, Roche) or phosphatase inhibitors (PhosSTOP, Roche). Samples were quantified using DC Protein Assay (Bio-Rad) and electrophoresed on 4–12% Bis Tris Plus Bolt gels in MES buffer (Invitrogen). Proteins were transferred to nitrocellulose using an iBlot2 (Invitrogen). Blots were blocked with 5% non-fat dry milk in Tris-buffered saline with 0.1% Tween-20 (TBST). Primary and secondary antibodies were diluted in 1-2% milk in TBST, and all washes were performed with TBST. Blots were rinsed briefly with PBS before the addition of ECL Prime Western Blotting Detection Reagent (Amersham, UK). Images were captured on ChemiDoc™ Gel Imaging System (Bio-Rad). The antibodies of anti-E4F1 (Santa Cruz, sc-514718), anti-IRF2 (Santa Cruz, sc-101069,) and anti-β-actin (Santa Cruz, sc-47778) were used as primary antibodies, and goat anti-mouse HRP (Santa Cruz, sc-2005) for secondary antibodies.

### Analysis of melanocyte and melanoma expression data

Transcriptomic data were generated as described previously from human primary melanocytes^36^, TCGA-SKCM^36,41,42^, early passage melanoma cultures^43^, the Leeds Melanoma Cohort^91^, and GTEx tissues (v.7; https://gtexportal.org/home/index.html)^40^. In brief, RNA Sequencing by Expectation maximization (RSEM, version 1.2.31, http://deweylab.github.io/RSEM/) was used to quantify the gene expression followed by the quantile normalization to obtain the final RSEM for primary melanocytes, TCGA-SKCM, and GTEx tissues as previously described^36^. For background correction and quantile normalization, Robust Multi-array Average (RMA) algorithm with default settings (Affymetrix) was performed for melanoma cell lines^43^. A primary melanoma transcriptomic dataset from the Leeds Melanoma Cohort study on 703 patients was generated using the Illumina DASL Human HT12 v4 array platform, as previously described^91^ (European Genome-phenome Archive accession EGAS00001002922; https://ega-archive/org/studies/EGAS00001002922). 16 samples were removed in quality control (expression data from only one extraction per patient) leaving a cohort of 687 patients^91^. LUMI^92^ was used for background correction and quantile normalization as previously reported^92^. For GTEx (v.7) transcriptomic data, genes were selected based on expression thresholds of > 0.5 RSEM and ≥ 6 reads in at least 10 samples, and expression values for each gene were inverse quantile normalized to a standard normal distribution across samples and described previously^36^. We used R (version 4.1.0) for Pearson correlation to analyze the correlations between candidate target genes and allelic-specific binding transcription factors. The significance level was set at *P* < 0.05 and [slope] > 0.4 for all these tests. Using transcriptomic and genotype datasets from multiple tissues, we also ran multiple linear regression on R (version 4.1.0) to evaluate the influence of allelic-specific binding transcription factor or genetic variable of rs3769823 on the *CASP8*.

## Acknowledgements

This research was supported in part by the Intramural Research Program of the National Institutes of Health (NIH). The contributions of the NIH author(s) are considered Works of the United States Government. The findings and conclusions presented in this paper are those of the author(s) and do not necessarily reflect the views of the NIH or the U.S. Department of Health and Human Services. We would like to thank members at the National Cancer Institute’s Cancer Genomics Research Laboratory (CGR) for help with sequencing efforts and Rouf Banday from the National Cancer Institute for assistance with minigene assay design and data interpretation. This work has been supported by the Intramural Research Program (IRP) of the National Cancer Institute, US National Institutes of Health. We thank Jeremy Bravo Narula for assistance with the manuscript. We thank all the cohorts, funders, and investigators who contributed to the melanoma GWAS, as originally acknowledged by Landi, 23andMe, and colleagues^10^; data from this GWAS were used toward fine-mapping. JN receives funding from Horizon Europe 101136622. The Leeds Melanoma Cohort recruitment, follow up and data collection was supported by CRUK C588/A19167, C8216/A6129, C588/A10721 and NIH CA83115. Mark Iles is supported in part by the National Institute for Health and Care Research (NIHR) Leeds Biomedical Reearch Centre (BRC) (NIHR203331). The views expressed are those of the author(s) and not necessarily those of the NHS, the NIHR or the department of Health and Social Care.

## Declaration of interest

The authors declare no competing interests.

## Data and code availability

Conditional fine-mapping data are available in **Tables S1** and **S11**; Bayesian fine-mapping results are available in **Tables S5**-**S7**. Colocalization analyses are available are in **Tables S2, S3**, **S12**, **S13**, **S19**, and **S20**. Data from the 2020 melanoma GWAS meta-analysis performed by Landi and colleagues was obtained from dbGap (dbGap: phs001868.v1.p1), with the exclusion of self-reported data from 23andMe and UK Biobank. The full GWAS summary statistics for the 23andMe discovery dataset will be made available through 23andMe to qualified researchers under an agreement with 23andMe that protects the privacy of the 23andMe participants. Please visit the 23andMe website for more information and to apply to access the data. Summary data from the remaining self-reported cases are available from the corresponding authors of that manuscript (Matthew H Law; Mark M Iles; and Maria Teresa Landi). Melanocyte eQTL data and RNA-seq expression data from 106 individuals are available from dbGap (dbGap: phs001500.v1.p1), and pre-analyzed cis-eQTL data from GTEx (v.7) were downloaded from the GTEx portal. eQTL data from 59 early melanoma cell lines was available from NCBI Gene Expression Omnibus (accessions GSE78995 for expression data; GSE99193 for genotyping data). Leeds Melanoma Cohort data from 703 patients was available in the European Genome-phenome Archive accession EGAS00001002922. MPRA data are available in the NCBI Gene Expression Omnibus as a SuperSeries under the accession number GEO: GSE129250 and GSE210356. Capture-C data have been previously published^48,49^; interaction data including both called interactions as well as raw sequencing data are available through ArrayExpress (accession: E-MTAB-15079). Specifically, for the 2q33.1 locus restriction fragments baited for region-specific Capture-C assays are provided in **Table S32**, and loops called by ChICAGO are in **Tables S23** and **S24**. Pearson correlations are available in **Table S25**, **S29** and **S30**. Multiple linear regression results are available in **Tables S26**-**S28**. Luciferase assay fragments, other primers and Taqman assays are listed in **Table S31**.

## Web resources

23andMe, https://research.23andme.com/collaborate/#dataset-access/

GTEx portal, https://www.gtexportal.org/home/

https://ldlink.nci.nih.gov/?tab=home

NCBI GEO, https://www.ncbi.nlm.nih.gov/geo/

RegulomeDB, https://regulomedb.org/regulome-search/

The Cancer Genome Atlas (TCGA) Research Network, https://www.cancer.gov/about-nci/organization/ccg/research/structural-genomics/tcga

## Notes

### Competing Interest Statement

The authors have declared no competing interest.

### Summary of Updates

Submission has been updated to add Supplementary Figures and Supplementary Tables

