## Supplementary Figures for "Functional characterization of a multi-cancer risk locus on chromosome band 2q33.1 near *CASP8*"

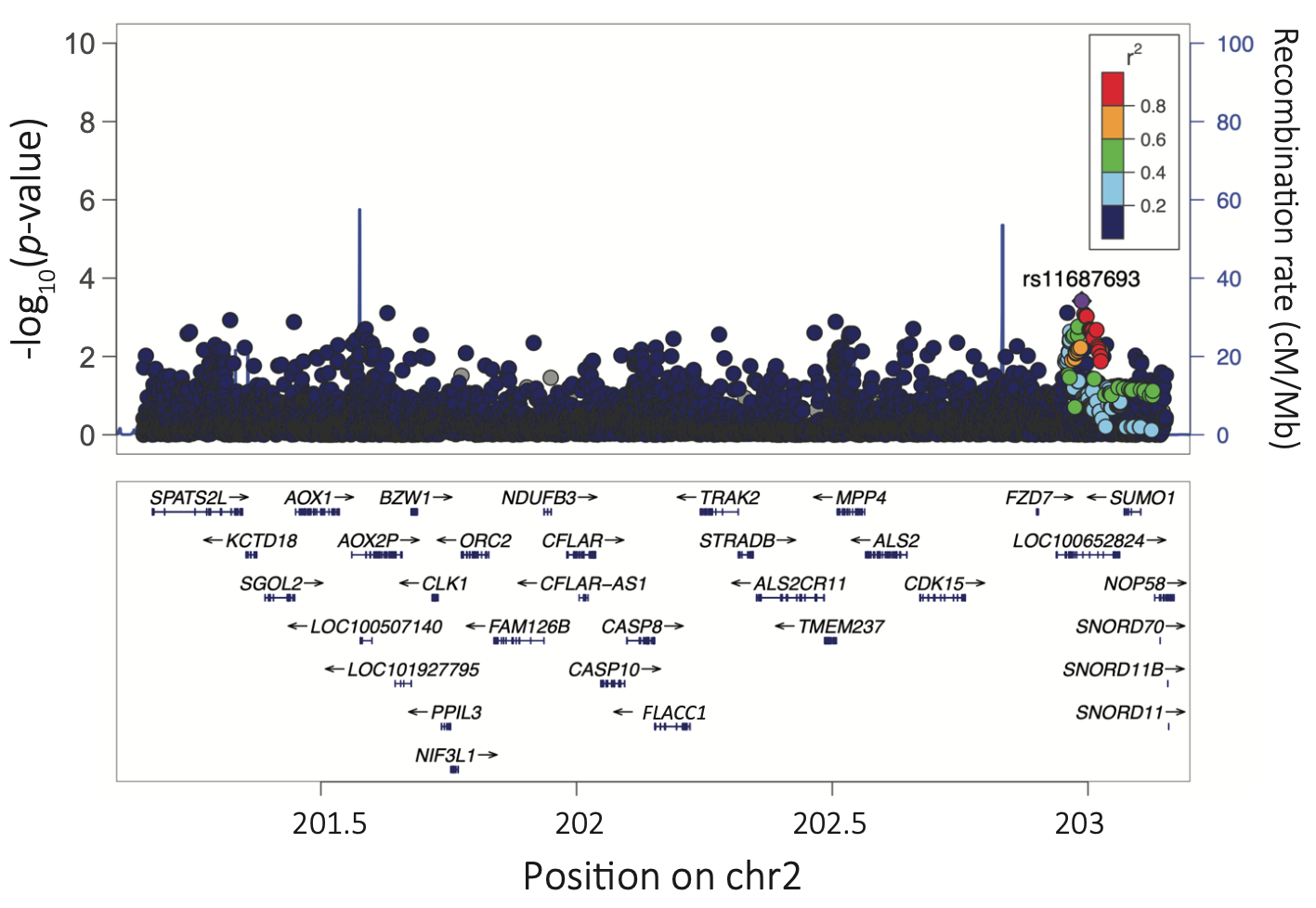

**Supplementary Figure S1. Manhattan plot of conditional association signals from the melanoma GWAS meta-analysis for the locus on chromosome band 2q33.1.** Negative log_10_ melanoma GWAS *P*-values are shown for variants at 2q33.1 when conditioned on the melanoma lead SNP rs10931936. The most significant signals are rs563855920 (*P*_conditional_ = 3.05 x 10^-4^, not labeled) and rs11687693 (*P*_conditional_ = 3.83 x 10^-4^, r^2^ = 0.0011 with rs10931936, labeled and highlighted in purple). LD relative to rs11687693 (r^2^, based on 1000G EUR) is color coded for all other variants. Genomic coordinates are based on hg19.

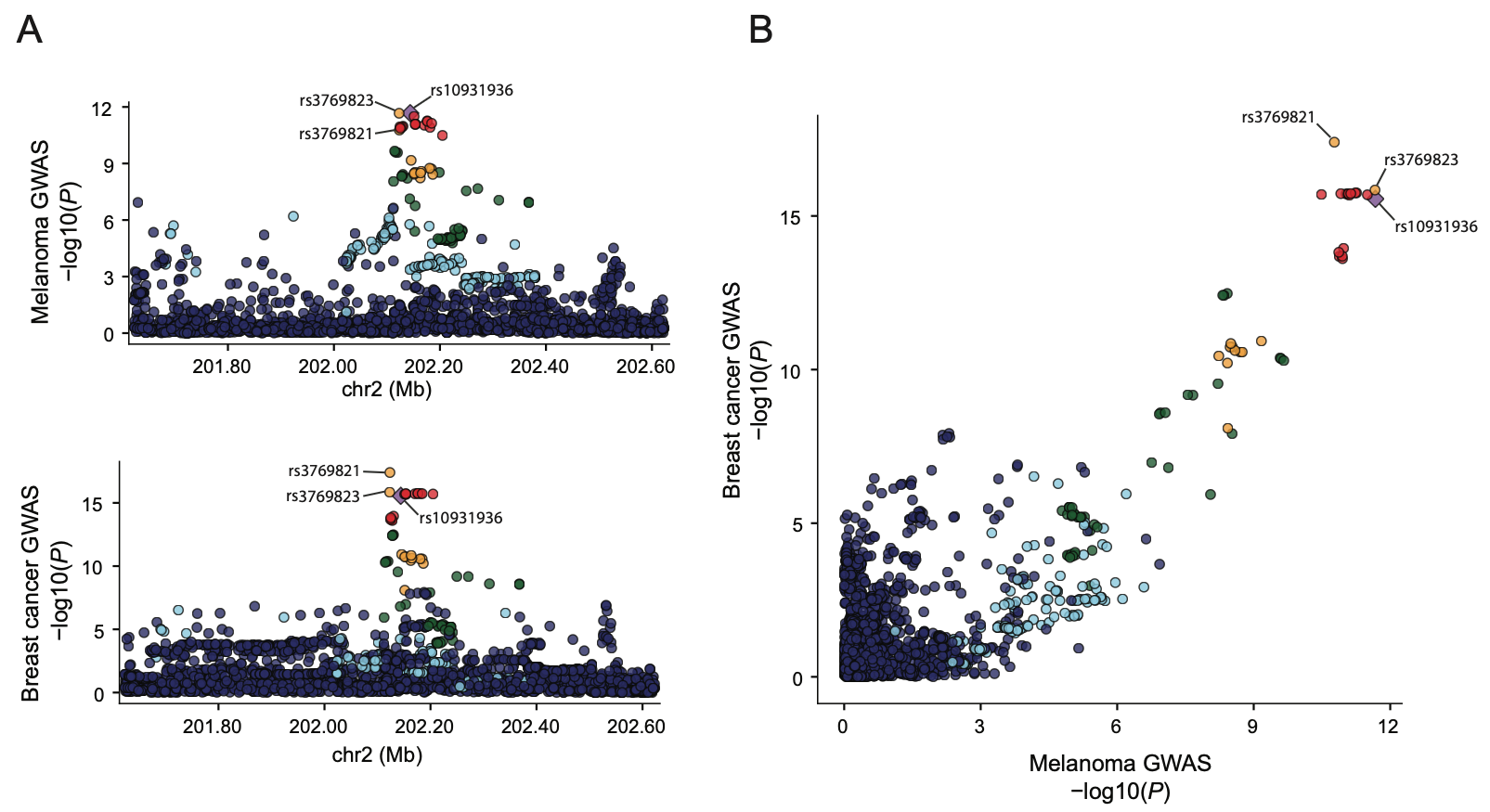

**Supplementary Figure S2. The GWAS signal in the melanoma for 2q33.1 locus colocalizes with the breast cancer GWAS signal.** (A) LocusZoom and (B) LocusCompare plots comparing *P*-values from melanoma and breast cancer GWAS for all commonly tested variants within a 1 Mb region encompassing rs10931936. The melanoma risk lead SNP rs10931936 is labeled and highlighted in purple, and LD (r^2^, based on 1000G EUR) of all other variants to the melanoma GWAS lead SNP is color-coded. The breast cancer GWAS lead SNP is rs3769821. Genomic coordinates are based on hg19.

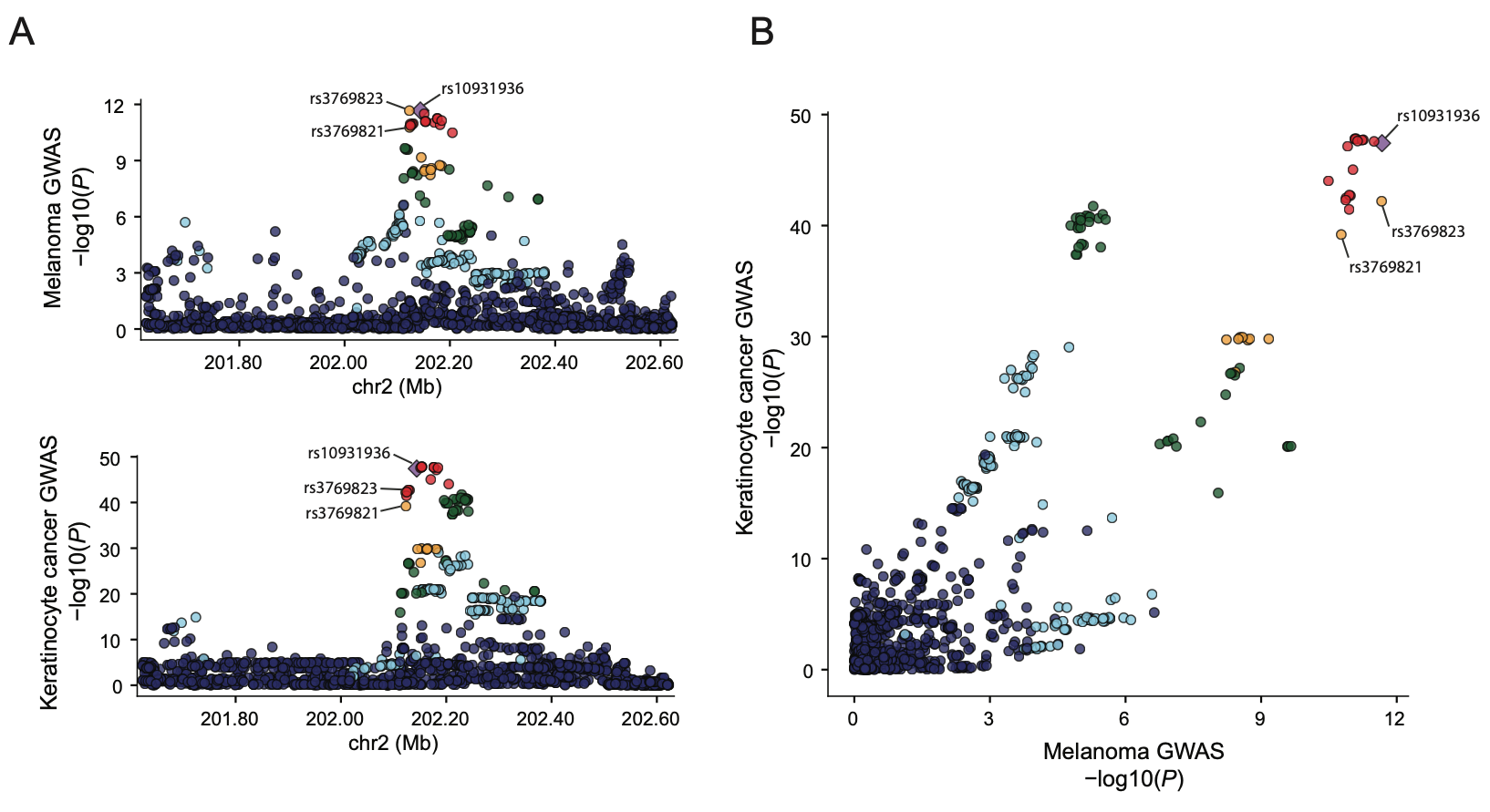

**Supplementary Figure S3. LocusCompare visualization of colocalization between the melanoma GWAS and keratinocyte cancer GWAS.** (A) LocusZoom and (B) LocusCompare plots comparing *P*-values from melanoma and keratinocyte cancer GWAS for all commonly tested variants within a 1 Mb region encompassing rs10931936. The melanoma GWAS lead SNP rs10931936 at 2q33.1 is labeled and highlighted in purple, and LD (r^2^, based on 1000G EUR) of all other SNPs to the GWAS lead SNP is color-coded. rs10931936 is the keratinocyte cancer risk lead SNP. Genomic coordinates are based on hg19.

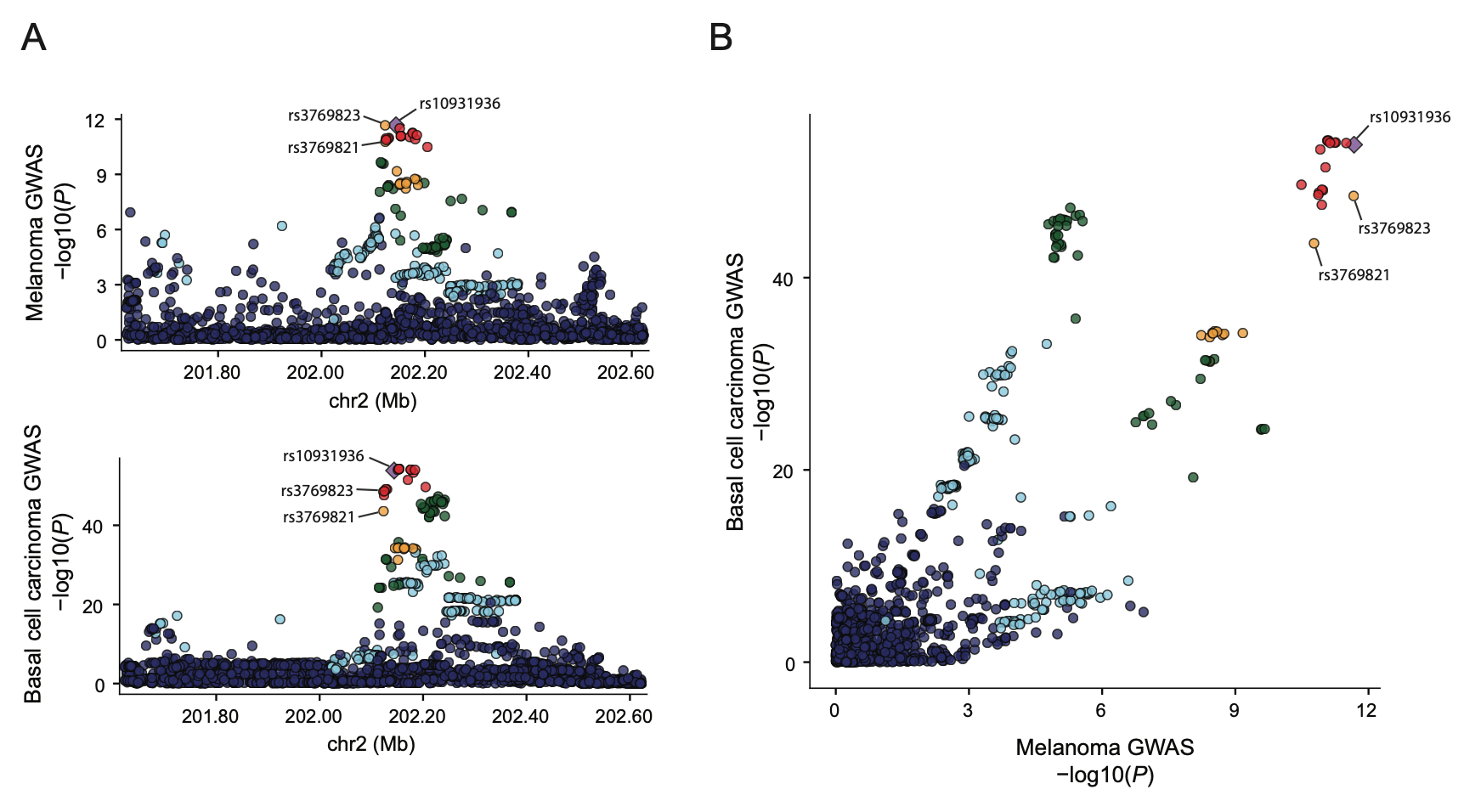

**Supplementary Figure S4. LocusCompare visualization of colocalization between the melanoma GWAS and cutaneous BCC GWAS.** (A) LocusZoom and (B) LocusCompare plots comparing *P*-values from melanoma and cutaneous BCC GWAS for all commonly tested variants within a 1 Mb region encompassing rs10931936. The melanoma GWAS lead SNP rs10931936 at 2q33.1 is labeled and highlighted in purple, and LD (r^2^, based on 1000G EUR) of all other SNPs to the GWAS lead SNP is color-coded. rs10931936 is the cutaneous BCC risk lead SNP. Genomic coordinates are based on hg19.

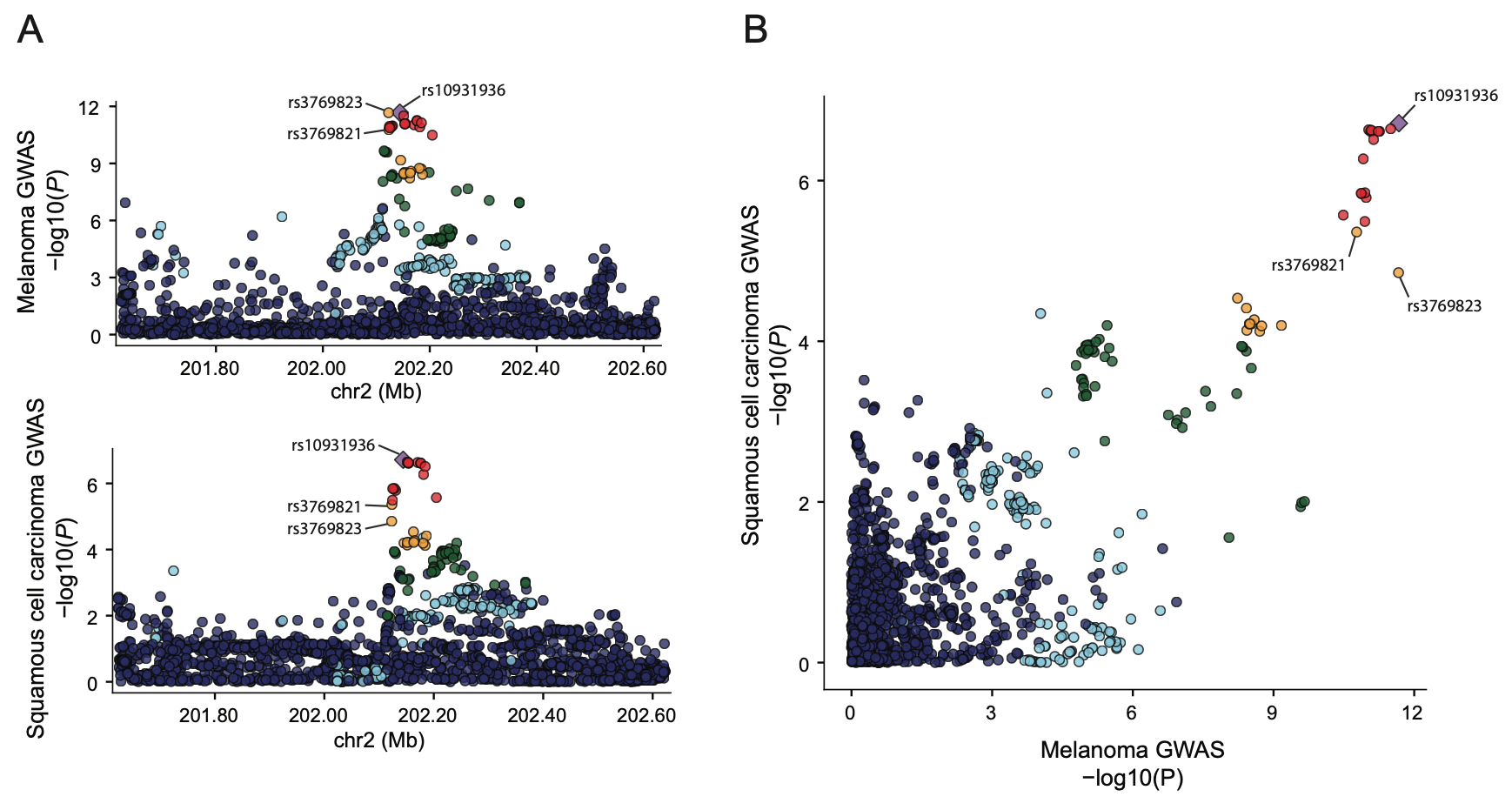

**Supplementary Figure S5. LocusCompare visualization of colocalization between the melanoma GWAS and cutaneous SCC GWAS.** (A) LocusZoom and (B) LocusCompare plots comparing *P*-values from melanoma and cutaneous SCC GWAS for all commonly tested variants within a 1 Mb region encompassing rs10931936. The melanoma GWAS lead SNP rs10931936 at 2q33.1 is labeled and highlighted in purple, and LD (r^2^, based on 1000G EUR) of all other SNPs to the GWAS lead SNP is color-coded. rs10931936 is the cutaneous SCC risk lead SNP. Genomic coordinates are based on hg19.

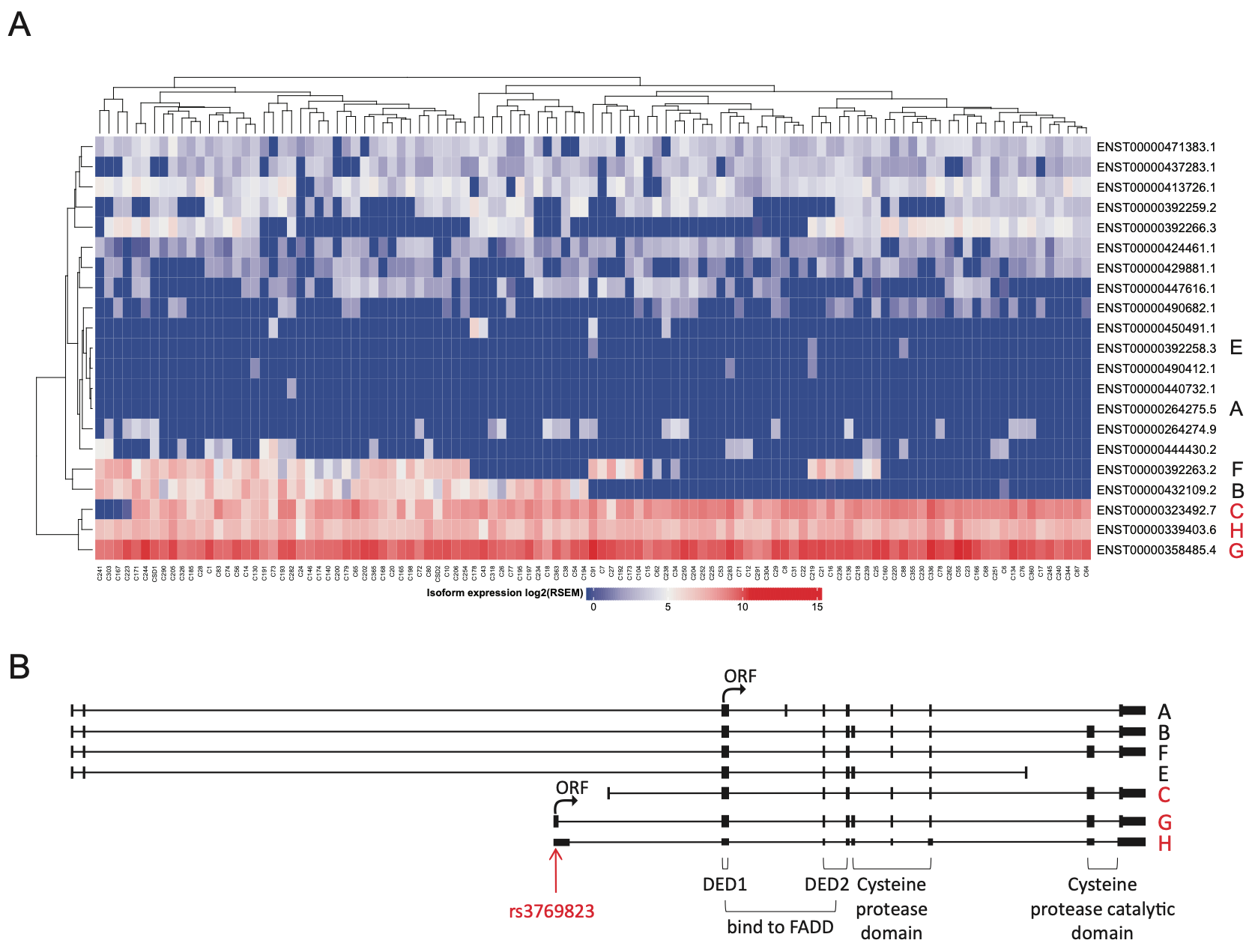

**Supplementary Figure S6. *CASP8* transcript variants expressed in human primary melanocytes and the transcript structure of *CASP8*.** (A) Heatmap of RNA sequencing data for clustered *CASP8* transcript isoform expression levels from 106 human primary melanocyte cultures. RNA sequencing data was performed using RSEM expression quantification package (iterations of Expectation-Maximization algorithms to assign reads to the isoforms from which they originate). Genotypes for *CASP8* splice QTL-associated variants are shown. (B) A schematic representation of *CASP8* gene structure is shown with protein domains annotated. ORF indicates translation start region. CASP8 can be divided into three domains; death effector domain (DED) at the N-terminal is required for Fas-associated death domain (FADD) binding, cysteine protease domain and catalytic domain, which contains two CASP8 cleavage sites and allows release of active CASP8 enzyme.

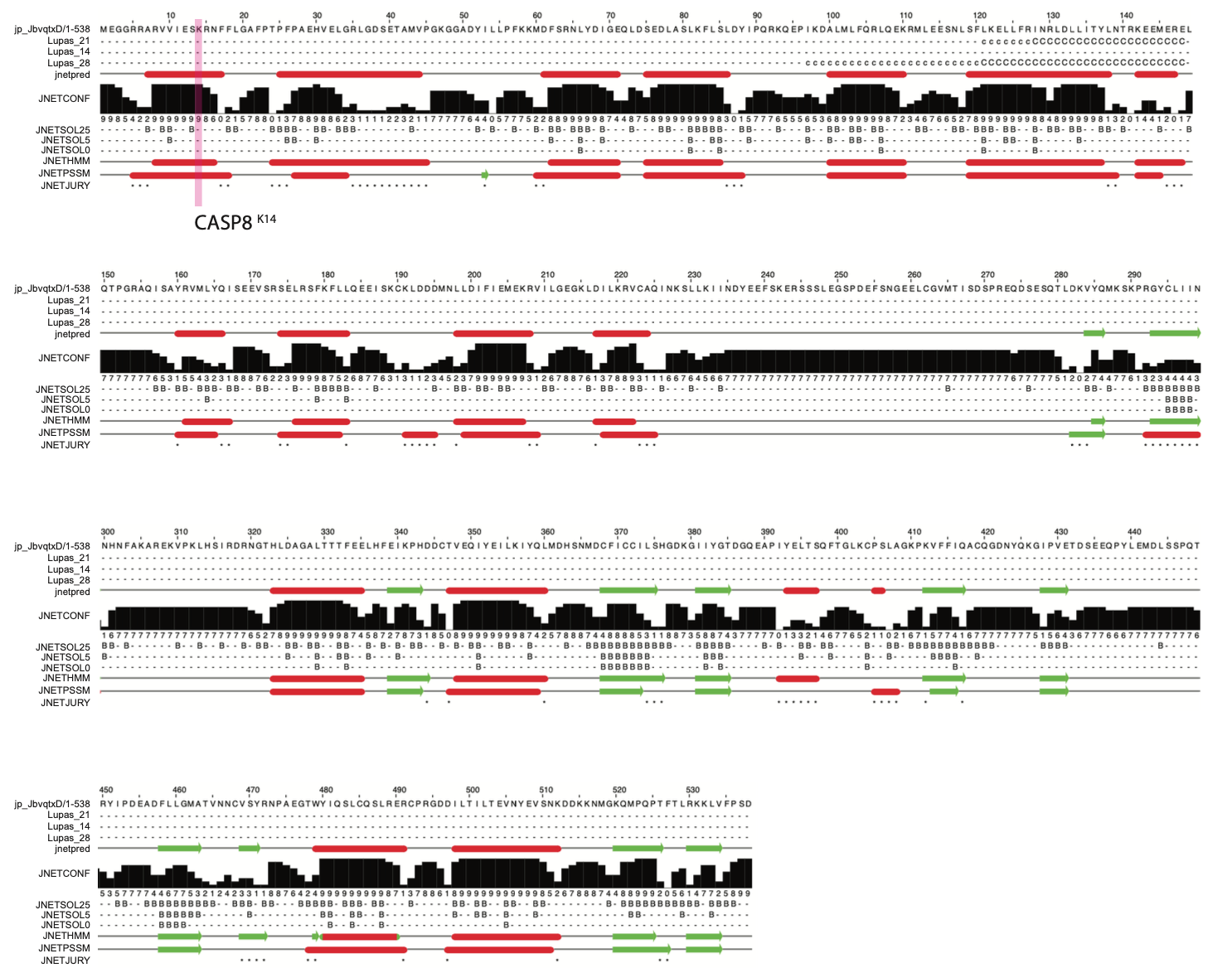

**Supplementary Figure S7. Prediction of the possible impact of rs3769823, CASP8^K14R^ on the structure and function of a human protein with Jpred4.** A view of the CASP8 secondary structure based on amino acid sequence. Lupas coiled-coil predictions for the sequence (window size of 14, 21, and 28; - = less than 50% probability; c = between 50% and 90 % probability; C = greater than 90% probability) are shown. Jnetpred is the consensus prediction (helices are marked as red tubes, and sheets as dark green arrows). JNETCONF is the confidence estimate for the prediction (high values mean high confidence). JNETSOL is the Jnet prediction of solvent accessibility (25, less than 25% solvent accessibility; 5, less than 5% exposure; 0, 0% exposure). JNET-HMM (Hidden Markov Model) and JNET-PSSM (Position-Specific Scoring Matrix) are probabilistic prediction models of a multiple sequence alignment of proteins (helices are marked as red tubes, and sheets as dark green arrows). A ‘*’ in this annotation indicates that the JNETJURY was invoked to rationalize significantly different primary predictions.

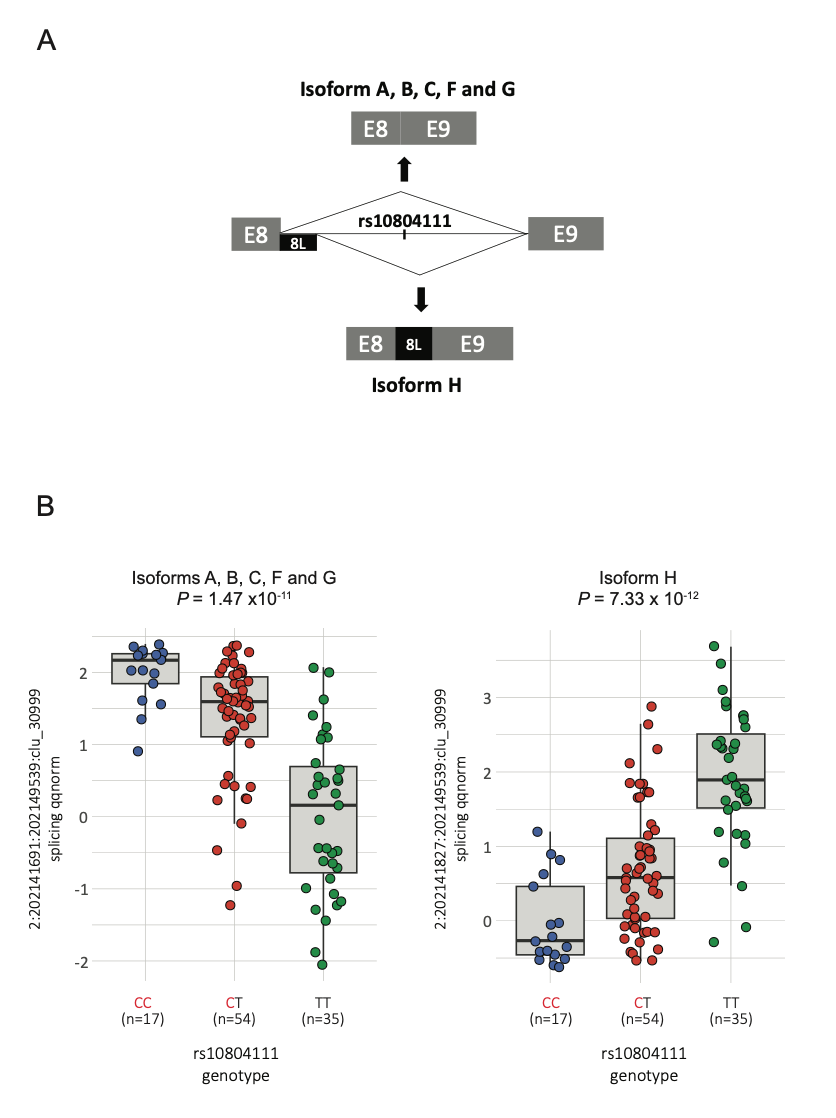

**Supplementary Figure S8. Melanocyte sQTL variant rs10804111 is associated with alternative splicing of *CASP8*.** (A) Schematic view of *CASP8* exons and splicing junctions. Melanoma GWAS variant rs10804111 is associated with alternative splicing events in the exon 8 to 9 junction. Alternative splicing generates isoform H retaining part of intron 8; ‘8L’ refers this isoform. (B) sQTL plots of *CASP8* junction usage as assessed by LeafCutter in RNA sequencing data from 106 human primary melanocyte cultures in relation to rs10804111 genotype. The *y*-axis displays standardized and normalized “percent spliced in” (ΔPSI) of the junction, ch2:202,141,691-202,149,539 or chr2:202,141,827-202,149,539, within the cluster 30999 (clu_30999) of introns sharing splice sites. sQTL nominal *P*-values and slope were derived from linear regression with no multiple-testing correction applied. C is the risk-associated allele, and T is the protective allele at rs10804111.

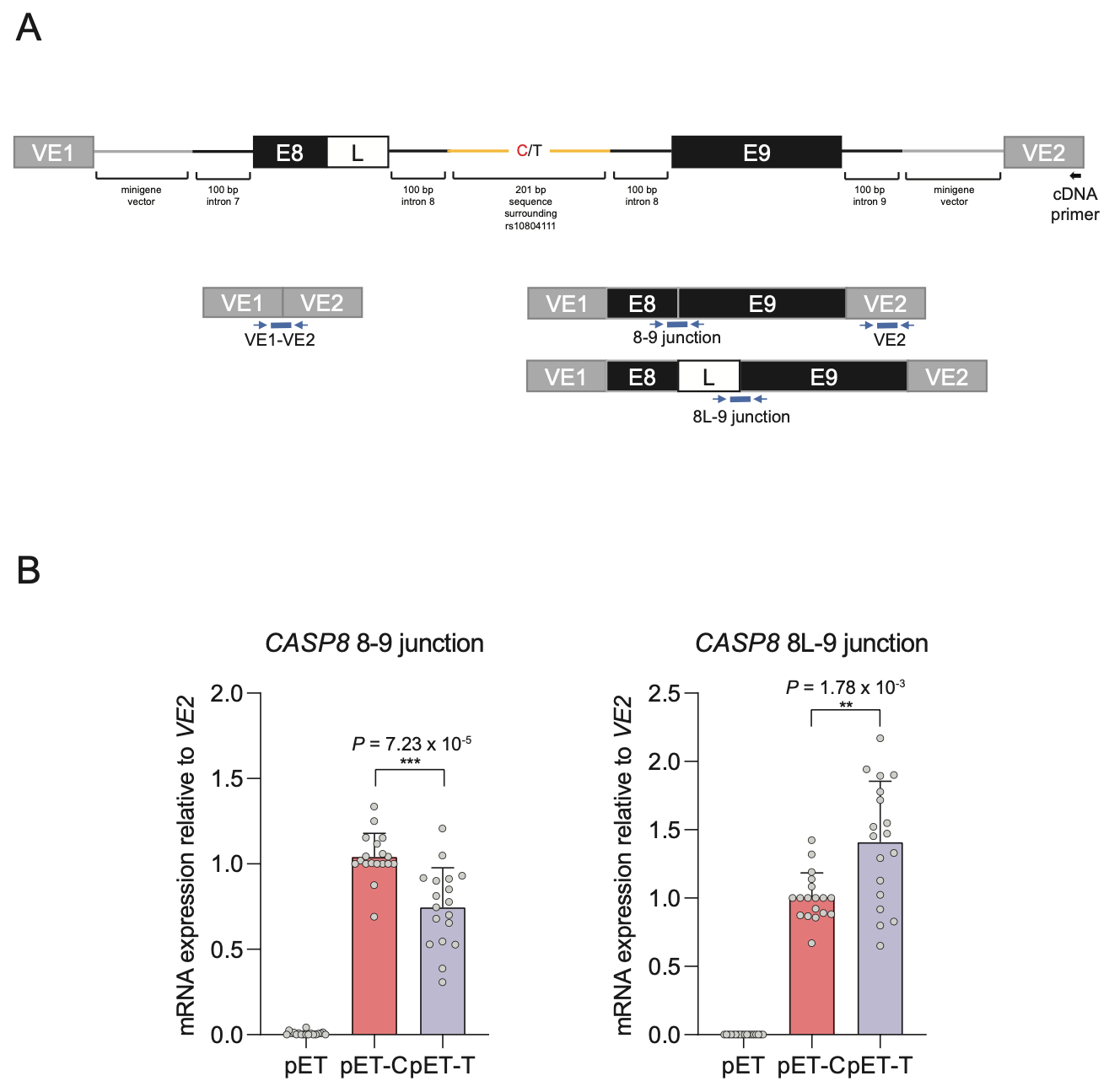

**Supplementary Figure S9. Splicing reporter minigene assay for the analysis of the effect of rs10804111 on alternative *CASP8* splicing.** (A) Minigene vectors were constructed to contain *CASP8* exon 8/8L and surrounding intron (100 bp upstream and 100 bp downstream intronic sequence) and *CASP*8 exon 9 and surrounding intronic sequence (100 bp upstream and 100 bp downstream intronic sequence), linked together with 201 bp of intronic sequence surrounding each allele of rs10804111. Two alternatively spliced minigene transcripts containing *CASP8* exons 8 and 9 can be produced with or without partial intron 8 retention. (B) Minigene vectors were transfected into UACC903 melanoma cells and 24h later relative levels of transcript harboring exon 8/9 and 8L/9 junctions were quantitated by qRT-PCR using fluorogenic probes designed to span specific intron/exon junctions. Transcript junction usage was normalized to exon VE2 mRNA levels. A paired two-tailed t-test assuming unequal variance was used to calculate all *P*-values shown for pET-C versus pET-T. C is the risk allele, and T is the protective allele; pET is empty minigene vector without *CASP8* intronic/exonic sequence. Data and *P*-values are from six replicate experiments combined.

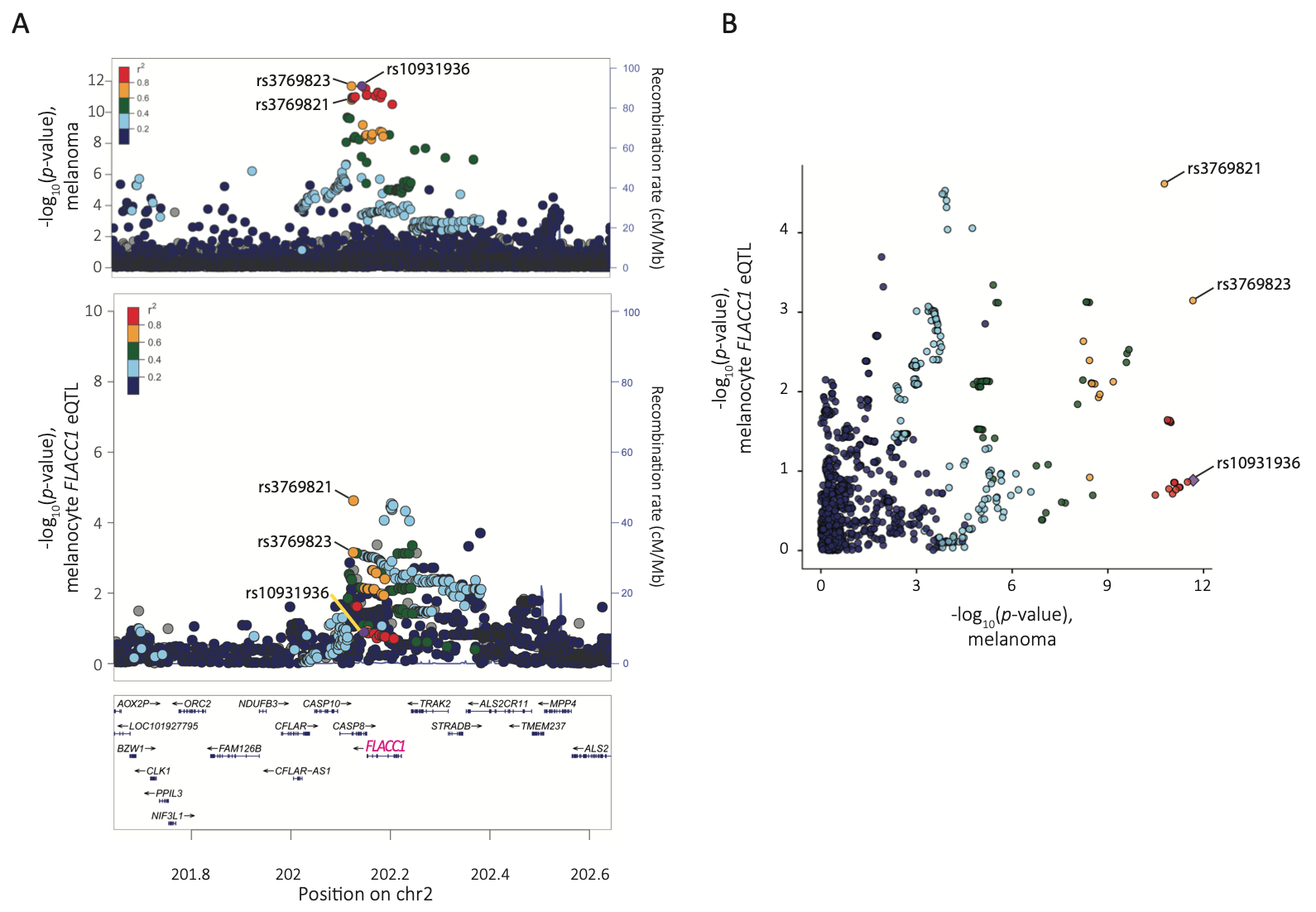

**Supplementary Figure S10. LocusCompare visualization of colocalization between the melanoma GWAS and the *FLACC1* eQTL signals in primary melanocytes.** (A) LocusZoom and (B) LocusCompare plots comparing *P*-values between melanoma GWAS and the *FLACC1* melanocyte eQTL for commonly tested variants within a 1 Mb region encompassing rs10931936. The melanoma risk lead SNP rs10931936 is labeled and highlighted in purple, and LD (r^2^, based on 1000G EUR) of all other SNPs to the melanoma lead SNP is color-coded. The best melanocyte *FLACC1* eQTL variant (rs3769821) is labeled. Genomic coordinates are based on hg19.

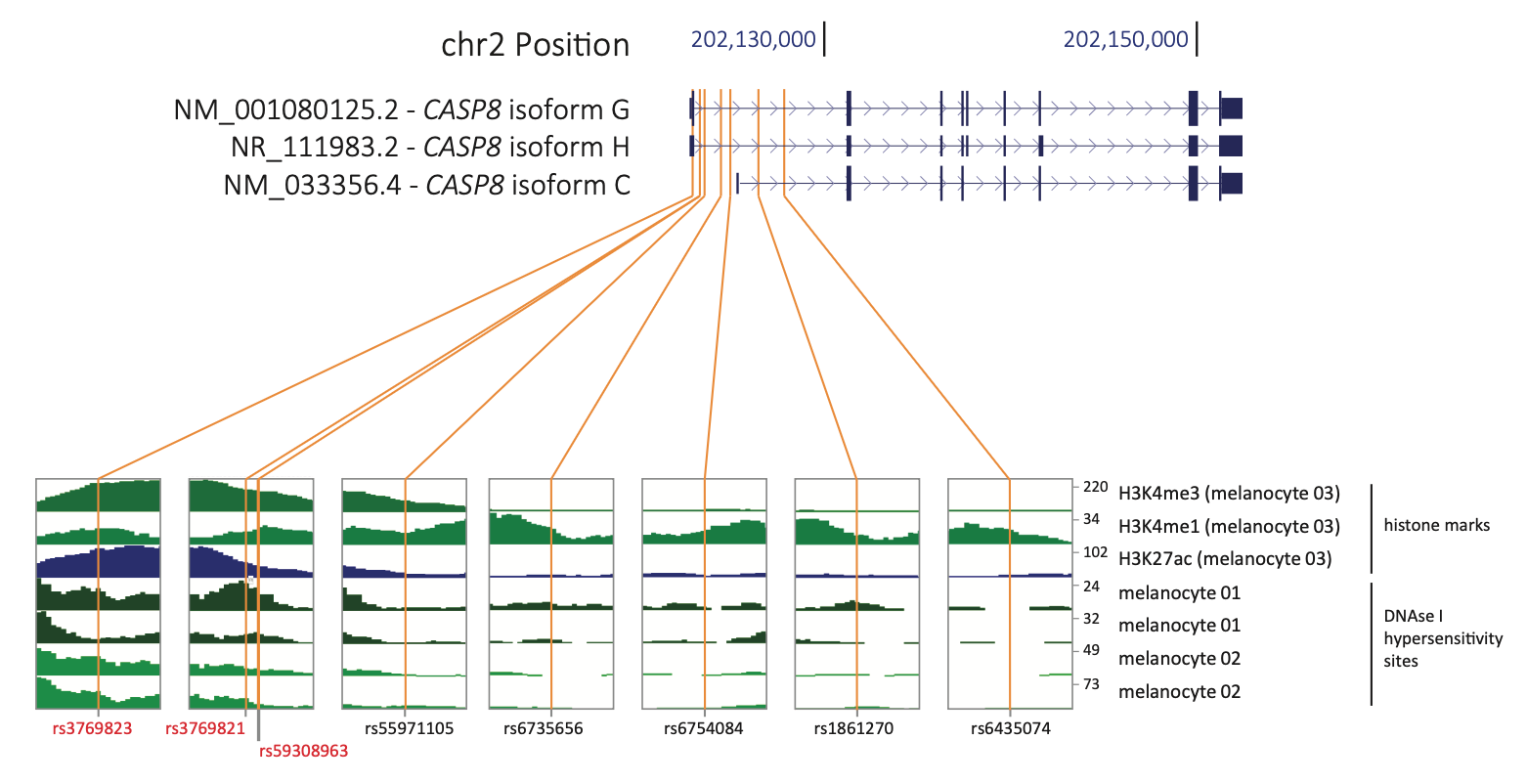

**Supplementary Figure S11. Functional annotation and prioritization of potentially *cis*-regulatory candidate sequence variants based on melanocyte-specific epigenomic data.** Prioritization of candidate causal variants was based on multiple fine-mapping approaches (LLR, Bayesian fine-mapping using DAP-G and PAINTOR, and LD r^2^), as well as location within potentially *cis*-regulatory regions in human melanocytes. All tracks for melanocyte DNaseI Hypersensitivity Site (DHS) and histone mark data were obtained from Roadmap Epigenome Projects through UCSC genome browser. Histone mark ChIP-seq signals (H3K4me3, H3K4me1, and H3K27ac) are shown for a representative individual melanocyte culture (melanocyte 03) among three analyzed. DNaseI hypersensitivity signals are shown for two melanocyte cultures (melanocyte 01, melanocyte 02). The scale of each track is uniformly set throughout the region of the *CASP8* gene to cover the highest peaks, with 0 as the baseline. Regions spanning 250 bp upstream and downstream of each candidate variant are enlarged in boxes. Genomic coordinates are based on hg19.

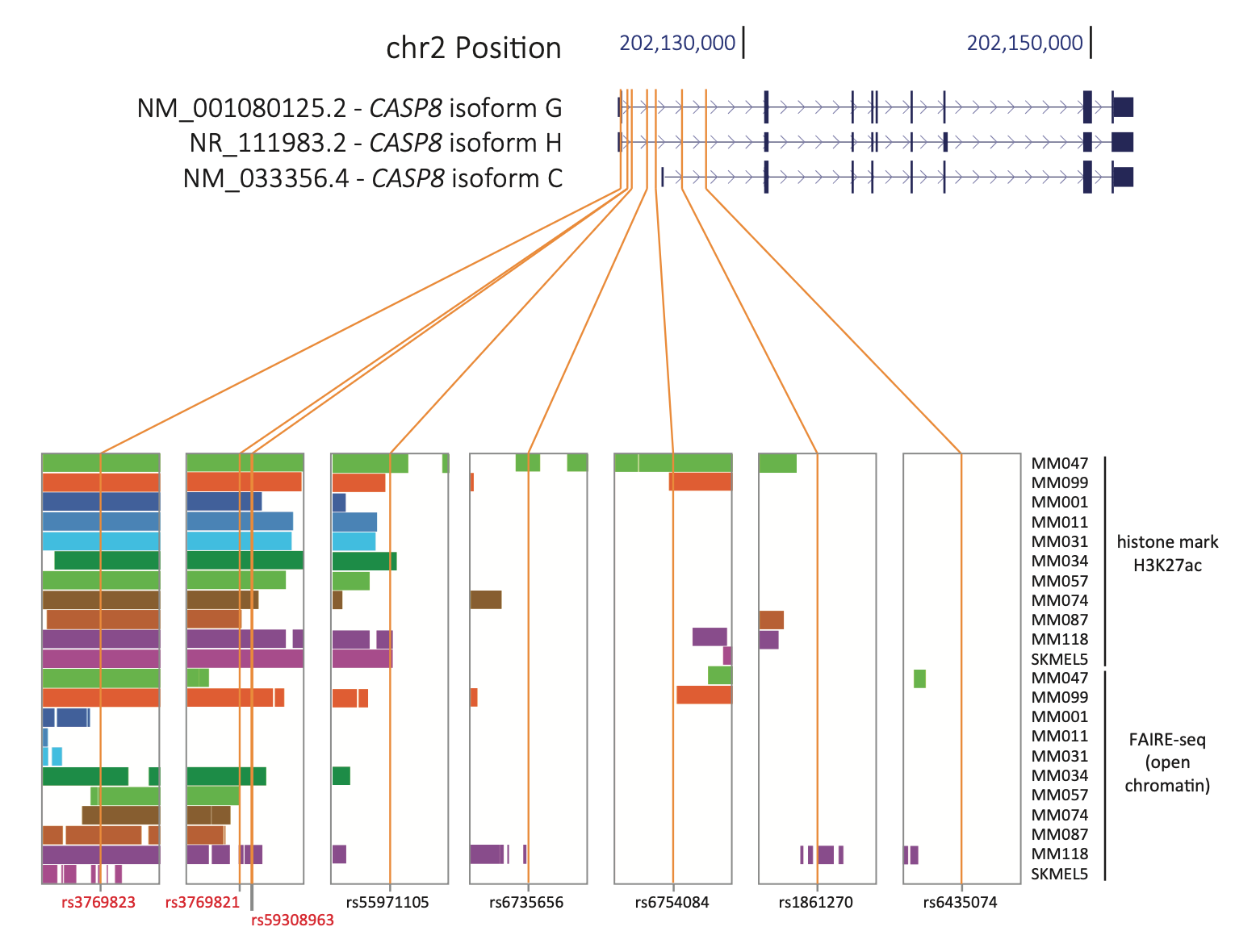

**Supplementary Figure S12. Functional annotation and prioritization of potentially *cis*-regulatory candidate sequence variants at the *CASP8* locus based on melanoma epigenomic data.** Prioritization of candidate causal variants was based on multiple fine-mapping approaches (LLR, Bayesian fine-mapping using DAP-G and PAINTOR, and LD r^2^) and both H3K27ac histone marks and open chromatin data (FAIRE-seq) derived from 11 melanoma cell cultures. All tracks were obtained from Melanoma Epigenome Project through the USCS genome browser. Peaks from H3K27ac ChIP-seq and FAIRE-seq are presented as interval of peaks. Regions spanning 250 bp upstream and downstream of each candidate variant are enlarged. Genomic coordinates are based on hg19.

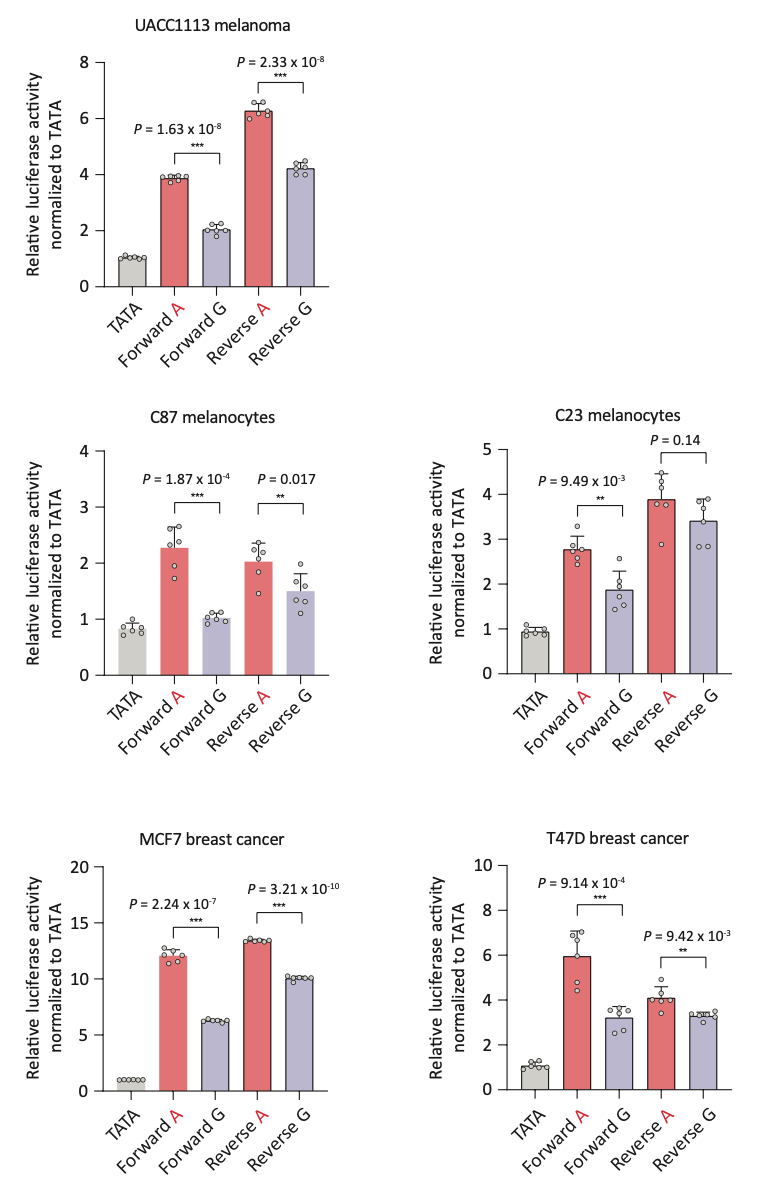

**Supplementary Figure S13. The melanoma-associated variant rs3769823 drives allelic-specific transcriptional activity in multiple cell lines.** Individual luciferase reporter assays for rs3769823 were conducted using the melanoma cell line UACC1113, two primary melanocyte cultures (C87 and C23), and two breast cancer cell lines (MCF7 and T47D). 138 bp regions encompassing rs3769823 were cloned 5’ of a minimal TATA promoter in the pGL4.23 vector, and cells were transfected. Luciferase activity was measured 24 h after transfection and was normalized against *Renilla* luciferase activity. Individual *P-*values are presented for A-risk allele versus G-protective allele (two-tailed, unpaired t-test assuming unequal variances). Data and *P*-values shown are from one representative experiment (*n*=6) from three biological replicates. TATA, minimal promoter control; A, risk allele construct; G, protective allele construct.

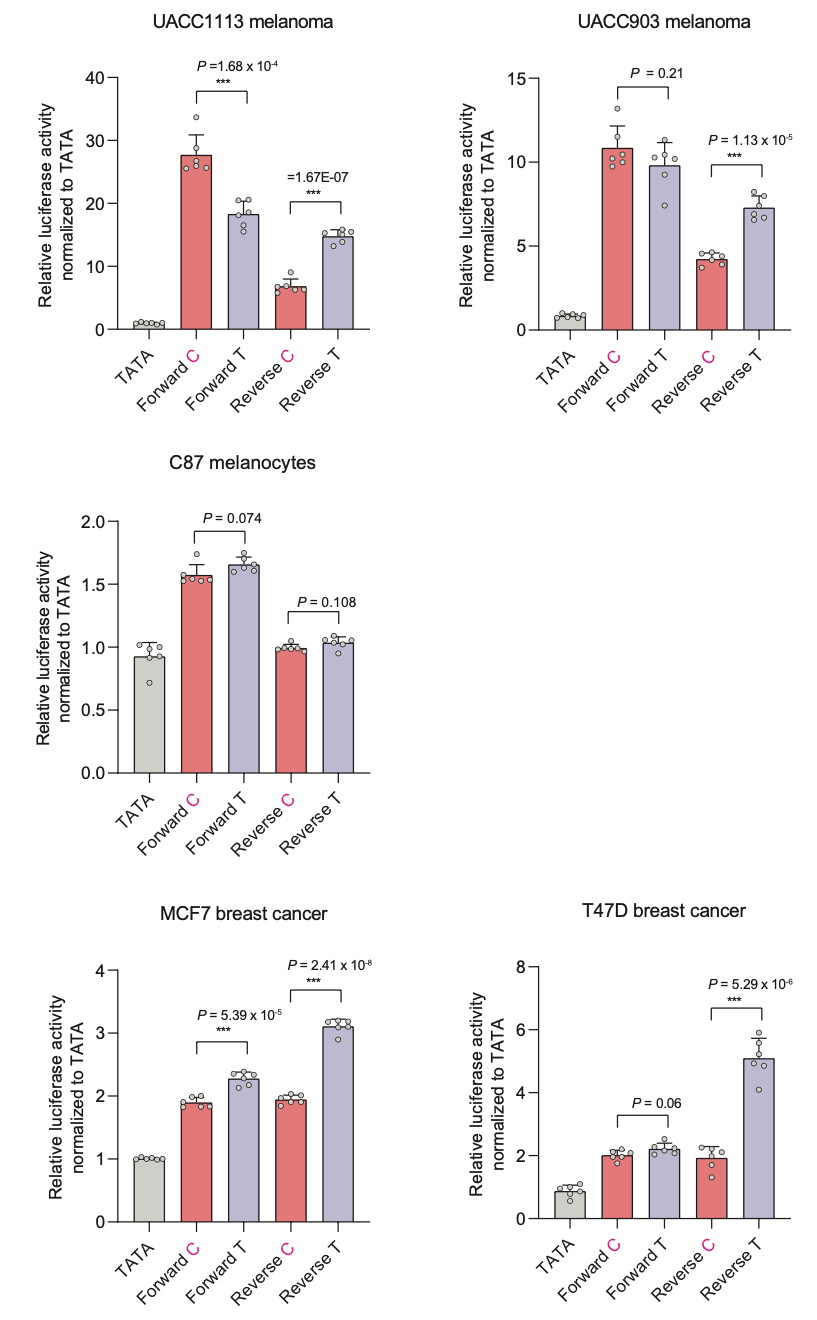

**Supplementary Figure S14. The melanoma-associated variant rs3769821 shows allelic-specific transcription activity in multiple cell lines.** Sequences containing rs3769821 (128 bp) were cloned 5’ of a minimal TATA promoter in the pGL4.23 vector and transfected into melanoma cell lines (UACC903 and UACC1113), C87 primary melanocytes, or breast cancer cell lines (MCF7 and T47D). Individual luciferase activity was measured 24 h after transfection and was normalized against *Renilla* luciferase activity. Individual *P*-values are shown for C-risk allele versus T-protective allele (two-tailed, unpaired t-test assuming unequal variances). One representative set (*n*=6) is shown from three biological replicates. TATA, minimal promoter control; C, risk allele construct in red; T, protective allele construct.

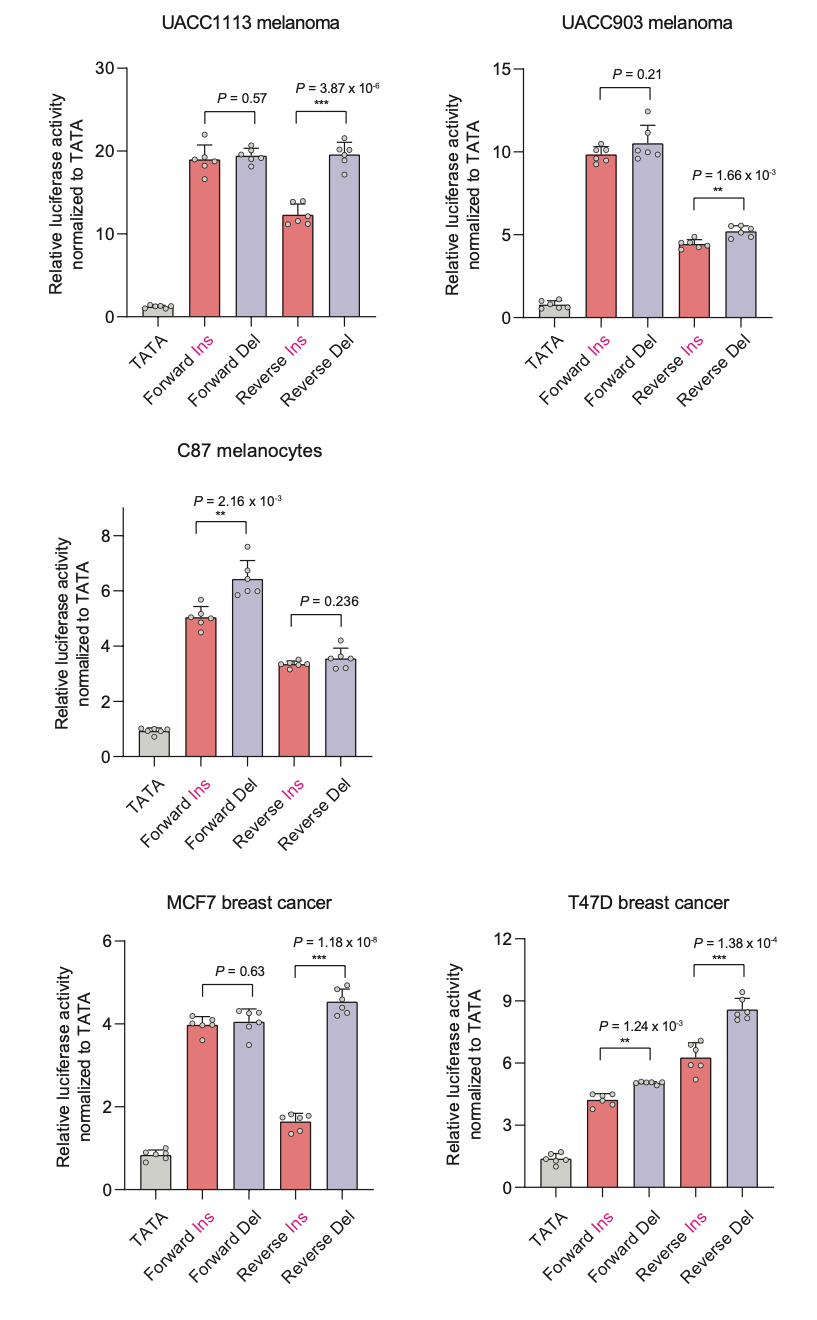

**Supplementary Figure S15. Allelic-specific transcriptional activity of the melanoma-associated variant rs59308963.** Sequences containing rs59308963 (117 and 125 bp) were cloned 5’ of a minimal TATA promoter in the pGL4.23 vector, and transfected into melanoma cell lines (UACC903 and UACC1113), C87 primary melanocytes, or breast cancer cell lines (MCF7 and T47D). Individual reporter activity was measured 24 h after transfection and was normalized against *Renilla* luciferase activity. Individual *P-*values are presented for insertion-risk (Ins) versus deletion-protective (Del) alleles (two-tailed, unpaired t-test assuming unequal variances). Data and *P*-values are from one representative set (*n*=6) from three biological replicates. TATA, minimal promoter control; Ins, risk allele construct in red; Del, protective allele construct.

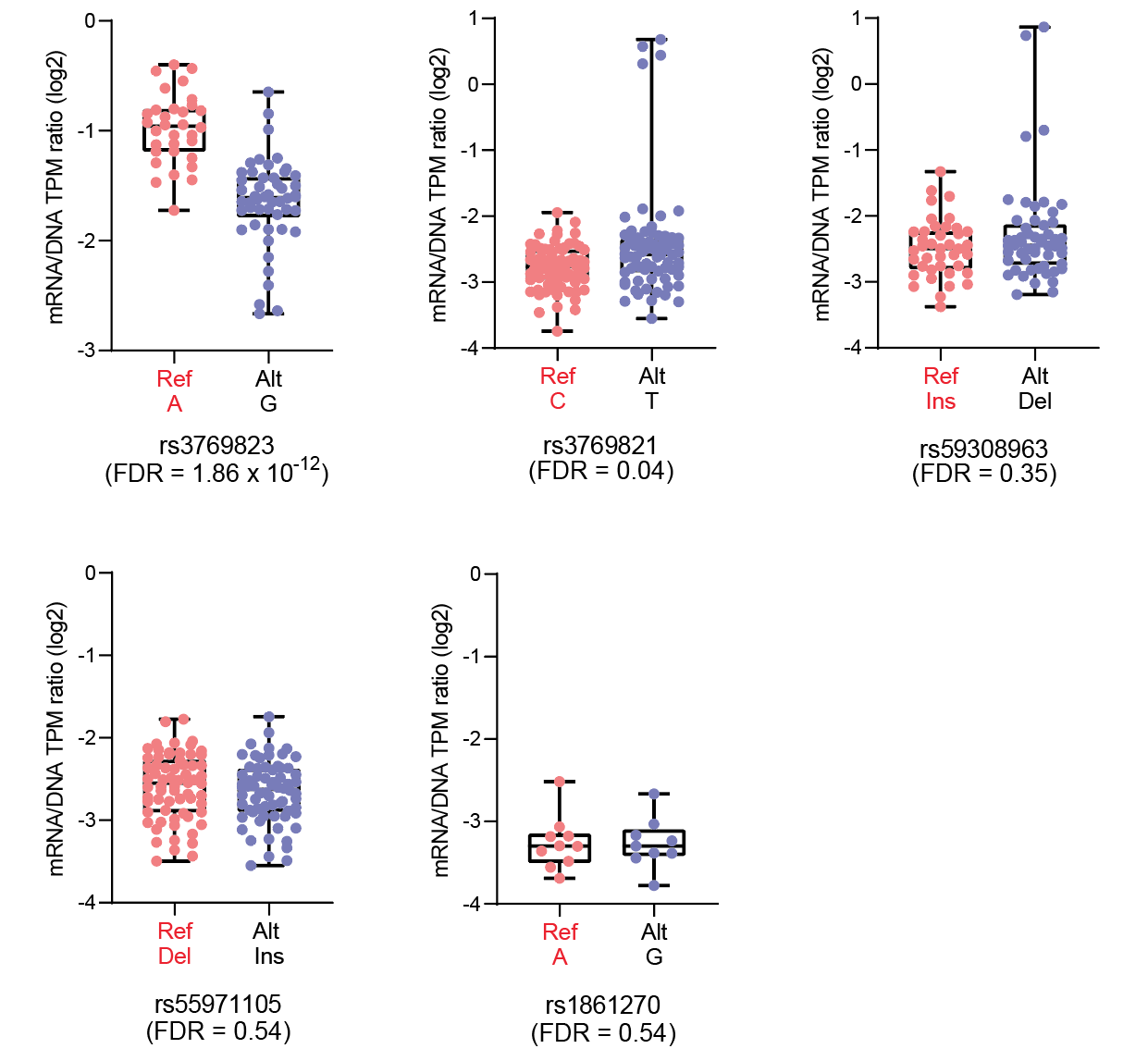

**Supplementary Figure S16. Massively Parallel Reporter Assay data for candidate causal variants at 2q33.1.** Transcriptional activity of 145 bp sequences of five candidate variants from a previously-published MPRA study are shown as normalized tag counts (log2 mRNA TPM/DNA TPM). Results from UACC903 melanoma cells are shown for both alleles in forward and reverse directions, where results from promoter and enhancer constructs were combined. The risk allele of each variant marked in red. FDR < 0.01 is considered significant.

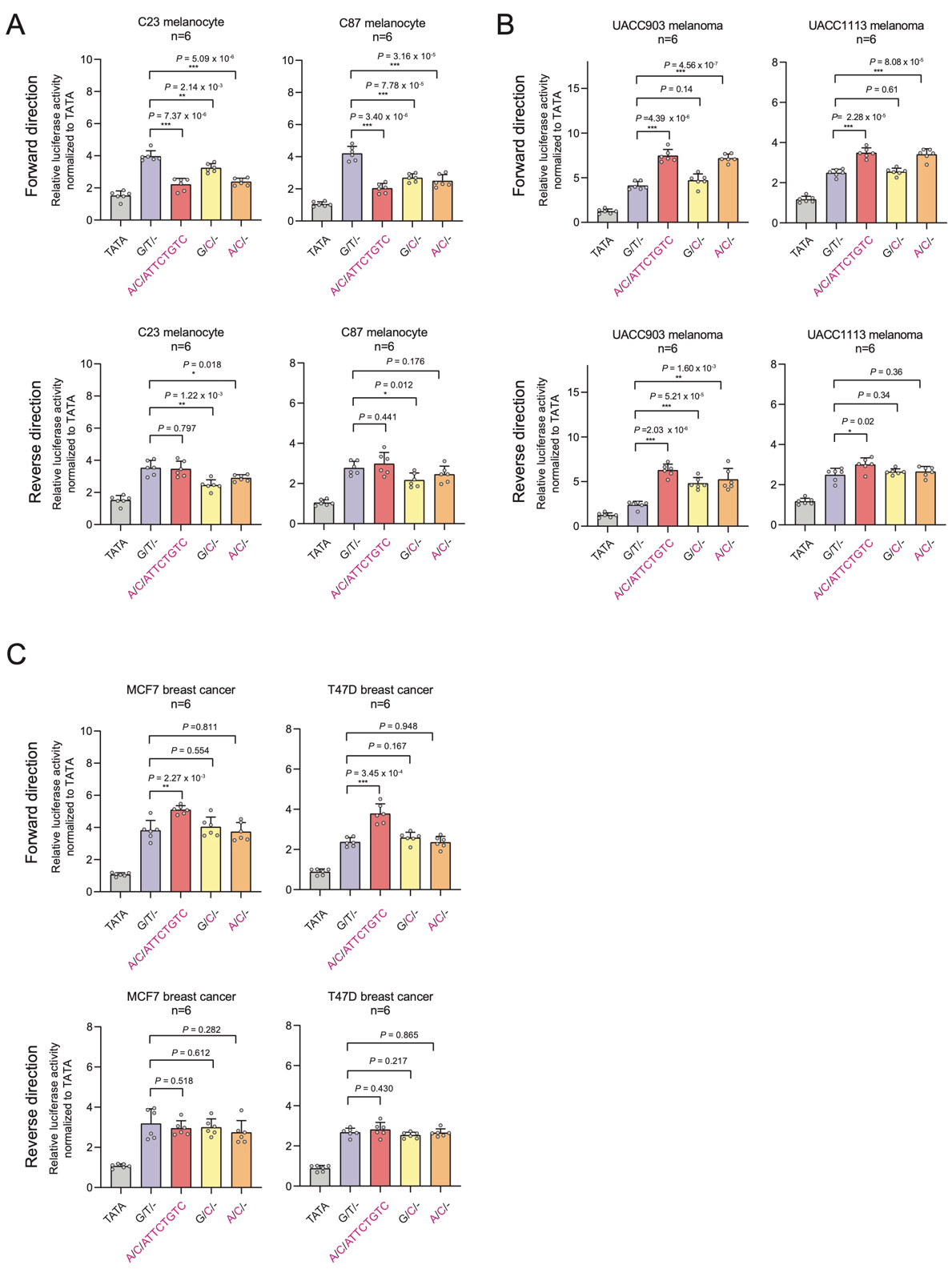

**Supplementary Figure S17. Haplotype-based luciferase assays.** Since rs3769823, rs3769821, and rs59308963 are 484 bp apart, in LD with each other, and in a continuous regulatory region, they were tested as a haplotype using a 641bp construct to assay the most commonly observed haplotypes. Sequences encompassing rs3769823, rs3769821, and rs59308963 (641 bp) were cloned 5’ of pGL4.23 minimal TATA promoter and transfected into (A) primary melanocytes C23 and C87, (B) melanoma cell lines UACC903 and UACC1113, and (C) breast cancer cell lines MCF7 and T47D. Individual luciferase reporter activity was measured 24 h after transfection and was normalized against *Renilla* luciferase activity. Individual *P*-values are shown relative to the all-risk allele haplotype (two-tailed, unpaired t-test assuming unequal variances). Data and *P*-values are from one representative set (*n* = 6) from three biological replicates. TATA, minimal promoter control; risk allele marked in red (rs3769823/rs3769821/rs59308963).

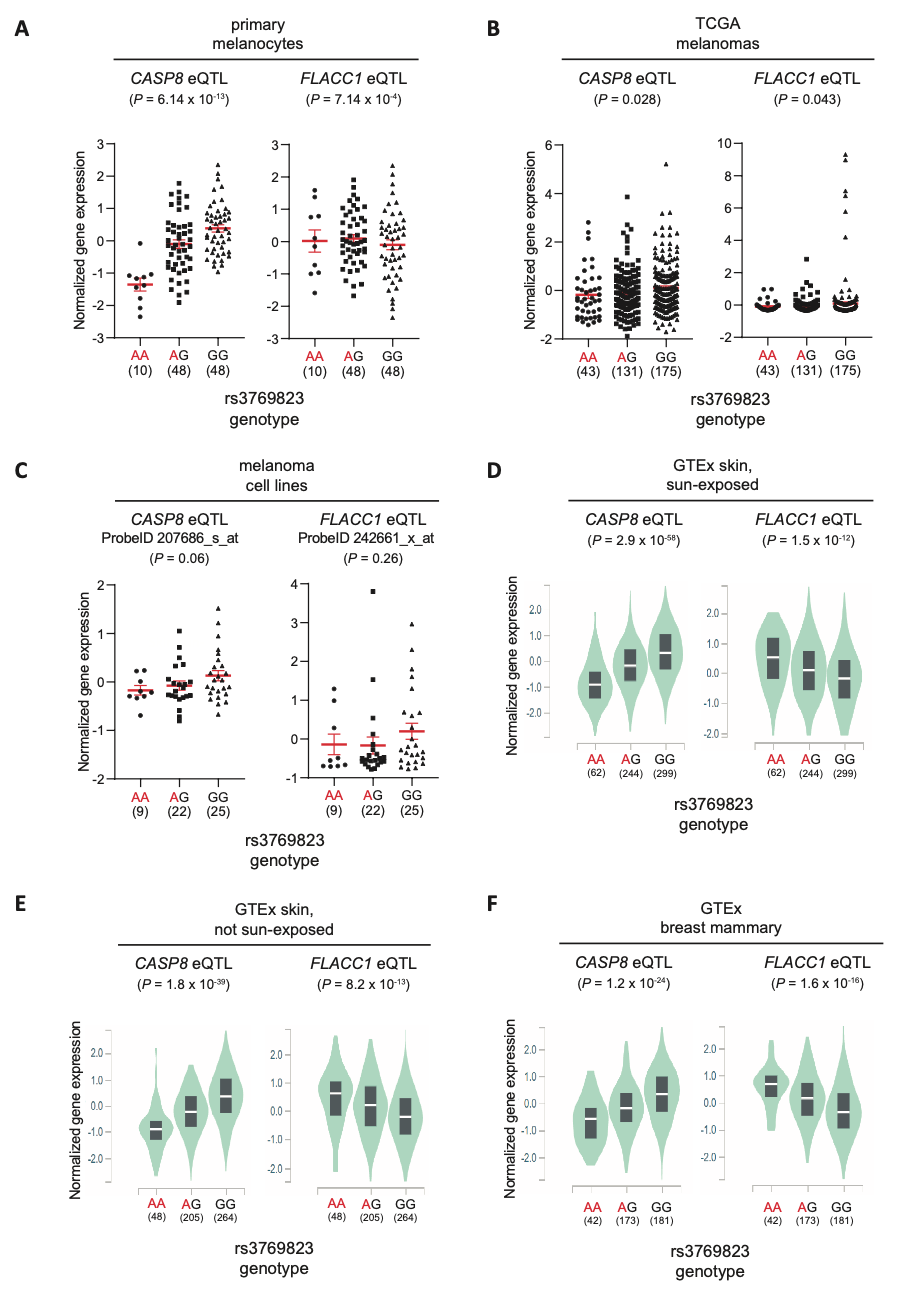

**Supplementary Figure S18. The melanoma risk-associated A allele of rs3769823 is correlated with lower *CASP8* expression and higher *FLACC1*.** eQTL plots of *CASP8* levels are shown for (A) human primary melanocytes cultures (*n* = 106), (B) TCGA melanoma tumors (*n*=349), (C) a panel of early-passage melanoma cell lines (*n* = 59), (D) GTEx sun-exposed skin tissues (*n* = 605), (E) GTEx not sun-exposed skin tissues (*n* = 517), and (F) GTEx breast mammary tissues (*n* = 396) in relation to rs3769823 genotype. *P*-values were derived from linear regression. The risk-A allele of rs3769823 is labeled in red.

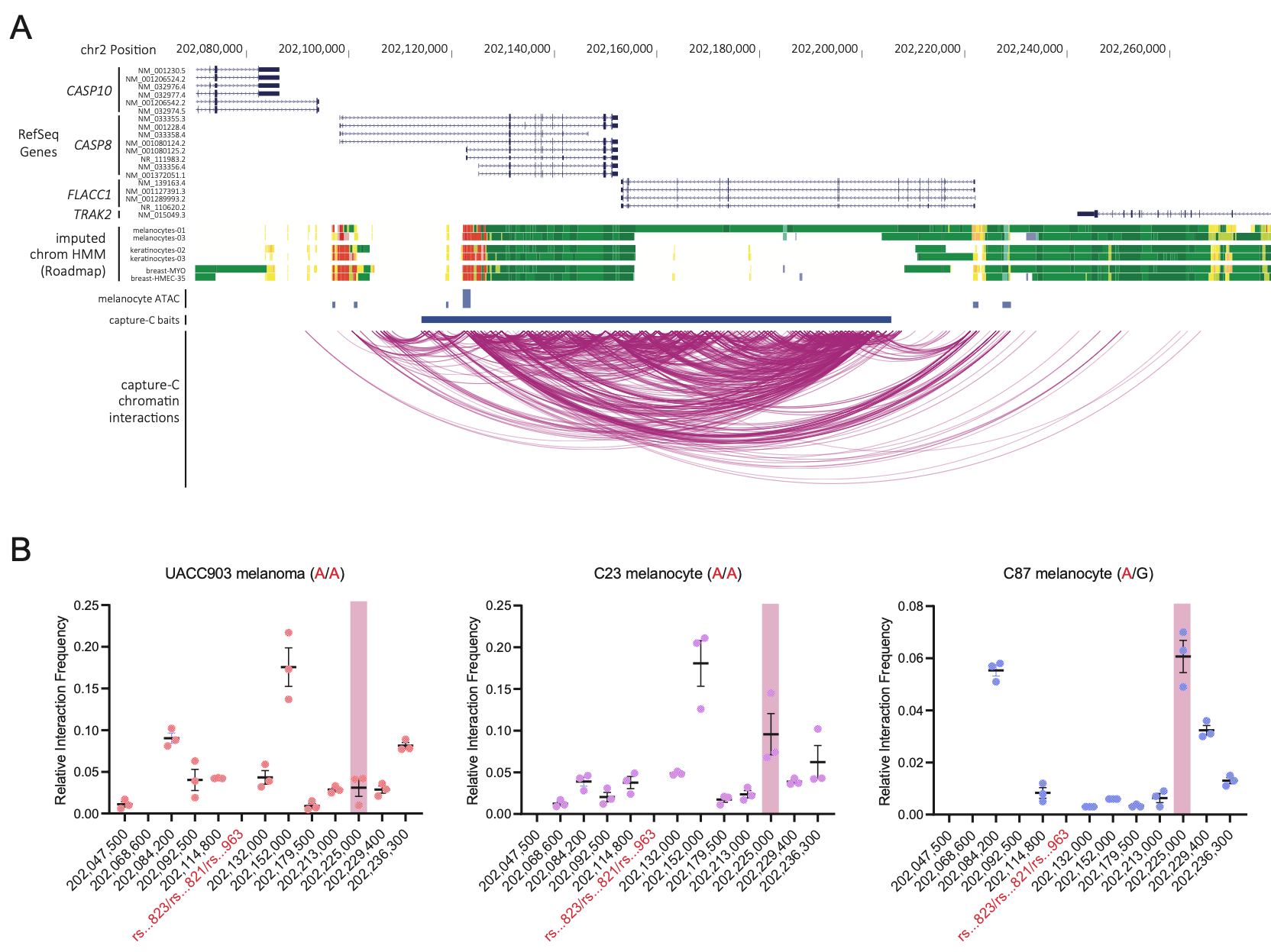

**Supplementary Figure S19. Region-specific Capture-C and chromatin conformation capture (3C) show a physical interaction between the region harboring rs3769823, rs3769821, and rs59308963 and the *FLACC1* promoter.** (A) Significant chromatin interactions identified by Capture-C across the entire region of association. Genomic locations for which capture baits were designed are noted, and significant interactions are shown as purple arcs. RefSeq genes, imputed ChromHMM and DNaseI hypersensitivity (DHS) data for two melanocyte cultures generated by the Roadmap Epigenomic Project are shown. Loops were called using data from five genetically unrelated melanocyte cultures (three biological replicates per culture) were analyzed together to call loops. (B) Chromatin conformation capture (3C) analysis of local chromatin interactions with the region harboring rs3769823, rs3769821, and rs59308963 (rs823/rs821/rs963; labeled in red). A restriction fragment harboring rs3769823, rs3769821, and rs59308963 was used as bait and physical interactions between this location and various regions across chromosome 2q33.1 were assessed by quantitative PCR in 3C libraries generated from UACC903 melanoma cells as well as two independent human primary melanocyte cultures (C23 and C87). For each experiment, PCR amplification for each target primer from 3C libraries was first normalized to the PCR amplification of the target primer from BAC library DNA and subsequently normalized to the PCR amplification of the target primer rs821/rs821/rs963. One representative experiment from three biological replicates is shown. Mean with SEM are plotted along with individual data values. The location corresponding to the *FLACC1* promoter is highlighted in pink.

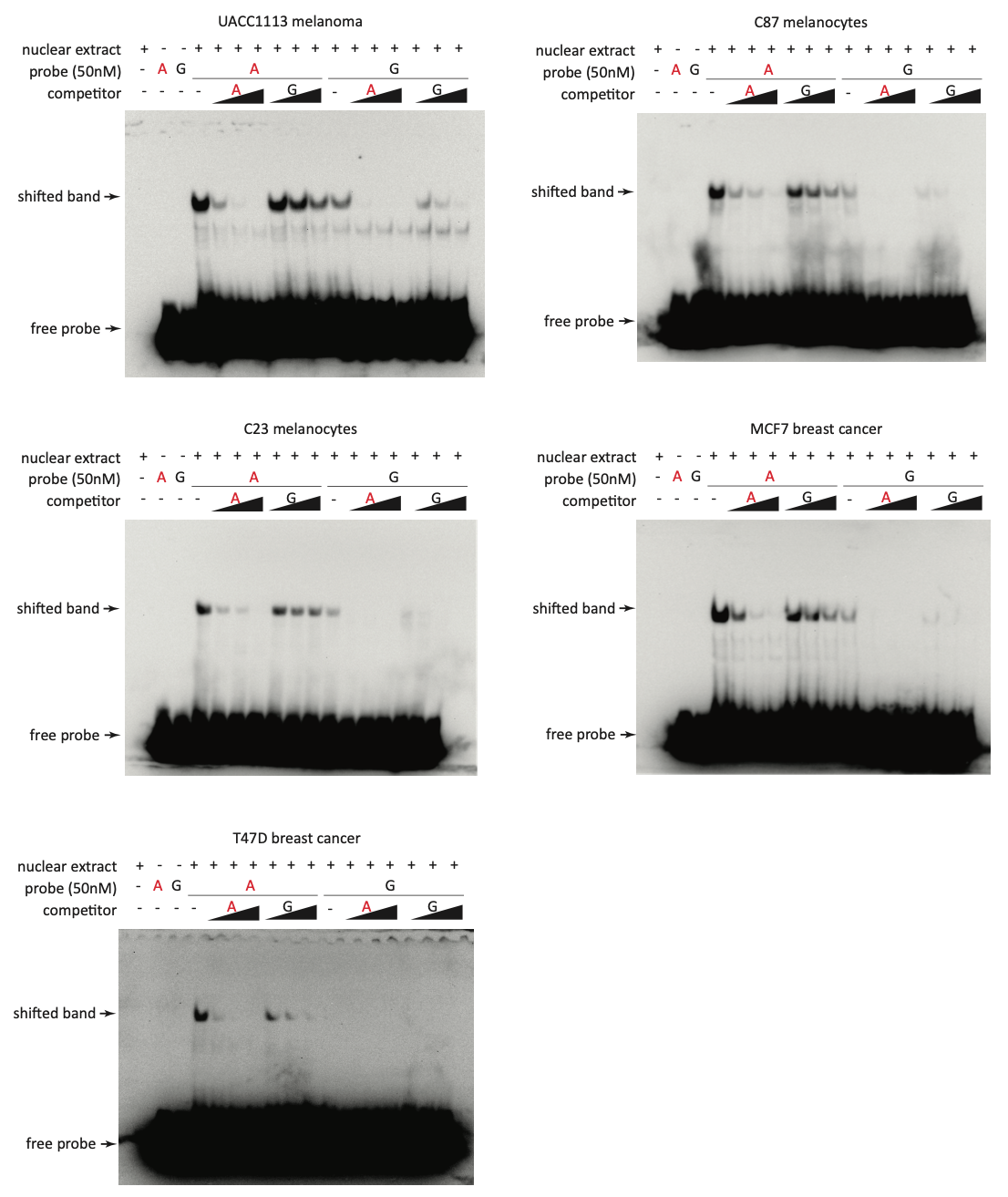

**Supplementary Figure S20. The melanoma-associated variant rs3769823 shows allelic-specific protein binding in multiple cell lines.** EMSAs were performed using biotin-labeled double-stranded oligonucleotides for the A-risk and G-protective allele of rs3769823 and nuclear extract from melanoma cell line UACC1113, two primary melanocyte cultures (C87, C23), and two breast cancer cell lines (MCF7 and T47D). 50 nM of 21 bp probes for the A-risk and G-protective alleles were used. 2x, 5x, or 10x molar excess of unlabeled competitor was added in specified lanes.

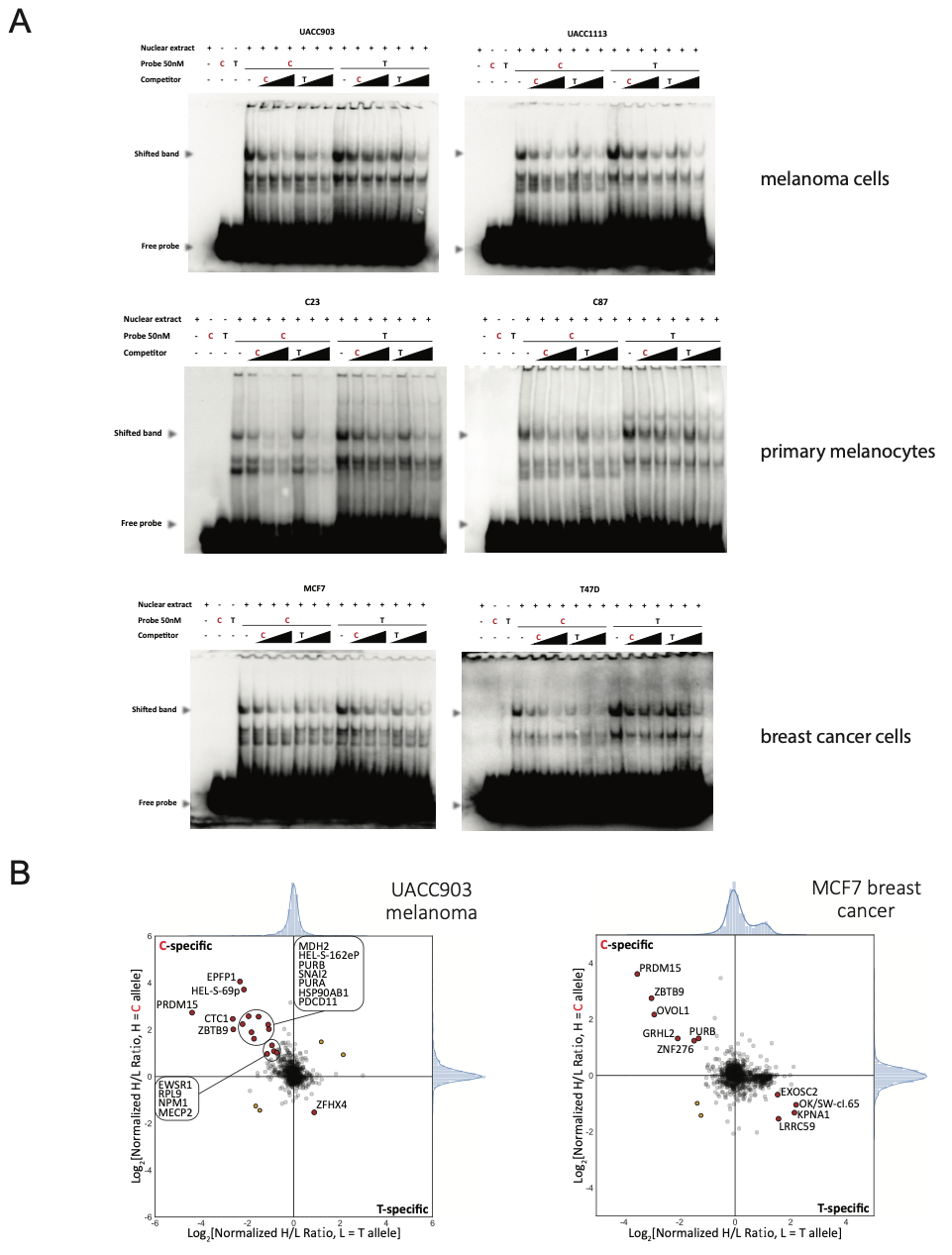

**Supplementary Figure S21. EMSAs and quantitative mass-spectrometry for the functional candidate variant rs3769821 in multiple cell lines.** (A) EMSAs were performed using biotin-labeled double-stranded DNA probes encompassing 10 bp to either side of rs3769821 with two different alleles. Nuclear extracts from melanoma cell lines (UACC903 and UACC1113), primary melanocytes (C23 and C87), or breast cancer cell lines (MCF7 and T47D) were used. 2x, 5x, or 10x molar excess of unlabeled competitor was added in specified lanes for competition. C is risk allele in red and T is protective allele. (B) Allele-specific binding proteins were identified by mass spectrometry using nuclear extracts of UACC903 melanoma cells (left) and MCF7 breast cancer cells (right) and 21 bp double-stranded DNA probes for C-risk (red) and T-protective alleles of rs3769821. The dimethyl-labeling ratios of proteins bound to C/T probes are plotted on the *x* and *y* axes for label-swapping experiments. Red circles highlight proteins enriched above the background in both experiments.

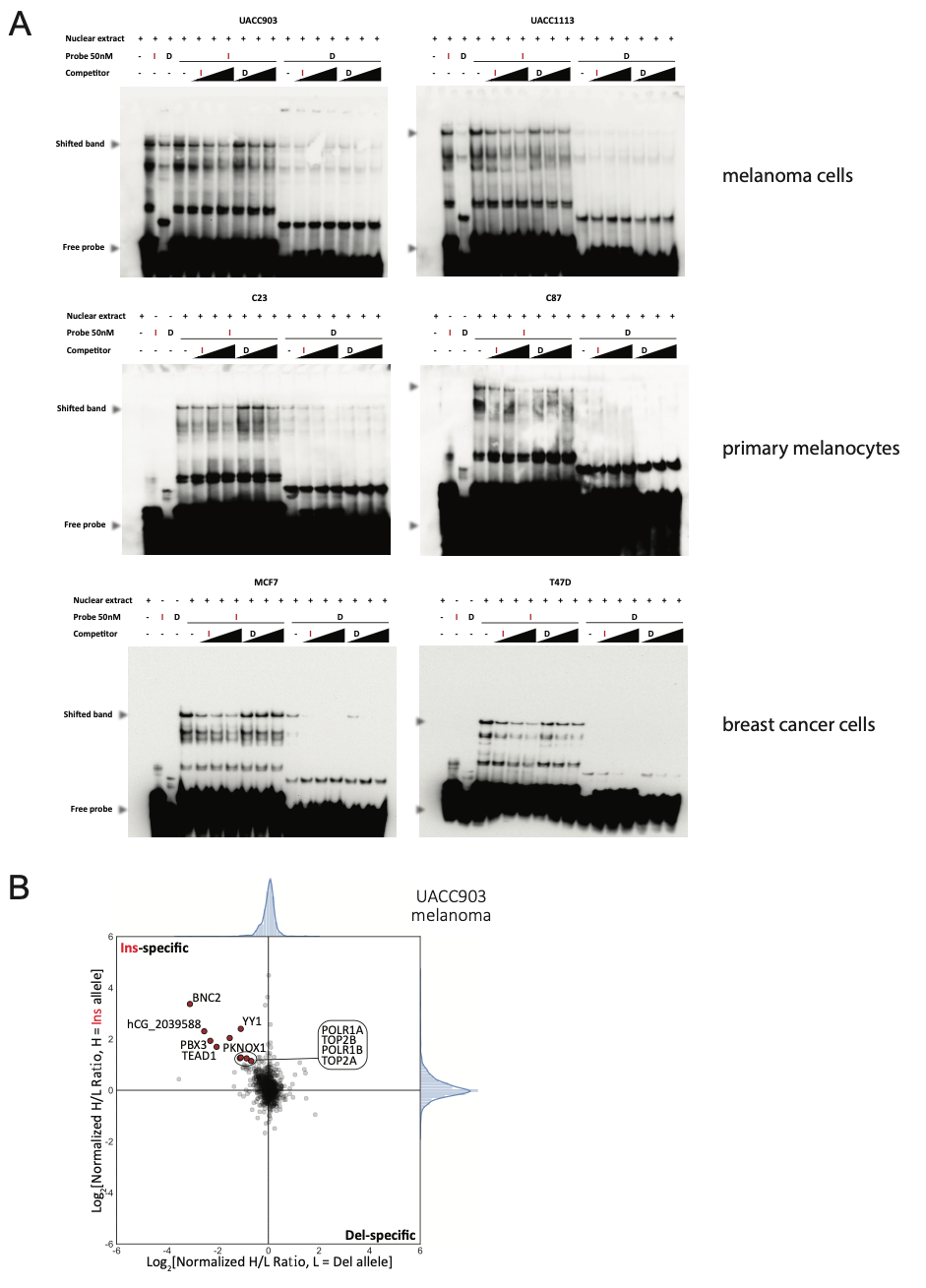

**Supplementary Figure S22. EMSAs and quantitative mass-spectrometry for the functional candidate variant rs59308963 in multiple cell lines.** (A) EMSAs were performed using biotin-labeled double-stranded DNA probes encompassing 10 bp to either side of rs59308963 with two different alleles. Nuclear extracts from melanoma cell lines (UACC903 and UACC1113), primary melanocytes (C23 and C87), or breast cancer cell lines (MCF7 and T47D) were used. 2x, 5x, or 10x molar excess of unlabeled competitor was added in specified lanes for competition. The insertion allele (I) is the risk allele while the deletion allele (D)is the protective allele. (B) Allele-specific binding proteins were identified by mass spectrometry using nuclear extracts of melanoma cell line UACC903 and 21 bp double-stranded DNA probes with insertion-risk (Ins) and deletion-protective (Del) alleles of rs59308963. The dimethyl-labeling ratios of proteins bound to Ins/Del probes are plotted on the *x* and *y* axes for label-swapping experiments. Red circles highlight proteins enriched above the background in both experiments.

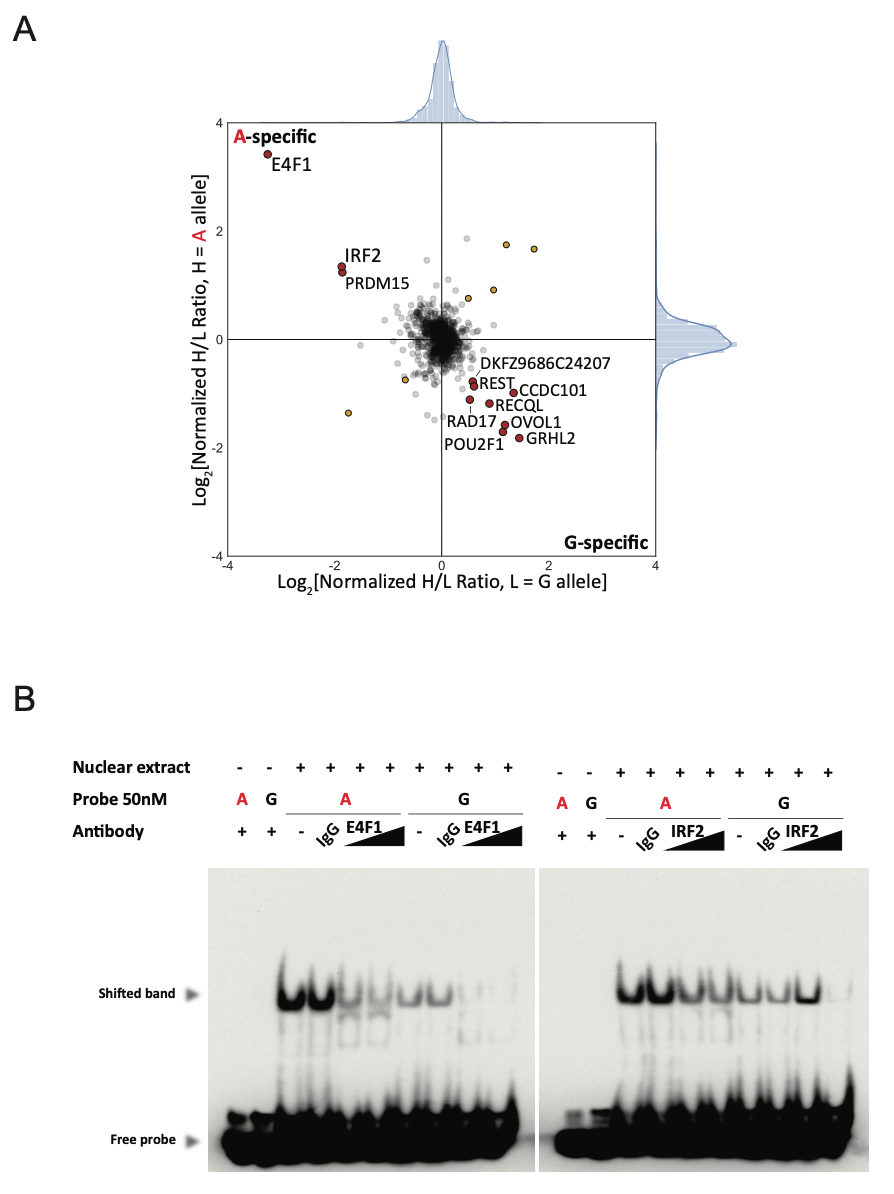

**Supplementary Figure S23. E4F1 and IRF2 bind to A-risk allele of rs3769823 in MCF7 breast cancer cells.** (A) Quantitative mass-spectrometry was performed to identify allele-specific rs3769823 binding proteins using nuclear extract of MCF7 and 21 bp double-stranded DNA probes with A-risk or G-protective alleles, respectively. A-allele specific interacting proteins are shown in the top left quadrant and G-allele specific interactors in the top left quadrant. Using label-swapping of high(H)-mass or low(L)-mass label, the A-bound/G-bound ratio is shown *y*-axis, and the G-bound/A-bound ratio on the *x*-axis for replicate label-swapping experiments. Red circles highlight proteins enriched above the background in both experiments. (B) EMSA and super-shifts assays using anti-E4F1 or anti-IRF2 antibody are shown for MCF7 breast cancer cells, where the A-risk specific shifted band (arrows) is diminished. A representative set from three independent experiments is shown.

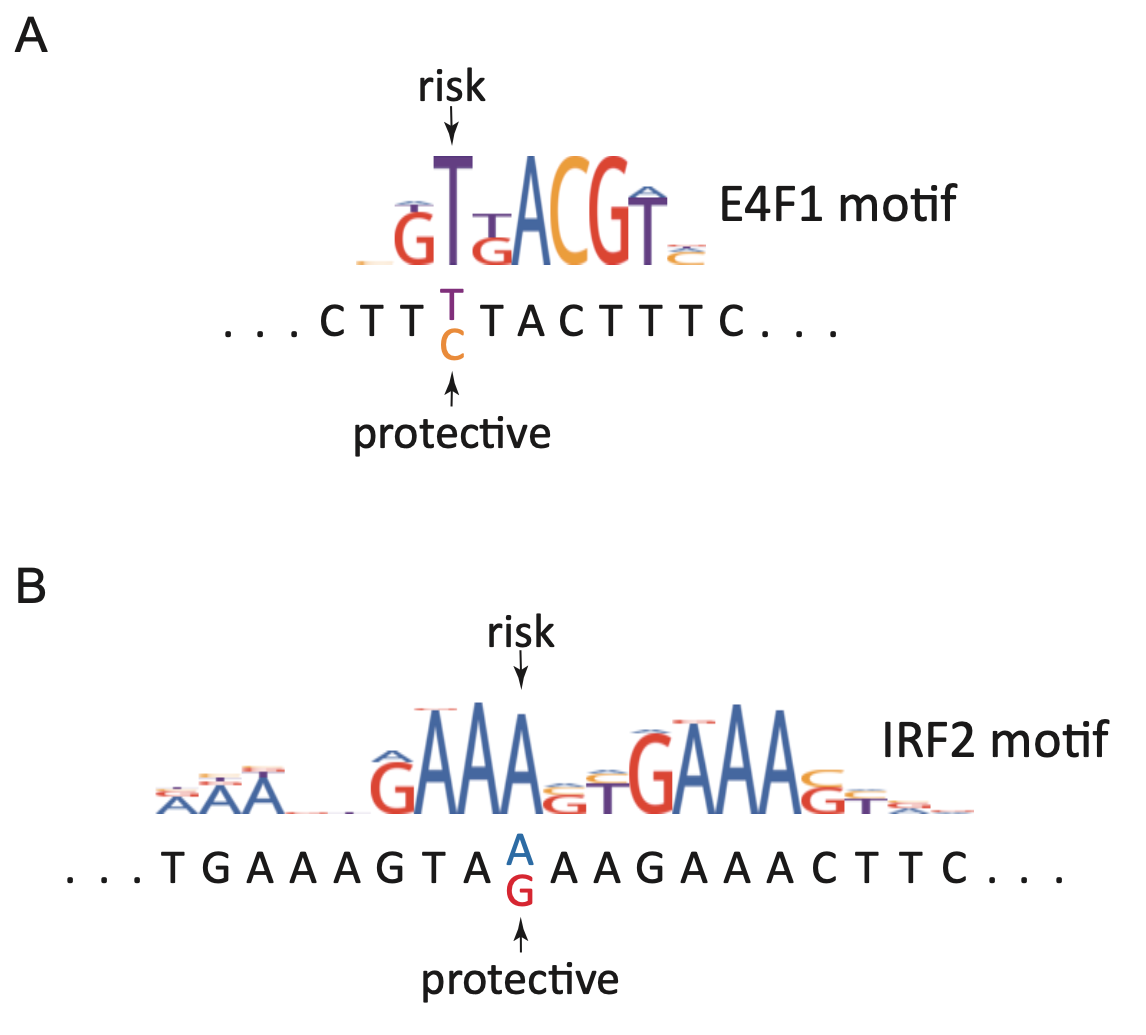

**Supplementary Figure S24. E4F1 and IRF2 transcription factor binding motifs relative to the risk- and protective-associated alleles of rs3769823.** (A) E4F1 and (B) IRF2 binding motifs are shown as position weight matrices (motifs obtained from HOCOMOCO database and plotted using weblogo3) are shown above the genomic sequence surrounding rs3769823, with the risk-associated allele matching consensus E4F1 and IRF2 binding motifs.

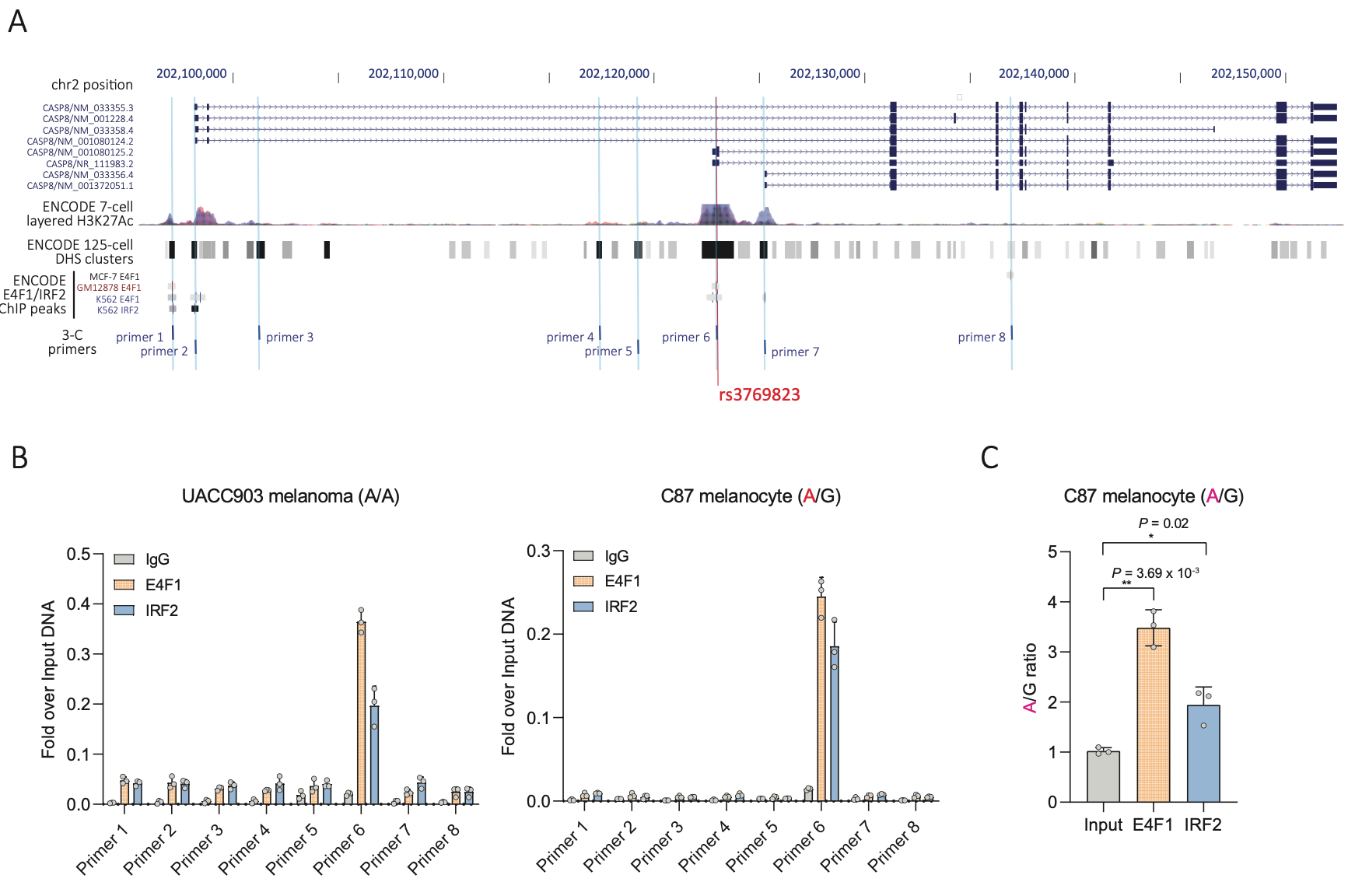

**Supplementary Figure S25. E4F1 and IRF2 bind to the risk associated A allele of rs3769823.** (A) The genomic positions of the amplicons for eight qPCR primer sets are shown relative to rs3769823 (red line) in the alternative first exon of *CASP8*. Primer sets 1, 2, 6, 7, and 8 were selected based on prior evidence of E4F1 or IRF2 in ChIP-seq data from ENCODE. Primer sets 3 and 5 are designed to assay open chromatin and enhancer regions annotated in ChromHMM data from two melanocyte cultures assayed by the RoadMap Epigenome Project. Primer set 4 was designed as a negative control over inaccessible chromatin with no evidence of E4F1/IRF2 binding, while primer set 6 was designed to specifically assay binding over rs3769823. (B) ChIP-qPCR data are shown using anti-E4F1 antibody, anti-IRF2 antibody or normal IgG for both UACC903 melanoma cells C87 primary melanocytes. Genotypes for rs3769823 for both cultures are indicated in the label. Relative quantities are shown as fold over input DNA. Means of PCR triplicates with SD are plotted. One representative set of three biological replicates for each cell line is shown. (C) Taqman genotyping data for rs3769823 with E4F1 or IRF2 ChIP DNA from C87 primary melanocytes (heterozygous for rs3769823) are shown normalized to input DNA. qPCR triplicates were analyzed. A representative plot and *P*-values from a single experiment from three biological replicates is shown. Individual *P* values are show against Input (two-tailed, unpaired t-test assuming unequal variances).

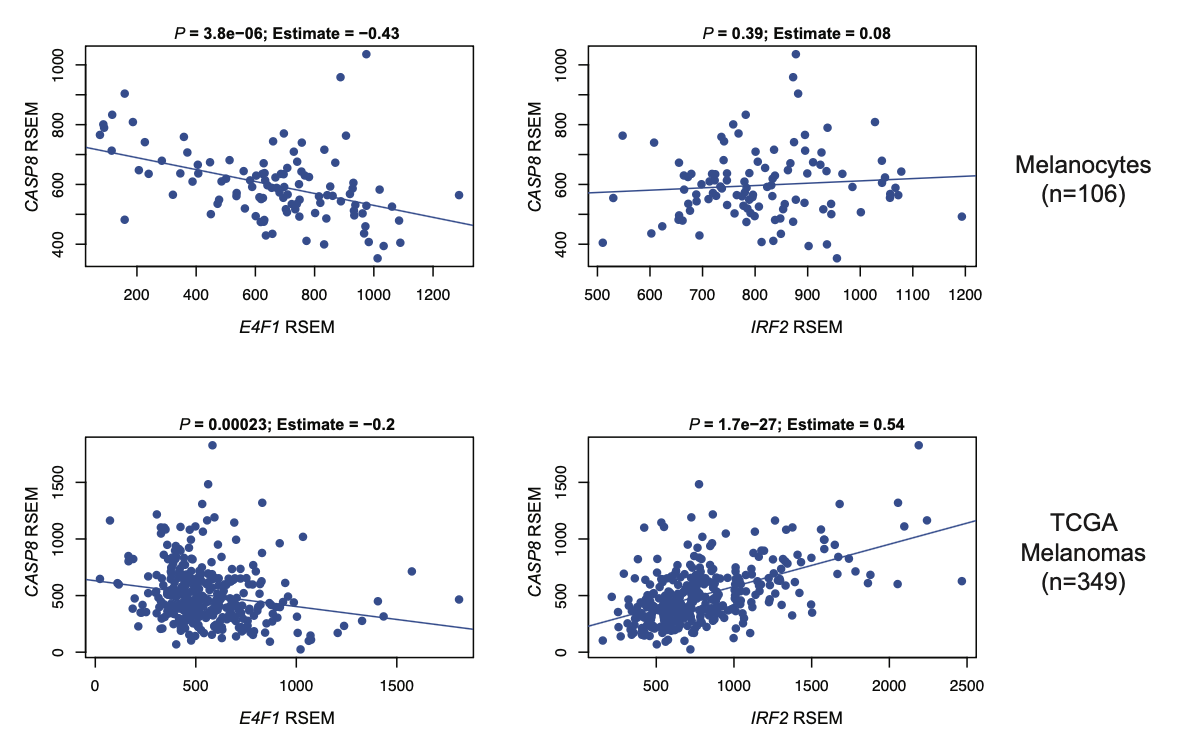

**Supplementary Figure S26. Correlation of CASP8 with E4F1 and IRF2 in primary melanocytes and melanomas.** Pearson correlation coefficient (Estimate) and *P*-value (*P*) between *CASP8* and *E4F1* or *IRF2* gene expression is shown for (top) 106 human primary melanocytes and (bottom) 349 TCGA melanoma tumors. Measurment of expression was using RNA Sequencing by Expectation Maximization (RSEM).

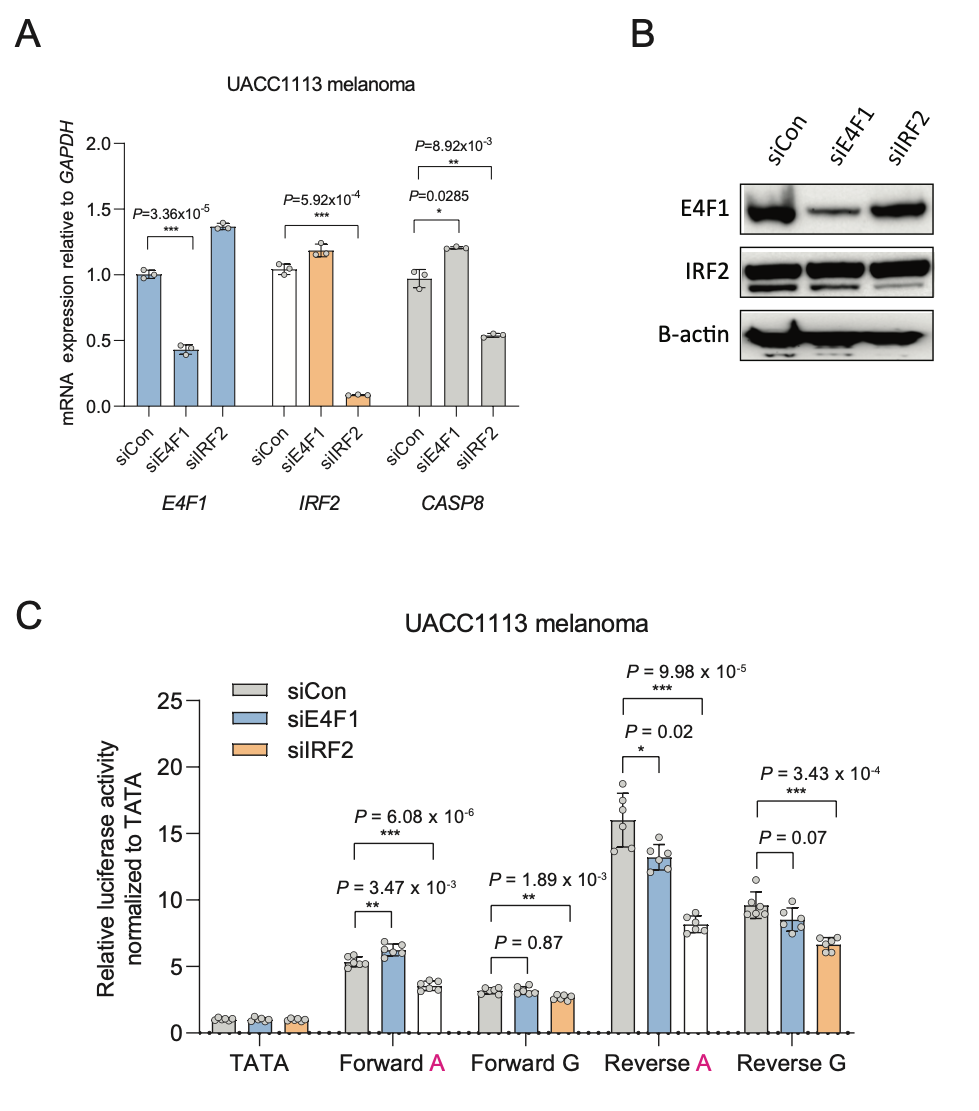

**Supplementary Figure S27. E4F1 or IRF2 and rs3769823 regulate *CASP8* expression in UACC1113 melanoma cells.** (A) *E4F1* or *IRF2* were knocked down using pools of four siRNAs each in UACC1113 melanoma cells, and *E4F1*, *IRF2* and *CASP8* levels were measured. *GAPDH*-normalized *E4F1*, *IRF2,* or *CASP8* mRNA levels are shown as fold-change over those from non-targeting siRNA. A representative experiment from three biological replicates is shown (individual datapoints, mean, and SEM are plotted). *P*-values are shown against control siRNA from one representative set. (B) Western blotting was performed using anti-E4F1, anti-IRF2, or anti-GAPDH antibodies and cell lysates of UACC1113 transfected with siRNAs targeting *E4F1* or *IRF2*. GAPDH were used as loading control. A representative experiment from three biological replicates is shown. (C) Individual luciferase assays were performed by co-transfecting rs3769823 constructs with *E4F1*, *IRF2,* or control siRNAs into UACC1113 melanoma cells. *Renilla*-normalized relative luciferase activities are measured relative to an empty vector control harboring only a minimal promoter (TATA). Data and *P*-values are shown for one representative experiment (*n* = 6 technical replicates) from three biological replicates. A two-tailed t-test assuming unequal variance was used to calculate all *P*-values shown against control siRNA.

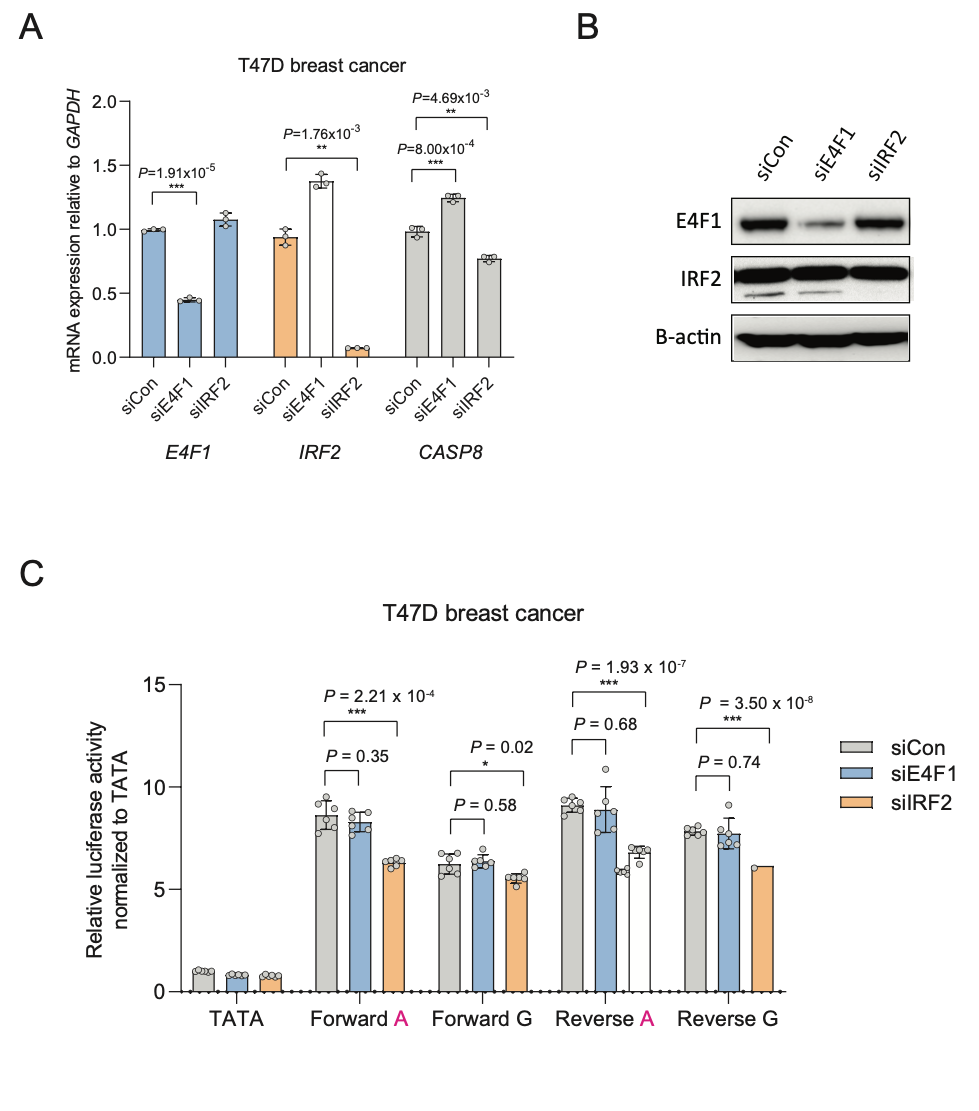

**Supplementary Figure S28. E4F1 or IRF2 and rs3769823 regulate *CASP8* expression in T47D breast cancer cells.** (A) *E4F1* or *IRF2* were knocked down using pools of four siRNAs each in T47D breast cancer cells, and *E4F1*, *IRF2* and *CASP8* levels were measured. *GAPDH*-normalized *E4F1*, *IRF2,* or *CASP8* mRNA levels are shown as fold-change over those from non-targeting siRNA. A representative experiment from three biological replicates is shown (individual datapoints, mean, and SEM are plotted). *P*-values are shown against control siRNA from one representative set. (B) Western blotting was performed using anti-E4F1, anti-IRF2, or anti-GAPDH antibodies and cell lysates of T47D transfected with siRNAs targeting *E4F1* or *IRF2*. GAPDH were used as loading control. A representative experiment from three biological replicates is shown. (C) Individual luciferase assays were performed by co-transfecting rs3769823 constructs with *E4F1*, *IRF2,* or control siRNAs into T47D breast cancer cells. *Renilla*-normalized relative luciferase activities are measured relative to an empty vector control harboring only a minimal promoter (TATA). Data and *P*-values are shown for one representative experiment (*n* = 6 technical replicates) from three biological replicates. A two-tailed t-test assuming unequal variance was used to calculate all *P*-values shown against control siRNA.

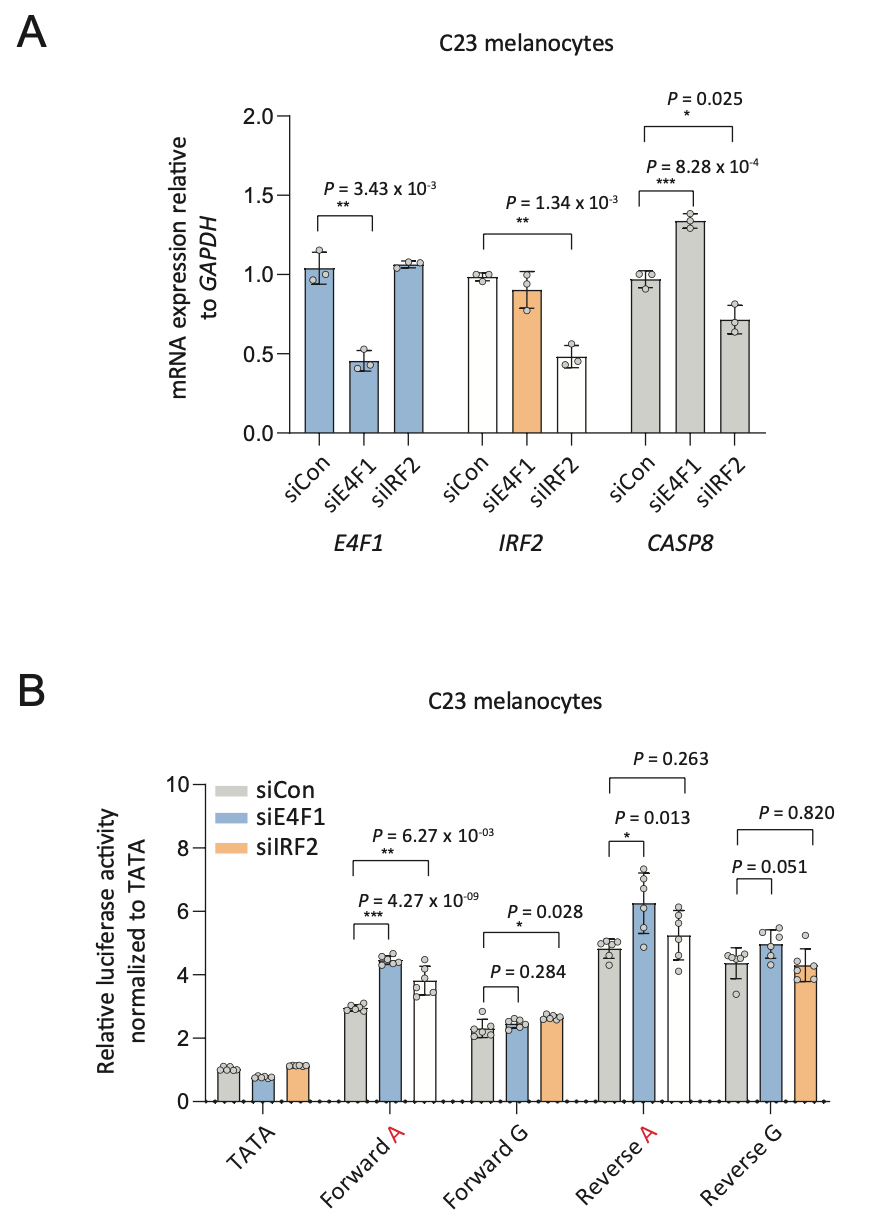

**Supplementary Figure S29. E4F1 or IRF2 and rs3769823 regulate *CASP8* expression in C23 primary melanocyte cultures.** (A) *E4F1* or *IRF2* were knocked down using pools of four siRNAs each in C23 human primary melanocyte cultures, and *E4F1*, *IRF2* and *CASP8* levels were measured. *GAPDH*-normalized *E4F1*, *IRF2* or *CASP8* mRNA levels are shown as fold-change over those from non-targeting siRNA. One representative experiment from three biological replicates is shown (individual datapoints, mean, and SEM are plotted). *P*-values are presented against non-targeting siRNA from one representative set. (B) Individual luciferase assays were performed by co-transfecting rs3769823 reporter constructs together with *E4F1*, *IRF2* or control siRNAs into C23 primary melanocyte cultures. *Renilla*-normalized relative luciferase activities are measured relative to an empty vector control harboring only a minimal promoter (TATA). Data and *P*-values are shown for one representative experiment (*n*=6 technical replicates) from three biological replicates. A two-tailed t-test assuming unequal variance was used to calculate all *P-*values shown against control siRNA.
